# The Dmc1 recombinase physically interacts with and promotes the meiotic crossover functions of the Mlh1-Mlh3 endonuclease

**DOI:** 10.1101/2023.11.13.566911

**Authors:** Gianno Pannafino, Jun Jie Chen, Viraj Mithani, Lisette Payero, Michael Gioia, J. Brooks Crickard, Eric Alani

## Abstract

The accurate segregation of homologous chromosomes during the Meiosis I reductional division in most sexually reproducing eukaryotes requires crossing over between homologs. In baker’s yeast approximately 80 percent of meiotic crossovers result from Mlh1-Mlh3 and Exo1 acting to resolve double-Holliday junction (dHJ) intermediates in a biased manner. Little is known about how Mlh1-Mlh3 is recruited to recombination intermediates and whether it interacts with other meiotic factors prior to its role in crossover resolution. We performed a haploinsufficiency screen in baker’s yeast to identify novel genetic interactors with Mlh1-Mlh3 using sensitized *mlh3* alleles that disrupt the stability of the Mlh1-Mlh3 complex and confer defects in mismatch repair but do not disrupt meiotic crossing over. We identified several genetic interactions between *MLH3* and *DMC1,* the recombinase responsible for recombination between homologous chromosomes during meiosis. We then showed that Mlh3 physically interacts with Dmc1 *in vitro* and at times in meiotic prophase when Dmc1 acts as a recombinase. Interestingly, restricting *MLH3* expression to roughly the time of crossover resolution resulted in a *mlh3* null-like phenotype for crossing over. Our data are consistent with a model in which Dmc1 nucleates a polymer of Mlh1-Mlh3 to promote crossing over.

## Introduction

Accurate segregation of homologous chromosomes during the reductional division (Meiosis I) in most sexually reproducing eukaryotes is accomplished by the physical tethering of homologs through crossing over and distal chromatid cohesion, enabling stable orientation of homologs on the meiotic spindle (Maguire 1974; Hunter 2015; Zickler and Kleckner 2015). Defects in crossing over result in gametes that are aneuploid and often non-viable. In humans such abnormalities have been linked to miscarriages, birth defects and developmental disabilities (Hassold et al. 2001; Nagaoka et al. 2010; Hunter 2015).

Meiotic recombination events in yeast and mammals are initiated in meiotic prophase through Spo11-induced double strand breaks (DSBs) that occur genome-wide (Keeney et al. 1997; Pan et al. 2011). These DSBs are resected in a 5’ to 3’ direction by the activities of factors that include Mre11-Rad50-Xrs2 (MRX; MRE11-RAD50-NBS1 in humans) and Exo1 (Cao et al. 1990; Fiorentini et al. 1997; Nicolette et al. 2010; Zakharyevich et al. 2010). The RecA homologs Rad51 and Dmc1 are loaded onto the resulting resected 3’ tails in a defined orientation, with Dmc1 located nearer to the 3’ end. Dmc1 then catalyzes a homology search for allelic loci, with Rad51 acting as an essential accessory factor in a non-catalytic role. Dmc1-coated 3’ tails invade a homologous donor locus, forming a displacement loop (D-loop) that can be extended by DNA polymerase activity (Bishop et al. 1994; Cloud et al. 2012; Brown et al. 2015; Hinch et al. 2020; Hunter 2015).

In baker’s yeast, approximately half of the 150-200 Spo11-induced meiotic DSBs are repaired as non-crossovers through the helicase-driven unwinding of extended D-loops (Synthesis-Dependent Strand Annealing (SDSA), reviewed in Pyatnitskaya et al. 2019). The remainder are stabilized by the functionally diverse ZMM family of pro-crossover factors consisting of Zip1-4, Spo16, Msh4-Msh5, and Mer3 to form single-end invasion intermediates (SEI) that commit recombination to the Class I crossover pathway. These events are biased towards the use of a homologous chromosome as a donor for repair (Fig. 1; Schwacha and Kleckner 1997; reviewed in Brown and Bishop 2014 and Hunter 2015). The SEI forms at roughly the same time as the synaptonemal complex, a structure that forms between homologs and is thought to aid in the removal of chromosomal interlocks during the homology search. (Hunter and Kleckner 2001; Snowden et al. 2004; Fung et al. 2004; Börner et al. 2004; Lynn et al. 2007; Storlazzi et al. 2010; De Muyt et al. 2018; Pyatnitskaya et al. 2019). Following DNA synthesis steps and capture of the second end of the broken homolog, SEIs mature into symmetric double Holliday junctions (dHJs) that are resolved asymmetrically to form crossovers through the actions of the mismatch repair (MMR) family member Mlh1-Mlh3 and the exo/endonuclease Exo1 to yield crossover products (Allers and Lichten 2001; Zakharyevich et al. 2012; reviewed in Pyatnitskaya et al. 2019). Crossovers dependent on Mlh1-Mlh3 bear the hallmarks of the Class I pathway; they are widely and evenly spaced (crossover interference) and display crossover assurance, a mechanism which ensures that each homolog pair receives at least one crossover (Szostak et al. 1983; Sym et al. 1993; Sym et al. 1994; Schwacha and Kleckner 1994; Schwacha and Kleckner 1995; Novak et al. 2001; Shinohara et al. 2003; Börner et al. 2004; Hillers 2004; Martini et al. 2006; Jones and Franklin 2006; Mancera et al. 2008; Shinohara et al. 2008; Zakharyevich et al. 2012; Chakraborty et al. 2017).

**Fig. 1.**
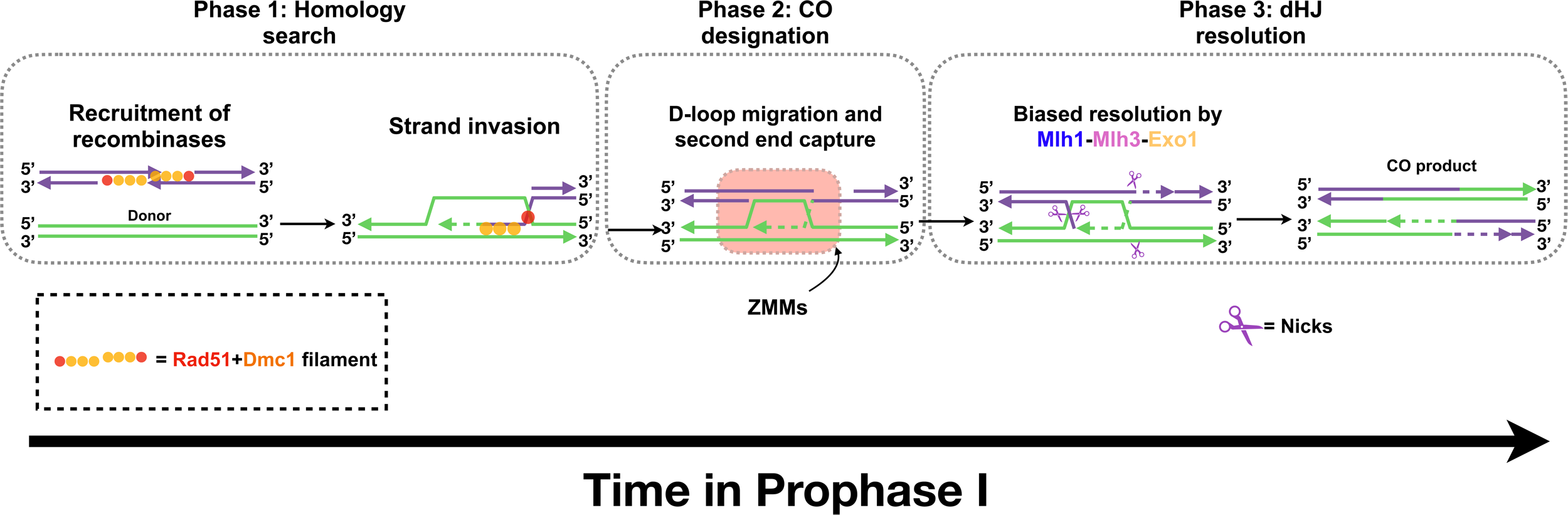
Model of canonical resolvase role for Mlh1-Mlh3 in meiotic recombination. A model for the Type I CO pathway in yeast meiotic prophase. Phase I, homology search: Following Spo11-mediated DSB formation and 5’ to 3’ resection of DSBs, a recombinase filament (Dmc1 filament initiates at the end of the Rad51 filament) forms at the 3’ resected ends. This filament initiates a search for the allelic locus on the homologous donor chromosome. Phase 2, CO designation: ZMM-facilitated stabilization of SEIs, D-loop migration, and second-end capture commit recombination intermediates to the Type I crossover pathway. Phase 3, dHJ resolution: Mlh1-Mlh3 and Exo1 participate in biased resolution of dHJs to produce crossover products.

How might Mlh1-Mlh3 act in crossover resolution? Mlh3 contains a highly conserved (DQHA(X)2E(X)4E) endonuclease motif within its C-terminus (Kadyrov et al. 2006; Nishant et al. 2008). Disruption of this motif (*mlh3-D523N*) abolishes the endonuclease activity of Mlh1-Mlh3 *in vitro* and confers *mlh3* null-like defects in crossing over (Nishant et al. 2008; Rogacheva et al. 2014; Ranjha et al. 2014). Curiously, Mlh1-Mlh3 does not display the biochemical activities characteristic of an intrinsic HJ resolvase. It is capable of binding model HJ substrates, but such binding inhibits its endonuclease activity. Rather, Mlh1-Mlh3 endonuclease activity is activated *in vitro* by polymer formation on large DNA substrates and *mlh3* mutations predicted to disrupt polymer formation confer crossover defects (Hall et al. 2001; Manhart et al. 2017; Dai et al. 2021). Mlh1-Mlh3 and Exo1 form a constitutive complex throughout meiosis (Sanchez et al. 2020) and recent *in vitro* studies showed that Mlh1-Mlh3 endonuclease activity is stimulated by Msh4-Msh5, Exo1, and the DNA polymerase processivity factor PCNA (Cannavo et al. 2020; Kulkarni et al. 2020). While we do not have a clear understanding of how Mlh1-Mlh3 endonuclease activity is directed to form crossovers, several dHJ resolution models have been presented that incorporate the above Mlh1-Mlh3 activities and a role for Exo1 in protecting DNA nicks (Manhart et al. 2017; Marsolier-Kergoat et al. 2018; Kulkarni et al. 2020; Gioia et al. 2023).

Mlh1-Mlh3 is thought to act late in prophase I during dHJ resolution. Interestingly, Sanchez et al. (2020) reported in budding yeast that Mlh3 localizes to DSB hotspots early in meiotic prophase, showing kinetics similar to the ZMM protein Zip3 (Serrentino et al. 2013). In mice, MLH3 forms foci in early pachynema prior to MLH1-MLH3 foci formation (Kolas et al. 2005). Other studies have suggested that Mlh3 promotes the early stages of meiotic recombination. For example, Al-Sweel et al. (2017) observed that the median gene conversion tract length associated with crossovers was longer in *mlh3*Δ compared to *wild-type*. Marsolier-Kergoat et al. (2018) inferred through an analysis of genome-wide heteroduplex DNA patterns that Mlh1-Mlh3 acts to limit inward branch migration (see Peterson et al. 2020; Ahuja et al. 2021; Martini et al. 2011). Recently, it was shown by Premkumar et al. (2023) that mouse MLH3 plays an early structural role in differentiating SEIs further towards Class I resolution. These and other studies are suggestive of an early role for Mlh3 but how it could act in early stages of recombination remains unclear.

To better understand the role of Mlh1-Mlh3 in meiotic prophase, we performed a haploinsufficiency meiotic screen aimed at identifying factors that genetically interact with *MLH3* in its procrossover role. Through this approach we obtained evidence that Mlh3 interacts with the recombinase Dmc1. We also showed that expression of Mlh3 in early meiotic prophase was critical for its meiotic crossover function, consistent with a role for Mlh3 in meiotic crossing over that precedes crossover resolution. These observations provide explanations for *mlh3*Δ phenotypes associated with early meiotic roles and led us to a model in which the Dmc1 recombinase filament provides a platform for Mlh1-Mlh3 polymer formation and subsequent endonuclease activation during crossover resolution.

## Materials and methods

### Media and chemicals

*S. cerevisiae* yeast strains used in this study were grown at 30°C in yeast extract-peptone-2% dextrose (YPD) or yeast extract-peptone-2% lactate (YPL) media (Rose et al. 1990). When required, geneticin (Invitrogen, San Diego), nourseothricin (Werner BioAgents, Germany), or Hygromycin B (Invitrogen, San Diego) were added to media at recommended concentrations (Goldstein and McCusker, 1999). Sporulation media was prepared as described (Argueso et al. 2004).

### Plasmids

Plasmids for this study are shown in Supplementary File 1 and oligonucleotides used to construct plasmids are listed in Supplementary File 2. All oligonucleotides used in this study were purchased from Integrated DNA Technologies (Coralville IA). The yeast genes present on these plasmids are from the SK1 strain background (Kane and Roth 1974).

pEAI522 (7.8 kb; *pCLB1-MLH3*) was created through a six fragment HiFi DNA Assembly (New England Biolabs) with *Bam*HI digested pUC18 as the backbone and the following five fragments: (1) 560 bp homology 5’ of *MLH3* ORF amplified from the SKI *S. cerevisiae* genome with AO4954/AO4955, (2) 540 bp *CLB1* promoter amplified from SKI genome with AO4956/AO4957, (3) full *MLH3* ORF consisting of 2288 bp with 100 bp downstream of the stop triplet amplified from SKI genome with AO4958/AO4959, (4) *KANMX* amplified from genomic prep of *KANMX* containing SKI with AO4960/AO4961, (5) 560 bp homology downstream of *MLH3* ORF amplified from SKI genome with AO4962/AO4963. Digestion of pEAI522 with *Xba*I and *Eco*RI yielded the DNA fragment used for integration.

pEAI523 (7.7 kb; *pGAL1-MLH3*) was created in the same manner as pEAI522 except for fragment (2) which was the *GAL1* promoter amplified from SKI genome with AO4965/AO4966. To yield compatible fragment overhangs, 560 bp homology 5’ of *MLH3* ORF was amplified from SKI genome with AO4954/AO4964 (fragment 1); likewise, the full *MLH3* ORF consisting of 2288 bp with 100 bp downstream of the stop triplet amplified from SKI genome with AO4967/AO4959 (fragment 3). All other fragments are identical to pEAI522.

pEAI524 (8.4 kb; *pGAL1-MLH3-13X Myc*) was created using a two fragment HiFi assembly. pEAI523 was amplified with AO5465/AO5466 to serve as the backbone. A 4.7 kb insert was amplified from pFA6a (Longtine et al. 1998; Bahler et al. 1998) with AO5467/AO5468 to fuse the 13XMyc tag to the C-terminal residues of *MLH3*.

pEAM370 (8.9 kb; *pADH1-MLH1-2µ-URA3*) was designed by a two-step HiFi assembly procedure. First, the *MLH1* ORF along with 500 bp flanking sequence 5’ relative to the *MLH1* start codon was amplified from the SK1 genome with AO5249/AO5250 and inserted into pRS426 at *Eco*RI using HiFi assembly. In the second step, an *ADH1* promoter, amplified from pGADT7 (Supplementary File 1) with AO5375/AO5376, was inserted into the pRS426 2µ plasmid (Christianson et al. 1992) containing *MLH1* which itself was amplified with AO5373/AO5374 to delete the native *MLH1* promoter. The resulting plasmid contained an *ADH1* promoter driving expression of *MLH1* in a *2µ* plasmid.

pEAM371 (8.9 kb; *pADH1-MLH3-2µ-URA3*) was designed in the same manner as pEAM370 apart from the *MLH1* ORF being replaced by the *MLH3* ORF. The *MLH3* ORF along with 500 bp flanking sequence 5’ relative to the *MLH3* start codon was amplified from the SK1 genome with AO5251/AO5252 and inserted into pRS426 at *Eco*RI using HiFi assembly. The *ADH1* promoter was amplified from pGADT7 with AO5379/AO5380 inserted into the pRS426 2µ plasmid containing *MLH3* amplified with AO5377/AO5378 to delete the native *MLH3* promoter.

pEAM372 (12 kb; *pADH1-MLH1-MLH3-2µ-URA3*) was designed with HiFi assembly using fragments from pEAM370 and pEAM371. pEAM370 was amplified with AO5410/AO5411 whereas the *pADH1-MLH3* fragment from pEAM371 was amplified with AO5412/AO5413 to insert *pADH1-MLH3* downstream of *pADH1-MLH1*. The resulting plasmid contained both *MLH1* and *MLH3* ORFs in tandem, both driven by *ADH1* promoters in a *2µ* plasmid.

pEAM373 (*pADH1-DMC1-2µ-URA3*) was designed with HiFi assembly using pEAM370 as the backbone, amplified with AO5414/ AO5415. The *DMC1* ORF was then amplified from the SKI genome with AO5416/AO5417. The resulting plasmid contained the *DMC1* ORF driven by the *ADH1* promoter in a *2µ* plasmid.

pEAA734 (6.6 kb; *DMC1, CEN6-ARSH4, URA3*) was constructed using two-fragment HiFi assembly (New England Biolabs). The *DMC1* open reading frame with 300 bp upstream and 296 bp downstream sequences was amplified from the SK1 genome with AO4920/AO4921 and cloned by HiFi assembly into pRS416 (Christianson et al. 1992) digested with *Bam*HI to generate pEAA734. *dmc1* mutations were introduced into pEAA734 using Q5 site directed mutagenesis (New England Biolabs).

Yeast two-hybrid vectors pEAM350-369 (Supplementary File 1) were assembled using HiFi assembly primers (Supplementary File 2) designed for N-terminal *GAL4* activation domain fusions (pGADT7), C-terminal *GAL4* activation domain fusions (pGADCg), N-terminal *GAL4* binding domain fusions (pGBKT7), and C-terminal *GAL4* binding domain fusions (pGBKCg) to the complete open reading frames of *MLH1*, *MLH3*, *DMC1*, *RAD51*, and *PMS1*. *MLH*-N-terminal domain fusions (NTT) to these vectors were constructed that contained amino acids 1 to 504 of MLH1 and 1 to 476 of MLH3. *MLH* C-terminal domains fusions (CTT) to these vectors were constructed that contained amino acids 345 to 769 of Mlh1 and 375 to 715 of Mlh3.

The above plasmid sequences were confirmed with Sanger sequencing or Oxford Nanopore long read sequencing using Plasmidsaurus (https://www.plasmidsaurus.com).

### Strains

Isogenic yeast strains used in this study were from the SK1 background (Kane and Roth 1974) except for strains used for *RTK1* tetrad analysis which are SK1 congenic (Argueso et al. 2004; Supplementary File 3). Gene knockouts in yeast were generated as described below apart from EAY4651-EAY4653 (*MATa, mlh2*Δ::*KANMX*), which were generated by integrating a *Xba*I and *Eco*RI restriction digest fragment of pEAI480 (Supplementary File 1) using standard methods (Rose et al. 1990; Goldstein and McCusker 1999; Gietz et al. 1995). Genomic integration events were confirmed by PCR amplification of genomic extracts as described previously (Hoffman et al. 1987). Mutant *dmc1* alleles were generated through Q5 site-directed mutagenesis (New England Biolabs) of pEAA734 (*ARS::CEN*, *URA3*) and transformed into EAY4911-EAY4913/EAY4914-EAY4916 diploids. Plasmid selection was maintained though omission of uracil in all growth and sporulation media.

*dmc1-T159A* was introduced to EAY4619 via integration of a *Hind*III digested fragment of pNH301-T159A. After selection for Ura^+^ colonies to identify plasmid integration events upstream of the *DMC1* ORF, transformants were screened on 5-FOA media to select for single-strand annealing-derived pop-outs of the *URA3* and *KANMX* markers. Transformants were verified to be G418^−^, Ura^−^, and capable of growing on YPL (Lactate^+^). Integration of the *dmc1-T159A* allele was further confirmed by Sanger sequencing of a PCR product using AO4180/AO4181 to generate the amplicon sequenced with AO4931 and AO4932. Promoter swaps were achieved through integration of the appropriate plasmids (Supplementary File 1) designed as described above.

### Spore-autonomous fluorescence assay

Crossing over in the *CEN8-THR1* interval on Chromosome VIII was assessed in SKY3576/SKY3575 derived diploids using spore-autonomous fluorescence reporter constructs (Thacker et al. 2011). Briefly, haploids of opposite mating type were struck to singles on YPL plates and incubated at 30°C for 2-3 days. Single colonies for each mating type were then mixed in 25 µl of ddH_2_O, spotted on YPD plates and incubated at 30°C for 5 hrs. Diploids were selected for by streaking the spot to single colonies on minimal media lacking tryptophan and leucine and incubated at 30°C for 48 hrs. Diploids were then patched on YPD and allowed to form a lawn before being transferred to sporulation media plates. After 2-4 days, spores were treated with 0.5% NP40 and briefly sonicated before analysis using a Zeiss AxioImager.M2. At least 500 tetrads for each genotype were counted to determine the % tetratype (Supplementary Files 4, 5). At least two independent transformants were studied per genotype and the vast majority of genotypes were analyzed on separate days. When appropriate, *ARS-CEN* plasmid maintenance was achieved through growth and sporulation media lacking the necessary nutrient to ensure plasmid retention.

### Tetrad analysis

Diploids derived from SK1 congenic strains EAY1108/EAY1112 were sporulated using the zero*-*growth mating protocol (Argueso et al. 2003). Haploid parental strains were mixed in 25 µl ddH_2_O, mated on YPD for 5 hrs at 30°C, and then struck to singles on minimal plates lacking leucine and tryptophan and incubated at 30°C for 48 hrs. Diploids were then patched on YPD and allowed to form a lawn before being transferred to sporulation media plates and grown for 2-4 days at 30°C. Tetrads were dissected on minimal complete plates and then incubated at 30°C for 3 to 4 days. Spore viability was then assessed (Supplementary File 6). Spore clones were replica-plated onto relevant selective plates and assessed for growth after an overnight incubation. Genetic map distances were determined by the formula of Perkins (1949; Supplementary File 7). Statistical analysis was done using the Stahl Laboratory Online Tools (https://elizabethhousworth.com/StahlLabOnlineTools/).

### Gene dosage screen for haploinsufficiency

Gene knockout transformation cassettes were generated consisting of an *MX* antibiotic resistance marker flanked by 300 bp of upstream and downstream homology with respect to the open reading frame (ORF) of each gene of interest. These cassettes were amplified by PCR from genomic preps of the appropriate strains from the *Saccharomyces* genome deletion project (Giaever et al. 2014). In this collection, each ORF has been replaced with *KANMX4*. Other *MX* knockout cassettes (e.g., *NATMX*, *hphMX6*) were generated through marker swapping of appropriate deletion strains.

EAY3486 (Supplementary File 3), a *mlh3Δ*::*NATMX* strain carrying a gene encoding a cyan fluorescent protein (CFP) on chromosome VIII, was transformed with the PCR amplified knockout cassette. Cells were then plated on YPD-G418 plates and grown at 30°C for three days. At least two independent transformants were verified by confirming resistance to G418 and nourseothricin and PCR amplification using genomic preps of G418^r^/NAT^r^ transformants. For PCR verification, primers annealing 350 bp upstream and downstream of the ORF of the gene of interest were utilized to ensure integration at the proper locus. Haploids were then crossed to four strains each carrying a gene encoding a red fluorescent protein (RFP) on chromosome VIII. These four strains are as follows: EAY3252 (MLH3), EAY3255 (*mlh3*Δ), EAY3572 (*mlh3-42*), and EAY3596 (*mlh3-54*). Diploids were isolated by selecting on media lacking tryptophan and leucine and analyzed in the spore-autonomous fluorescence assay described below.

Our criteria for allele-specific interactions were those in which there was little to no change in percent tetratype in either an *MLH3* and *mlh3*Δ background, but there was a significant drop of percent tetratype in either *mlh3-42* or *mlh3-54* backgrounds. Significance was assessed within backgrounds by chi-squared test. To minimize ⍺ inflation due to multiple comparisons, we applied a Benjamini-Hochberg correction at a 1% false discovery rate (Benjamini and Hochberg, 1995). At least two independent transformants were analyzed for each genotype.

To compare trends across genetic backgrounds, percent tetratypes in one-copy backgrounds were first normalized to their respective two-copy background. The following equations were used to plot the strength of each genetic interaction while penalizing any deviation in crossover rates in control backgrounds (i.e., *MLH3* or *mlh3*Δ) to ensure crossover defects were specific to the interplay between *MLH3* and the candidate gene and not the result of a broader meiotic dysfunction:

X-coordinate: (1-normalized %TT in *mlh3-42*) - |1- normalized %TT in *MLH3*| - |1- normalized %TT in *mlh3*Δ|
Y-coordinate: (1-normalized %TT in *mlh3-54*) - |1- normalized %TT in *MLH3*| - |1- normalized %TT in *mlh3*Δ|

## 2-hybrid analysis

Yeast-two-hybrid assays were performed as previously described (Bentolila et al. 2021). Yeast-two-hybrid expression strains, PJ69-4a and PJ69-4α (Supplementary File 3), were individually transformed with *GAL4* activation domain fusion vectors or *GAL4* binding domain fusion vectors, respectively. Vectors lacking any protein fusions served as negative controls. Single haploid transformants were selected for by plating on selective media. Crosses were performed as previously described and diploids were selected for by streaking to singles on minimal media plates lacking leucine and tryptophan. Interaction assays were performed by growing test diploids overnight in 3 ml minimal media lacking leucine and tryptophan. The following day, cultures were diluted in ddH_2_O to OD_600_ 0.5, 0.05, and 0.005 and spotted on - leucine, -tryptophan plates to assess plating efficiency and -leucine, -tryptophan, -histidine, - adenine plates to assess growth after 3 days of incubation at 30°C.

## *In-vitro* co-immunoprecipitation

yMlh1-FLAG-His_10_-Mlh3-HA was purified using methods described in Rogacheva et al. (2014). yMlh1-FLAG-Pms1 was purified as described in Plys et al. (2012). 6XHis-yDmc1 and 6XHis-yRad51 (6xHis tag in Rad51 removed by a SUMO protease) were purified using previously published procedures (Steinfeld et al. 2019).

6xHis-hDMC1 was purified using the same procedure used to purify 6xHis-yDMC1 and is as follows. Rosetta 2 Bl21 (DE3) cells bearing pET11C-6xHis-hDMC1expression plasmid (a kind gift from Michael Sehorn) was grown to an OD_600_ between 0.6 and 0.8 at 37 °C. The temperature was lowered to 16 °C and cells were induced with 0.5 mM IPTG overnight. Cells were harvested by centrifugation and stored at −80°C. Cells were resuspended in buffer A (30 mM Na-Hepes pH 7.5, 1000 mM KCl, 5 mM MgCl_2_, 0.01% NP-40, 2 mM ATP, 10% Glycerol, 5 mM 2-Mercaptoethanol, 0.2 mM PMSF, and Protease Inhibitor cocktail) and lysed by freeze thaw. Cell lysate was sonicated on ice with 10 cycles with 65% duty cycle 15 sec on and 45 sec off. Lysate was centrifuged at 10,000 x g for 45 minutes at 4 °C. Clarified extract was mixed with pre-equilibrated Talon resin and incubated in batch for 1 hr at 4 °C. After 1 hr the resin was centrifuged and washed 2x with buffer A, followed by 2x washes with buffer B (30 mM Na-Hepes pH 7.5, 200 mM KCl, 5 mM MgCl_2_, 0.01% NP-40, 2 mM ATP, 10% Glycerol, 5 mM 2-Mercaptoethanol, 0.2 mM PMSF, and Protease Inhibitor cocktail). The resin was then added to a disposable column. The protein was eluted in buffer C (30 mM Na-Hepes pH 7.5, 1000 mM KCl, 5 mM MgCl_2_, 0.01% NP-40, 2 mM ATP, 200 mM Imidazole, 10% Glycerol, 5 mM 2-Mercaptoethanol, 0.2 mM PMSF, and Protease Inhibitor cocktail). Fractions were analyzed by SDS-PAGE and the peak fractions were pooled. The protein was further purified using a HiTrap Heparin HP column. The column was resolved using a 30 CV 20-100% gradient of buffer D (30 mM Na-Hepes pH 7.5, 0 mM KCl, 5 mM MgCl_2_, 0.01% NP-40, 2 mM ATP, 10% Glycerol, 5 mM 2-Mercaptoethanol, and 1 mM EDTA) and buffer E (30 mM Na-Hepes pH 7.5, 1000 mM KCl, 5 mM MgCl_2_, 0.01% NP-40, 2 mM ATP, 10% Glycerol, 5 mM 2-Mercaptoethanol, and 1 mM EDTA). Fractions were analyzed by SDS-PAGE. Peak fractions were pooled and diluted to ∼200 mM KCl and loaded on a HiTrap Q HP column. The column was resolved using a 30 CV 20-100% gradient of Buffer F (30 mM Na-Hepes pH 7.5, 0 mM KCl, 5 mM MgCl_2_, 0.01% NP-40, 2 mM ATP, 10% Glycerol, 5 mM 2-Mercaptoethanol, and 1 mM EDTA) and Buffer G (30 mM Na-Hepes pH 7.5, 1000 mM KCl, 5 mM MgCl_2_, 0.01% NP-40, 2 mM ATP, 10% Glycerol, 5 mM 2-Mercaptoethanol, and EDTA). The fractions were analyzed by SDS-PAGE and peak fractions were pooled and concentrated by centrifugation at ∼1,100 x g at 4 °C using a 10 kDa MWCO Vivaspin 6 column. The concentrated protein was stored in single use aliquots at −80 °C.

Invitrogen Dynabeads™ Protein G Immunoprecipitation Kit (Catalog No. 10007D) was used for immunoprecipitations involving purified yMlh1-Mlh3, yMlh1-Pms1, yDmc1, yRad51, and hDmc1. For all anti-FLAG immunoprecipitations, 2 µg of purified recombinase (yDmc1, yRad51, and hDmc1) were mixed with 1 µg MLH complex (yMlh1-Mlh3 or yMlh1-Pms1) in 100-400 µl total volume Binding/washing buffer at 4°C for 30 minutes. For DNase treated samples, 2 units of DNase I (NEB) was added along with MgCl_2_ to a final concentration of 2.5 mM. DNase I treated samples were incubated at 37°C for 5 minutes before incubating protein mixtures at 4°C for 30 minutes. If 1 µg lambda DNA (NEB) was added, to check for DNA degradation, 5 µl of DNase treated samples was removed after 37°C incubation, treated with Proteinase K, and incubated at room temperature for 15 minutes before running on an 1% agarose gel. While protein mixtures were incubating at 4°C, 50 μl (1.5 mg) Dynabeads^TM^ magnetic beads were prepared by removing the supernatant and incubating the beads at room temperature for 20 minutes with mild agitation in a total volume of 200 µl Binding/washing buffer with 5µg ANTI-FLAG® M2 antibody (Sigma, catalog number F1804). Beads were then washed once with Binding/washing buffer before addition of the pre-mixed proteins. The protein-bead-Ab mixture was incubated at room temperature for 30 minutes with mild agitation. The protein-bead-Ab mixture was then placed on a magnetic rack to remove the flow through fraction containing unbound proteins. 3X sample buffer supplemented with 50 mM DTT was added to the unbound fraction and proteins were loaded on a 10% SDS-PAGE gel. The protein-bead-Ab mixture was then washed 3X with 200 μl washing buffer and re-suspended in 100 μl washing buffer. The bead suspension was then transferred to a clean tube to prevent co-elution of protein bound to the tube wall. After removal of the supernatant, the beads were re-suspended in 30 µl elution buffer and 15 µl 3X sample buffer with 50 mM DTT was added. The beads eluate was heated at 99°C for 6 min and loaded on a 10% SDS-PAGE gel. Bands were imaged after Coomassie Blue staining.

### Meiotic time courses

SK1 yeast diploids were struck to singles on YPL plates and incubated at 30°C for 3 days. A single colony was used to inoculate a 3 ml YPD culture which was grown overnight to saturation. 700 µl of the saturated YPD culture was sub-cultured into 400 ml YPA (1% potassium acetate; 2% peptone; 1% yeast extract) and grown for 16-17 hrs at 30°C. For phenotypic analyses (e.g., spore autonomous assay of crossing over at *CEN8::THR1*), 50 ml of pre-sporulated cells were washed with water and resuspended in 25 ml CSHSPO (1% potassium acetate; 0.1% yeast extract; 0.05% glucose) in a 250 ml flask to ensure adequate aeration. For protein expression or co-immunoprecipitation experiments, 400 ml of pre-sporulated cells was washed with water and resuspended in ∼240 ml CSHSPO in a 2 liter flask at an OD_600_ of ∼1.5-2.5. All liquid sporulations were incubated at 30°C. T=0 for all time-courses denotes the time when sporulation cultures were transferred to a 30°C shaker. When appropriate, β-estradiol (Millipore-Sigma) was added to 10 nM (complementation) or 1 µM (overexpression). To monitor meiotic progression, ∼500 µl aliquots from each sporulation culture were collected at appropriate timepoints. Cells were then fixed in 900 µl 40% EtOH, 0.1 M Sorbitol and either incubated at room temperature for 30-60 minutes or stored at 4°C. After fixation, cells were washed twice with 1X PBS. 2 µl 0.5% NP40 and 1 µl DAPI (0.1 mg/ml stock) was added to cells prior to imaging and counting with a Zeiss AxioImager.M2. The number of cells with 1, 2, 3/4 nuclei were recorded. At least 600 cells were counted per genotype per timepoint except for the 0-hr timepoint where at least 300 cells were counted. Three independent transformants were analyzed per genotype and experiments were performed on separate days to avoid batch effects.

### Meiotic whole cell protein analysis

At appropriate timepoints, 10 OD_600_ units of cells were collected from sporulation cultures (∼30-50 mg cell mass). Cells were collected by centrifugation, washed once with 1 ml of 1X TE (10 mM Tris 8.0 1 mM EDTA) + 1 mM PMSF, and vortexed for 15 minutes at 4°C with 100 µl acid-washed glass beads (500 micron, Sigma G8772) and 200 µl lysis buffer (10% glycerol; 50 mM Tris-HCL pH 7.5; 0.2% NP-40; 150 mM NaCl; 5 mM EDTA; 1 mM PMSF; 1X Halt protease inhibitor cocktail (Thermo)). Lysate was cleared by centrifugation for 5 minutes at 4°C and 125 µl of cleared lysate was added to 50 µl 3X SDS protein sample buffer. Protein concentrations were determined with Bradford reagent (Thermo; Bradford 1976) and 20 µg total protein was loaded onto a 10% SDS-PAGE gel.

### Meiotic co-immunoprecipitation

1.2 × 10^9^ cells were harvested at the desired time-point in meiosis and washed with 1X TE+1mM PMSF. Cells were lysed in ∼2 ml lysis buffer (10% glycerol; 50 mM Tris-HCL pH 7.5; 0.2% NP-40; 150 mM NaCl; 5 mM EDTA; 1 mM PMSF; 1X Halt protease inhibitor cocktail (Thermo); 0 or 125 Units/mL Benzonase (Sigma) with ∼600 µl glass beads in a bead beater 4 times, 5 min each, with at least 5 min rest on ice between bead beating rounds. Lysate was cleared by centrifugation in a microfuge at 13,000 RPM for 5 min at 4°C. 25 µl of supernatant was collected for input and mixed with 12.5 µl 3X sample buffer w/ fresh 50 mM DTT. 10 µl (10 μg) of anti-Myc (clone 4A6, Sigma) was added to lysate and incubated at 4°C for 1 hr with rotation. 100 μl of Dynabeads™ Protein G were washed twice 2:1 with lysis buffer. Lysate/antibody mixture was then added to washed magnetic beads and incubated at 4°C for 3 hr with rotation. Beads were washed four times with 500 µl of lysis buffer. On the final wash step, the bead suspension was transferred to a clean tube. Washed beads were resuspended in 35 µl 3x SDS protein sample buffer w/ fresh 50 mM DTT and heated at 95°C for 10 minutes. The bead eluate was loaded onto a 10% SDS-PAGE gel.

### Western Blot Analysis

For both whole cell protein extract and immunoprecipitation analyses proteins electrophoresed in SDS-PAGE were transferred to a 0.2µm PVDF membrane (Thermo) at either 100V for 2 hrs, 70 V for 4 hrs, or 30V overnight. Proteins tagged with the Myc epitope were detected with mouse anti-Myc (1:1000; clone 4A6, Sigma) and peroxidase conjugated anti-mouse secondary antibody (1:20,000, Sigma). Dmc1 was detected with goat anti-Dmc1 (1:2000; kindly provided by Doug Bishop) and peroxidase conjugated anti-goat (1:5000; Thermo). Glucose-6-phosphate dehydrogenase was detected with rabbit anti-G6PDH (1:10,000; Sigma) and peroxidase conjugated anti-rabbit (1:20,000, Invitrogen). Signal was detected with Clarity Western ECL Substrate (Bio-Rad) with either film or ChemiDoc MP (Bio-Rad) and quantified with Image Lab software (Bio-Rad).

## Results

### A dosage suppression screen for *MLH3* interactors reveals known and novel interactions

We performed a haploinsufficiency screen in *mlh3-42* (*mlh3-R552A, D553A, K555A, D556A*) and *mlh3-54* (*mlh3-R682A, E684A*) genetic backgrounds (Al-Sweel et al. 2017). These alleles were chosen because they conferred separation of function phenotypes that we hypothesized would reveal synthetic phenotypes for crossing over with other factors that were also compromised for function. Strains bearing these alleles were proficient for meiotic crossing over but were defective in *MLH3*-dependent mismatch repair and were disrupted for Mlh1-Mlh3 interactions as measured by yeast two-hybrid and protein purification analyses (Al-Sweel et al. 2017).

Both *mlh3-42* and *mlh3-54* map to the structured C-terminal domain of Mlh3 which binds to Mlh1 (Fig. 2A; PDB 6RMN; Dai et al. 2021). Residues 552-556 mutated in *mlh3-42* map distal to the dimerization interface between Mlh1 and Mlh3 while residues 682-684 mutated in *mlh3-54* map proximal to the interface. Dai et al. (2021) identified three regions within the Mlh3 C-terminus predicted to mediate the formation of an Mlh1-Mlh3 polymer; mutations in these regions disrupt meiotic crossing over. Interestingly, Mlh3 residues 552-556 map within a loop opposite to the second region (Fig. 2B). Based on the above structural information, we hypothesize that *mlh3-54* destabilizes the canonical Mlh1-Mlh3 binding interface while *mlh3-42* indirectly impairs Mlh1-Mlh3 functions by altering the conformation of the Mlh3 C-terminus to disrupt Mlh1-Mlh3 polymer formation.

**Fig. 2.**
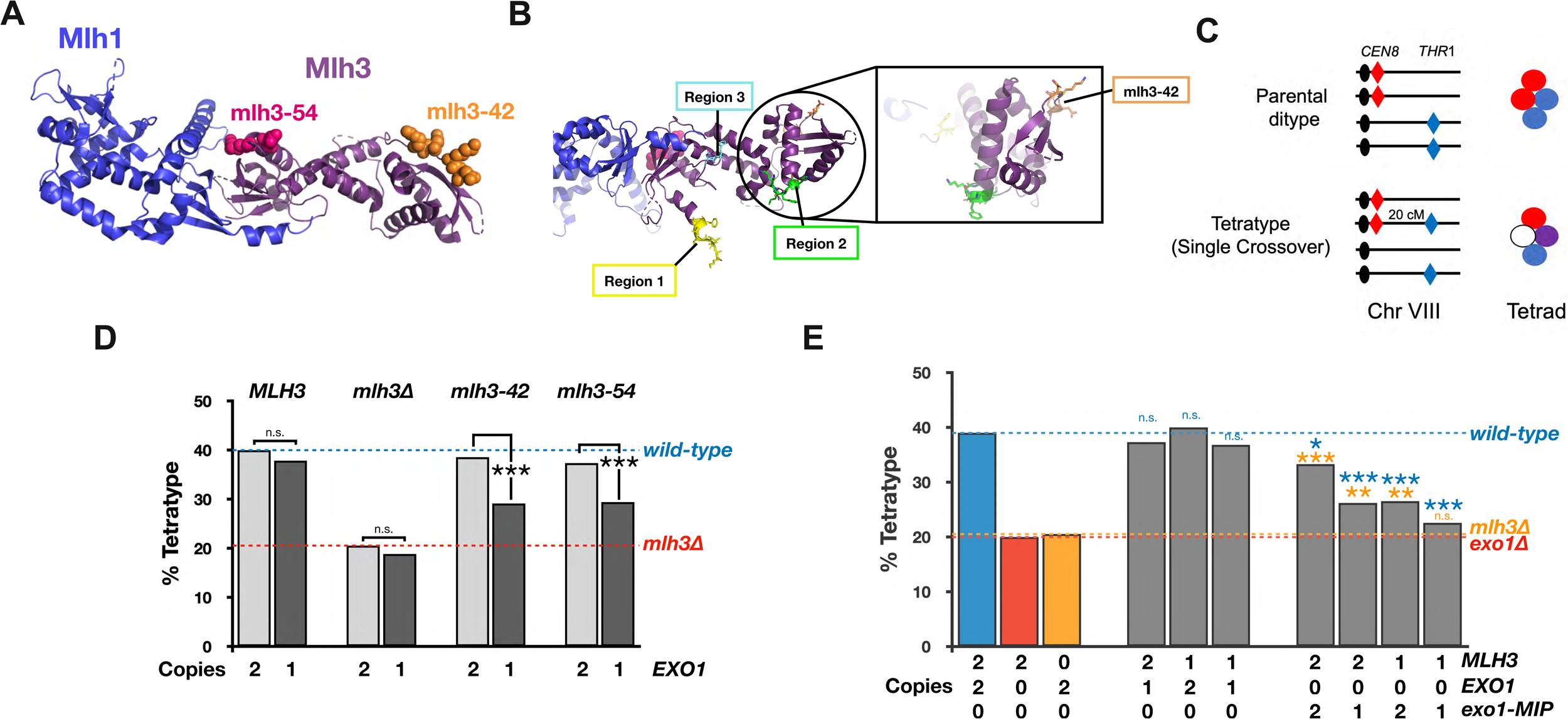
Rationale for using *mlh3-42* and *mlh3-54* as baits (MMR-CO+) in gene dosage suppression analysis. A. Crystal structure of the Mlh1-Mlh3 C-terminus (PDB 6RMN, Dai et al. 2020) with mutated residues highlighted by space filling (R552A, D553A, K555A, D556A for mlh3-42 and R682A, E684A for mlh3-54). B. Three regions within the Mlh3 C-terminus predicted to mediate the formation of an Mlh1-Mlh3 polymer (Dai et al. 2020) with amino acids changed in mlh3-42 highlighted as sticks for resolution purposes. C. Spore-autonomous fluorescence assay was used to measure tetratypes (single crossover events) in the *CEN8-THR1* interval on chromosome VIII (Thacker et al. 2011). D. Crossover analysis in the *CEN8-THR1* interval on chromosome VIII expressed as percent tetratype for indicated *EXO1* gene dosage in *wild-type*, *mlh3Δ, mlh3-32*, and *mlh3-54* strains. Dashed lines indicate *wild-type, mlh3Δ*, and *exo1Δ* levels of crossing over. E. Crossing over in the *CEN8-THR1* interval on chromosome VIII expressed as percent tetratype for indicated gene dosages of *MLH3, EXO1*, and *exo1-MIP*. Significance was assessed according to X^2^ p-values where p>0.05 = n.s. (not significant), p<0.05 = *, p<0.01 = **, and p<0.001 = ***. Blue asterisks indicate comparisons to *wild-type* and orange asterisks indicate comparisons to *mlh3*Δ. To account for multiple comparisons, we applied a Benjamini-Hochberg correction at a 5% false discovery rate (Benjamini and Hochberg, 1995).

The phenotypes exhibited by *mlh3-42* and *mlh3-54* mutants suggested to us that Mlh1-Mlh3 could be structurally stabilized by meiotic factors which promote its pro-crossover functions. An alternative hypothesis is that is that the pro-crossover function of Mlh1-Mlh3 is less dependent on complex stability than its MMR function. If our first hypothesis is correct, then lowering the protein abundance of factors that interact with Mlh3 would elicit allele-specific crossover defects. Because Mlh1-Mlh3 and Exo1 form a constitutive complex in meiotic prophase (Sanchez et al. 2020), we tested as proof of principle the effect of lowering gene dosage of *EXO1* on crossing over in strains containing the *mlh3-42* and *mlh3-54* mutations.

We assessed effects of haploinsufficiency by measuring single crossover frequencies (tetratype class, abbreviated as TT) using spore autonomous fluorescent markers inserted in the 20 cM *CEN8-THR1* interval on chromosome VIII (Fig. 2C; Thacker et al. 2011; Al-Sweel et al. 2017). As shown in Fig. 2D, reducing *EXO1* from two to one copy in *mlh3-42* (39% TT with two copies compared to 29% TT with one) or *mlh3-54* (37% TT with two copies compared to 29% with one) led to significant decreases in crossing over. Lowering *EXO1* dosage *in MLH3* and *mlh3Δ* strains did not significantly affect crossing over. To complement this analysis we examined *MLH3-EXO1* interactions using the *exo1-MIP* allele (*F447A,F448A*) which disrupts interactions between Exo1 and Mlh1 and confers a meiotic crossover defect (Amin et al. 2001; Gellon et al. 2002; Tran et al. 2007, Zakharyevich et al. 2010). We detected an intermediate crossover defect (33% TT) in strains containing two copies of both *exo1-MIP* and *MLH3*; however, strains containing only single copies of *exo1-MIP* and *MLH3* displayed crossover frequencies (23% TT) indistinguishable from *mlh3*Δ (Fig. 2E). We concluded that the reduction in crossover rates due to lowering the gene dosage of partner proteins in *mlh3-42, mlh3-54* and *exo1-MIP* backgrounds was likely due to compromising the integrity of the Exo1-Mlh1-Mlh3 complex.

The above observations encouraged us to test haploinsufficiency of 34 meiotic genes in the *mlh3-42* and *mlh3-54* backgrounds (Supplementary File 3). Genes were chosen to represent various stages of meiotic recombination, including helicases, ZMMs, structure selective nucleases (SSNs), synaptonemal complex factors, and those involved in the recombinase machinery and chromosome segregation. We generated haploid knockouts in the *mlh3*Δ strain EAY3486 and crossed the resulting strains to *wild-type* (EAY3252), *mlh3*Δ (EAY3255), *mlh3-42* (EAY3572), and *mlh3-54* (EAY3596) to generate diploids harboring single copies of the 34 genes. These strains also contained markers to detect crossing over in the *CEN8-THR1* interval (Supplementary File 3). For all 34 genes we analyzed the statistical significance of reductions in crossing over in each *MLH3*/*mlh3* background, comparing two copies of the gene candidate to one copy, and categorized the strength of haploinsufficiency according to χ^2^ p-values, following Benjamini-Hochberg (Benjamini and Hochberg 1995) correction for multiple comparisons, where p>0.05 = not haploinsufficient, p<0.05 = mildly haploinsufficient, p<0.01 = moderately haploinsufficient, and p<0.001 = strongly haploinsufficient (Supplementary File 4). Lowered gene dosage rarely led to a significant drop in tetratype frequencies in *wild-type* or *mlh3*Δ backgrounds. Two exceptions were the ZMM gene *MER3* which conferred haploinsufficiency in the *MLH3* background (30% TT; p<0.001) and the Rad51 nucleoprotein filament disassembly gene *SRS2*, which conferred haploinsufficiency in the *mlh3*Δ background (14% TT; p<0.01; Krejci et al. 2003; Crickard et al. 2018; Hunt et al. 2019).

Eleven and eight genes displayed haploinsufficiency in *mlh3-42* and *mlh3-54* backgrounds, respectively, with six genes conferring phenotypes in both backgrounds (*EXO1, MSH5, ZIP3, DMC1, RDH54*, and *CHD1*). To quantify the strength of the genetic interactions, we compared trends across backgrounds by plotting normalized crossover rates (Fig. 3A; Materials and methods). We developed a scoring method for each candidate by plotting the decrease in crossover rates in *mlh3-42* and *mlh3-54* backgrounds; this analysis included a penalty for deviations in crossover rates not specific to the *mlh3* alleles (changes seen in *wild-type* or *mlh3*Δ; Materials and methods). Most gene candidates clustered to the bottom left of the graph in Fig. 3A, indicating a lack of genetic association in both *mlh3* allele backgrounds. We focused our attention on genes with the strongest genetic interactions in both *mlh3* allele backgrounds (top right, Fig. 3A). Known physical interactors of Mlh3 clustered within this group including *EXO1*, *MSH5*, *PMS1* (Mlh1-Pms1 forms the major MMR complex), and newly identified interactors such as the *CHD1* chromatin remodeler and the putative kinase *RTK1* (Wild et al. 2019). Additionally, *NDJ1*, which regulates telomere bouquet formation and chromosome pairing in meiosis, was in this group (Trelles-Sticken et al. 2000) as were three factors that act in or stimulate strand invasion steps in meiotic recombination, *DMC1*, *RAD51*, and *RAD54.* We cross-referenced these nine genes for statistical significance in the *MLH3/mlh3* backgrounds (Fig. 3B; Supplementary Fig. 1). When analyzed through both scoring methods, we determined that the strongest candidates for genetic interaction were *EXO1*, *DMC1*, *MSH5*, *PMS1*, and *RTK1*. The allele-specific haploinsufficiency conferred by *DMC1* was remarkably similar to that seen by *EXO1* (compare Fig. 2D to Fig. 3C). Given the temporal separation between the functions of Dmc1 and Mlh3 in most models for meiotic recombination (Fig. 1), we regarded the *MLH3-DMC1* interaction as unexpected and pursued it further. We performed an additional genetic analysis of *RTK1* but concluded that Rtk1 is not essential for either Class I or Class II crossovers but may function to partially stabilize Mlh1-Mlh3 (Supplementary Fig. 2).

**Fig. 3.**
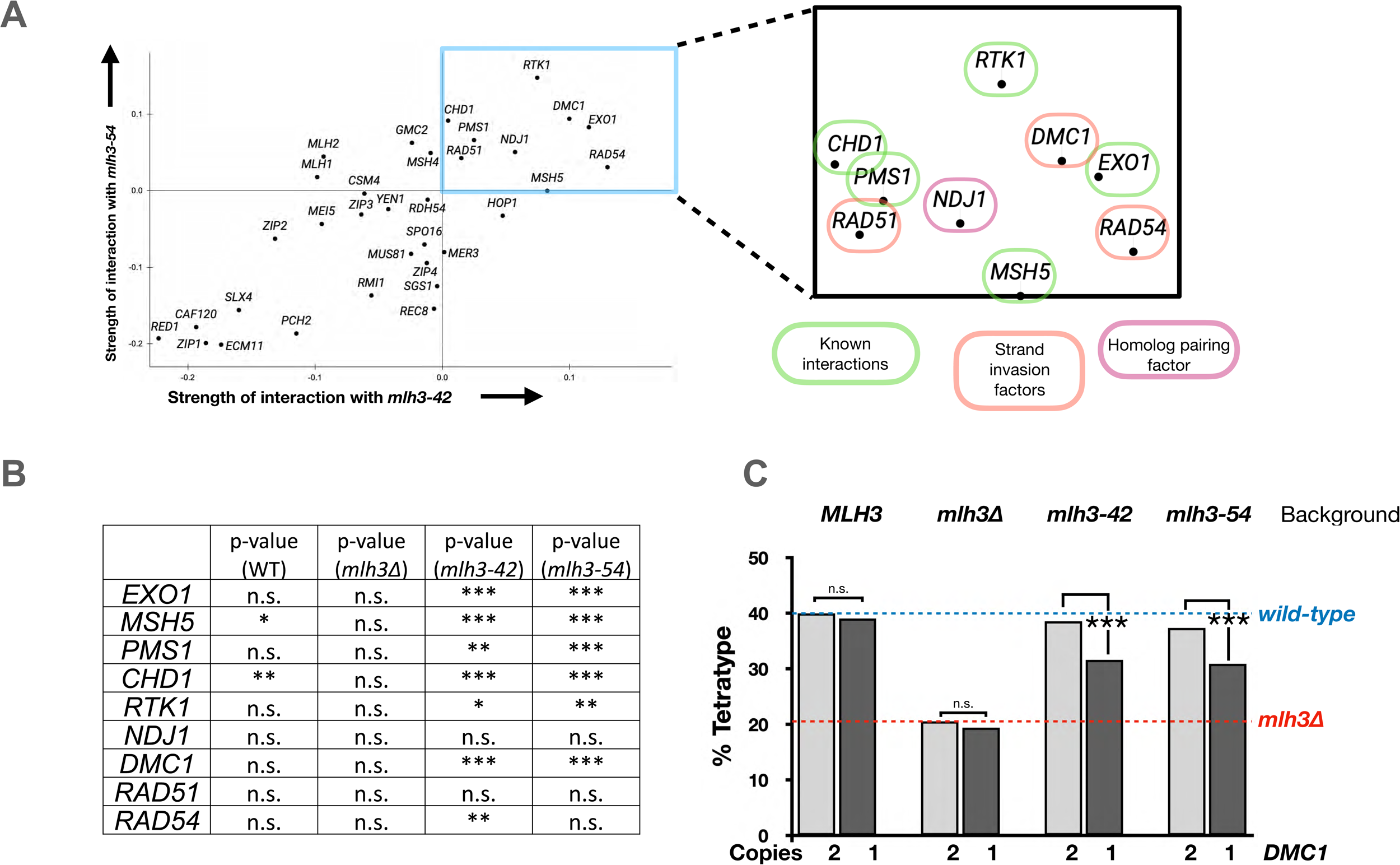
*EXO1, MSH5, PMS1, RTK1, and DMC1* are allele-specific interactors of *MLH3*. A. Analysis of the strength of genetic interactions with *mlh3-42* (x-axis) and *mlh3-54* (y-axis) for 34 meiotic genes from various stages of meiotic prophase. See Materials and methods for rationale and Supplementary File 4 underlying data for raw tetrad counts and significance measurements. B. Statistical confirmation of allele-specific interactions (see Materials and methods). C. *DMC1* displays dosage-specific interaction with *mlh3-42* and *mlh3-54* strains. Dashed lines indicate *wild-type* and *mlh3*Δ levels of crossing over. Significance was assessed as described in the Materials and methods.

### Yeast Mlh1-Mlh3 physically interacts with yeast Dmc1 and yeast Rad51 but not with human Dmc1

We purified yeast (y) Mlh1-Mlh3, yMlh1-Pms1, yRad51, yDmc1 and human (h) DMC1 and tested them for interactions by co-immunoprecipitation. yMlh1-Mlh3 robustly immunoprecipitated yDmc1 and yRad51 but not hDMC1 (Fig. 4A, B). The yMlh1-Mlh3-yDmc1 interaction was retained in the presence of DNase I and in the absence of ATP, suggesting that the interaction was unlikely to be mediated through DNA (Supplementary Fig. 3; Busygina et al. 2013). Like yMlh1-Mlh3, yMlh1-Pms1 also immunoprecipitated yDmc1, suggesting that yMlh1 was likely important for interactions with yDmc1 (Fig. 4C).

**Fig. 4.**
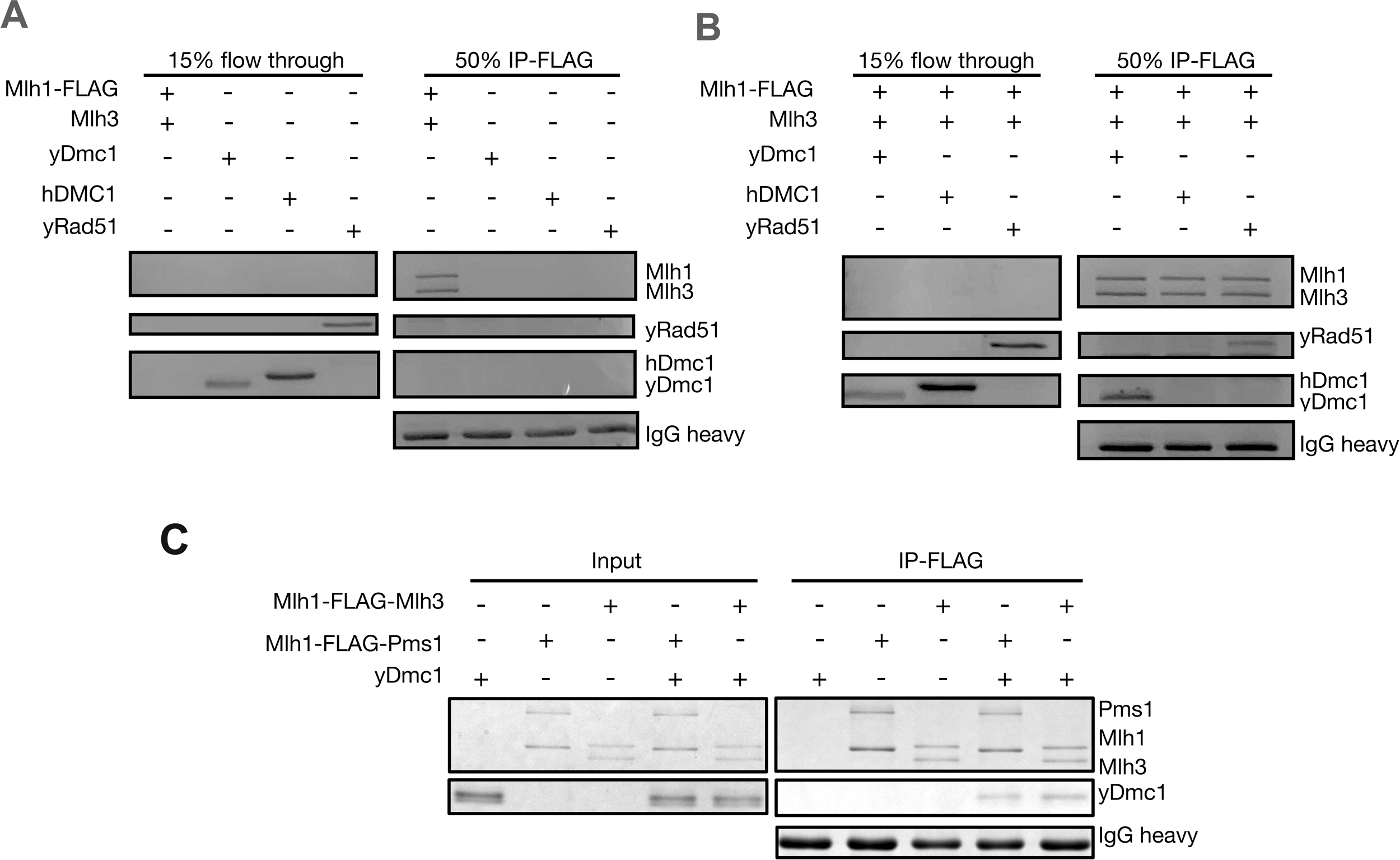
*In vitro* co-immunoprecipitation reactions performed with yMlh1-FLAG-Mlh3, yMlh1-FLAG-Pms1, yDmc1, yRad51, and hDmc1. A. Unbound protein flow throughs showing that yDmc1, yRad51 and human Dmc1 do not bind non-specifically to Dynabeads Protein G conjugated to anti-FLAG M2 antibody. B, C. Immunoprecipitation reactions containing the indicated purified proteins and Dynabeads Protein G conjugated to anti-FLAG M2 antibody. In all panels proteins were electrophoresed in 10% SDS-PAGE. See Materials and methods for details.

Our genetic analysis and *in vitro* pulldown experiments encouraged us to test if Mlh3 physically interacts with Dmc1 during meiosis through co-immunoprecipitation experiments. We used a Dmc1 polyclonal antibody to detect soluble Dmc1 present in cell extracts lysed in physiological (150 mM) NaCl concentrations (Fig. 5A; Materials and methods). We then used an anti-Myc antibody to immunoprecipitate Mlh3-13xMyc from cell extracts prepared from cultures harvested 0-8 hrs after transfer to sporulation media. We only detected Mlh3 in Myc-tagged strains; untagged immunoprecipitates contained no cross-hybridizing bands. Interestingly, we detected a weak band corresponding to Dmc1 in Mlh3 pulldowns from extracts prepared 4 hrs after inducing sporulation, but a more robust Dmc1 band was detected at 6 hrs (Fig. 5B). The Mlh3-Dmc1 interaction was retained when cell extracts were treated with Benzonase, confirming that DNA is not required for co-immunoprecipitation. We concluded that Mlh3 interacts physically with Dmc1 in meiotic prophase.

**Fig. 5.**
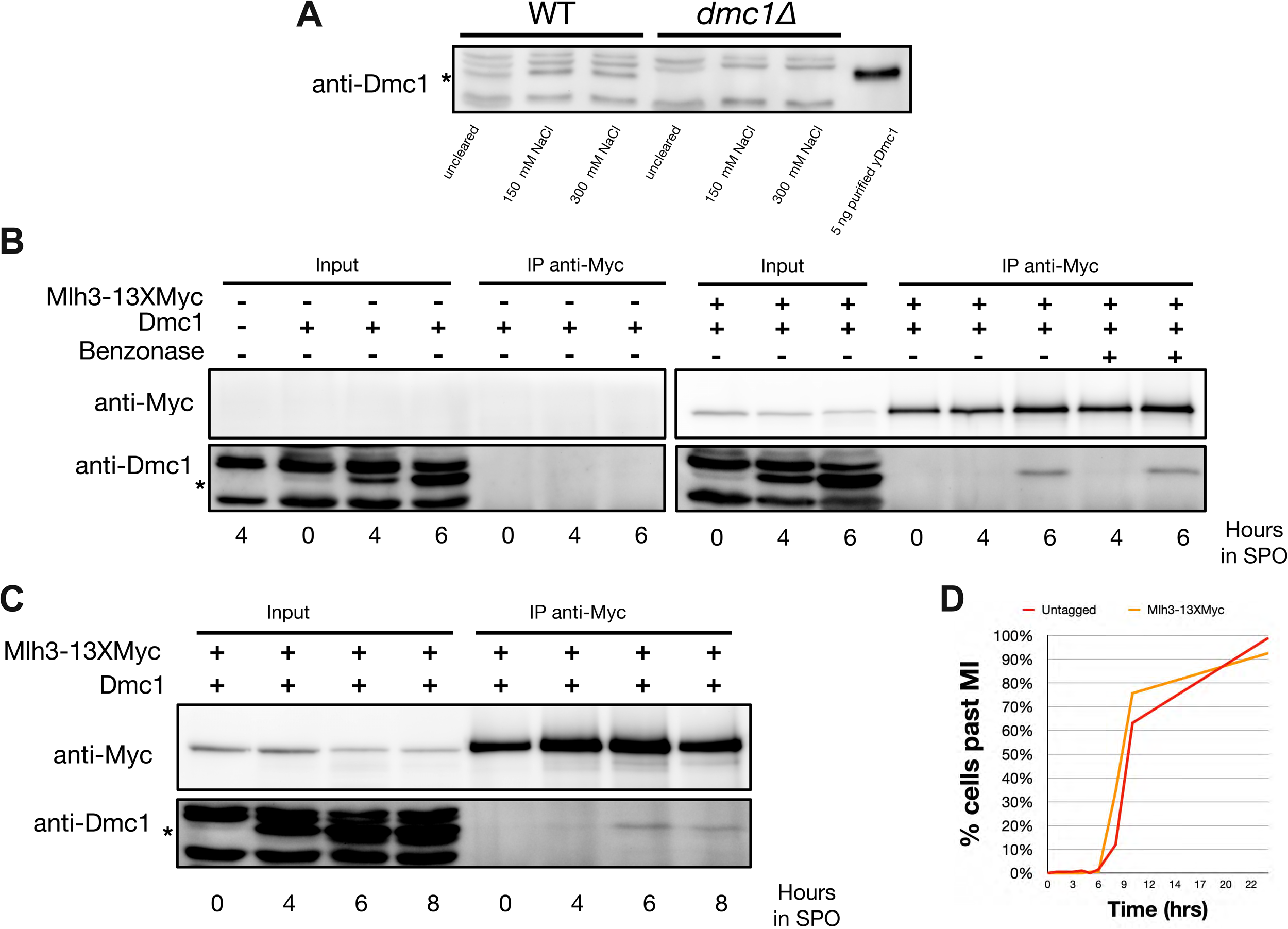
Mlh3 interacts with Dmc1 *in vivo*. A. Western blot showing specificity of yDmc1 antibody. 20 μg of whole cell extracts prepared in the indicated NaCl concentrations were loaded per lane. 5 ng of purified yDmc1 was loaded in the yDmc1 lane. B. Time course pulldown of C-terminally tagged *Mlh3-13xMyc* at 0, 4, 6 hr following transfer to sporulation media. Controls include *dmc1Δ* input (lane 1, left most lane) and inputs/pulldowns for untagged, *wild-type* strains (lanes 2-7). Inputs/pulldowns for Mlh3-13xMyc are in lanes 8-13. Lanes 14-15 show Mlh3-13xMyc pulldowns at 4 and 6 hr when Benzonase was added to lysis buffer (125 Units/mL). C. Time course pulldown showing inputs/pulldowns for Mlh3-13xMyc at 0, 4, 6, 8 hr. D. Meiotic progression, as measured by completion of Meiosis I, is presented for the Mlh3-13X Myc tagged strain compared to an untagged strain. The amount of sample loaded in panels B and C were normalized based on cell mass.

We tracked meiotic progression in sporulating cultures by DAPI staining of nuclei at 0, 4, 6, and 8 hrs post meiotic induction. We detected the Dmc1-Mlh3 interaction at 6 and 8 hrs (Fig. 5C). Meiotic progression was delayed by ∼2 hrs relative to time courses performed with smaller culture volumes (compare Fig. 5D to 6D). This delay was also observed in the untagged *wild-type* strain. We inferred that a difference in methods used to sporulate small vs. large meiotic cultures accounts for the 2 hr delay rather than a phenotype associated with the tagged strain utilized for pulldown experiments (see Discussion).

**Fig. 6.**
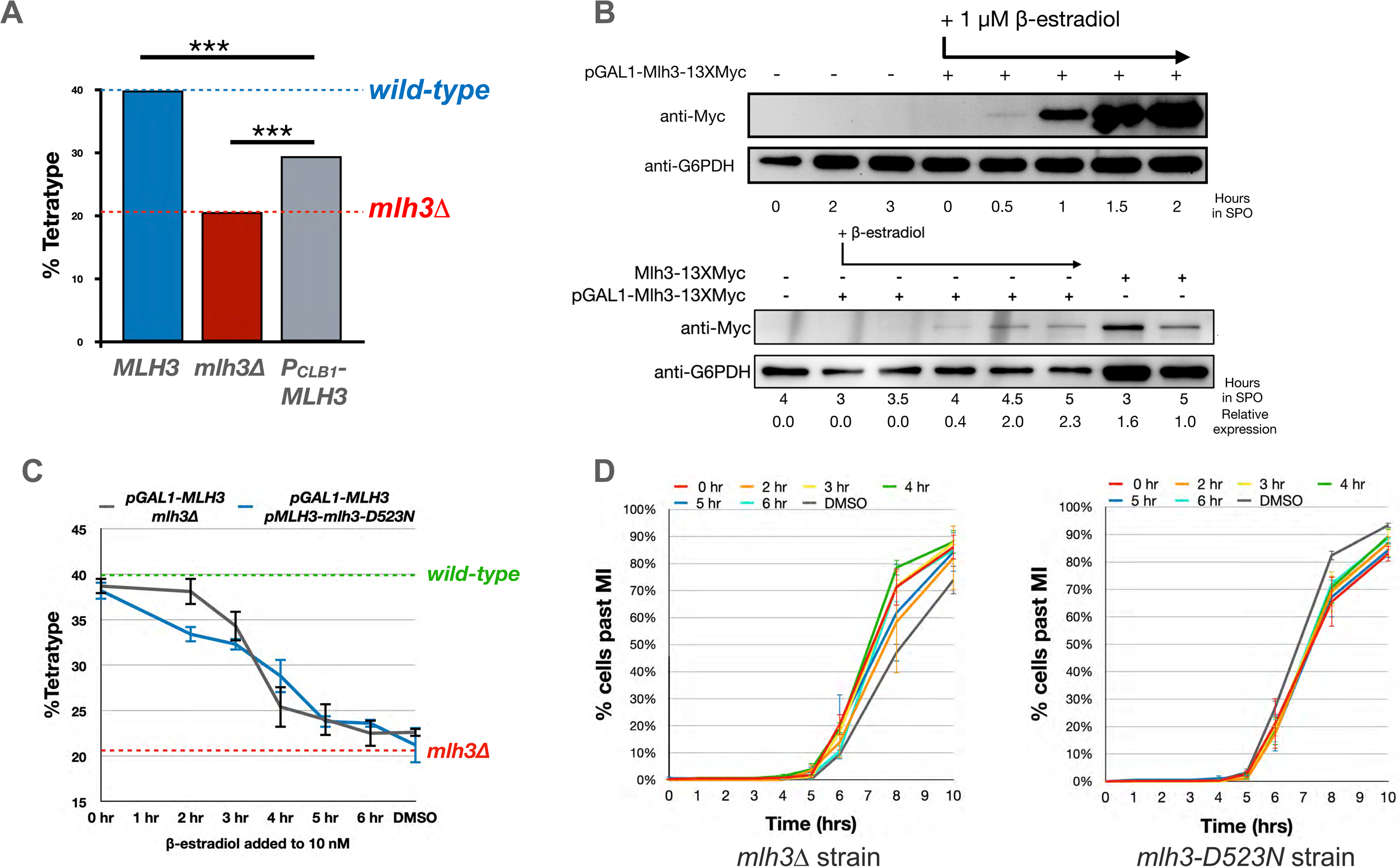
Modulating expression of *MLH3* in meiotic prophase. A. *MLH3* expression was placed under the control of the cyclin *CLB1* promoter to drive late meiotic prophase expression. Crossing over in the *CEN8-THR1* interval is presented. B. Western blot showing temporal lag between addition of β-estradiol (1μM, added at T= 0 hr, top, and 10 nM, added at T = 3 hrs, bottom) and subsequent Mlh3 protein expression. C. Comparison of crossover rates for strains expressing *MLH3* from the *GAL1* promoter upon addition of β-estradiol to sporulation media to a final concentration of 10 nM from 0 to 6 hr post transfer to sporulation media. Error bars represent standard error for three independent transformants analyzed on different days. D. Meiotic progression, as measured by completion of Meiosis I (MI), of sporulated cultures presented in panel C. Error bars represent standard error for the same three independent experiments shown in Panel C. Underlying data can be found in Supplementary File 4 and Supplementary File 5.

### Delayed expression of *MLH3* causes defects in meiotic crossing over

We performed an alanine scan mutagenesis of *DMC1* with the goal of identifying *dmc1* alleles which phenocopy the *DMC1* haploinsufficiency phenotype observed in *mlh3-42* and *mlh3-54* backgrounds. This was performed by: (i) Aligning yDmc1 to human DMC1 (Supplementary Fig. 4A). (ii) Locating primarily charged residues that were not conserved between the yeast and human proteins. (iii) Mapping residues to the yDmc1 postsynaptic complex (PDB: 7EJ7) to confirm they were surface exposed and not part of Dmc1’s self-interacting, or DNA binding interfaces (Steinfeld et al. 2019; Xu et al. 2021; Supplementary Fig. 4B, C). 21 *dmc1* alleles were analyzed for crossing over in the *CEN8-THR1* interval. While one allele, *dmc1-K101A,* mildly reduced crossing over and deleting the first 21 amino acids conferred a null phenotype, no other mutations tested conferred significant defects in crossing over (Supplementary Fig. 4D).

Because the *DMC1* mutagenesis did not identify specific residues that were likely to be important for Mlh1-Mlh3-Dmc1 interactions, and two-hybrid analysis was not successful in detecting such interactions (Supplementary Fig. 5), we performed experiments aimed at disrupting the Mlh3-Dmc1 interaction by modulating the timing of *MLH3* expression. We thought that if Mlh3 interacts with Dmc1 on recombination intermediates, such a connection would likely be established shortly after strand invasion when Dmc1 is highly expressed and the local concentration of the two factors is likely to be high. Dmc1 levels reach a peak of ∼90,000 molecules/diploid cell in meiosis (Chan et al. 2019) while Mlh3 is constitutively expressed in meiosis at levels similar to that seen in mitosis (∼500 copies per haploid cell; Ho et al. 2018; Wild et al. 2019; Fig. 5). We hypothesized that delaying *MLH3* expression to later timepoints in meiosis would impair Mlh3’s ability to form a functional interaction with Dmc1 and disrupt its crossover functions. Such a phenotype contrasts with studies showing that late meiotic expression of unrestrained alleles of the *YEN1* SSN (Yen1 does not act in the Class I crossover pathway) could restore crossing over in the *CEN8-THR1* interval in *mlh1Δ* and *mlh3Δ* strains (Arter et al. 2018; Wild et al. 2019).

We confirmed that *MLH3* is expressed constitutively in meiosis (Fig. 5B, C; Wild et al. 2019). We delayed *MLH3* expression in meiosis by placing it under the control of the *CLB1* cyclin promoter, which is activated by the *NDT80* transcription factor at a time when dHJs are resolved into crossovers (Chu and Herskowitz, 1998; Brar et al. 2012; Supplementary Fig. 6A). We monitored crossing over in the *CEN8-THR1* interval in *P_CLB1_-MLH3* strains. As shown in Fig. 6A, crossing over was significantly impaired in *P_CLB1_-MLH3*. We then determined the temporal requirement of *MLH3* expression by placing *MLH3* under the control of the *GAL1* promoter in strains carrying a Gal4-Estrogen Receptor fusion (Benjamin et al. 2003). In this construct Mlh3 contains 13 copies of the Myc tag at its C-terminus (*P_GAL1_-Mlh3-13XMyc*;). Strains bearing this allele were partially functional for meiotic crossing over (30% TT; Supplementary File 4). In *P_GAL1_-Mlh3-13XMyc* strains *MLH3* expression was induced by addition of β-estradiol to sporulation media and Mlh3-13XMyc protein levels were monitored by Western blot at various times after β-estradiol addition. We performed two different types of estradiol time courses (Fig. 6B). In the first we induced *Mlh3-13XMyc* expression immediately after transfer to sporulation media using a high concentration of β-estradiol (1μM). Addition of β-estradiol at this concentration caused a significant delay in meiotic progression but did not impact meiotic crossing over (Supplementary Fig. 6B, C). In this experiment significant levels of Mlh3-13XMyc were observed 60 to 90 minutes after estradiol addition. In the second time course β-estradiol was added at 3 hrs after transfer to sporulation media at a concentration (10 nM) that fully complemented Mlh3 functions without causing a delay in meiotic progression (Fig. 6D). Mlh3-13XMyc reached roughly native Mlh3 levels 60 to 90 minutes following induction (Fig. 6B). Based on these two different β-estradiol induction protocols we estimated that functional Mlh3-13XMyc levels were achieved 60 to 90 minutes post β-estradiol induction.

We induced *MLH3* (untagged) expression with 10 nM β-estradiol at 0 to 6 hrs after inducing sporulation and monitored the effect of varying *MLH3* induction on crossing over (Fig. 6C, gray line). Inducing *MLH3* upon transfer to sporulation media (defined as T=0 hr) resulted in full complementation of crossing over at *CEN8-THR1* (39% TT), but a lack of β-estradiol induction (DMSO treatment) resulted in *mlh3*Δ levels (23% TT). While a drop in crossing over at later induction timepoints was expected, we were surprised by the slope of the curve. *MLH3* induction at 3 hrs post transfer to sporulation medium was sufficient to impair crossing over (34% TT), while inductions at 4, 5, and 6 hrs after initiating sporulation led to crossover rates comparable to no induction (25, 24, and 23 % TT, respectively). The 3 to 5 hr window was roughly 3 hrs before 50% of cells had completed at least the MI division (T=7 hrs), indicating that this window overlapped with early events in meiotic prophase such as SEI formation (Zakharyevich et al. 2012; see Discussion).

We hypothesized that Mlh1-Mlh3 recruitment in early meiotic prophase may not require its endonuclease activity. To address this, we performed β-estradiol inductions of *MLH3* in strains bearing the endonuclease dead *mlh3-D523N* allele under the control of its native promoter (Nishant et al. 2008; Fig. 6C, blue line). If Mlh3 was needed to perform an early structural role independent of its catalytic function, then constitutive expression of *mlh3-D523N* in combination with induced expression of *MLH3* at later time points would be sufficient to promote crossing over. Such a phenotype was not seen, instead, we observed a significant decrease in crossing over at the 2 hr *MLH3* induction timepoint in *mlh3-D523N* (33% TT) compared to *mlh3*Δ (38% TT). The dominant negative effect conferred by *mlh3-D523N* can be explained by a defective Mlh1-Mlh3 being loaded at early times in meiotic prophase and that it cannot be easily substituted by newly synthesized subunits. Such a model is supportive of an early role (timing of loading and possibly catalytic) for Mlh1-Mlh3 in meiotic prophase that has functional consequences for crossing over.

### *DMC1* haploinsufficiency phenotype is not seen in *dmc1-T159A* and is rescued by increased dosage of *mlh3* alleles that impact Mlh1-Mlh3 stability

A parsimonious model to explain our genetic and biochemical data is that Dmc1 filaments contribute to the nucleation of a Mlh1-Mlh3 polymer that is active in crossover resolution. This model is supported by the following observations: 1. One copy of *DMC1* is haploinsufficient for crossing over in backgrounds in which Mlh1-Mlh3 is destabilized (e.g., *mlh3-42* and *mlh3-54*). 2. The temporal window when *MLH3* must be expressed for its full pro-crossover functions overlaps with robust Dmc1 expression and Dmc1-Mlh3 physical interactions (see Discussion). To test this model, we measured *CEN8-THR1* crossover rates in *MLH3, mlh3Δ, mlh3-42* and *mlh3-54* strains containing one copy of the hypomorphic *dmc1-T159A* allele. dmc1-T159A protein is defective in Dmc1 monomer-monomer contacts and forms less stable presynaptic filaments. However, it can interact with the Rdh54 accessory factor and mediate strand exchange *in vitro*, though less efficiently than Dmc1 (Liu et al. 2014). DSBs at the *HIS4::LEU2* meiotic crossover hotspot appeared with timing similar to *wild-type* but persisted for longer periods of time and accumulated to higher levels. Crossing over at the *HIS4::LEU2* meiotic hotspot, while delayed by about 2 hrs, reached approximately *wild-type* levels, and sporulation efficiency and spore viability were similar to or close to *wild-type* (Liu et al. 2014).

We hypothesized that if Mlh1-Mlh3 was functioning to facilitate Dmc1 strand invasion, *dmc1-T159A mlh3-42* or *dmc1-T159A mlh3-54* double mutants might display more dramatic meiotic defects than *mlh3-42* or *mlh3-54* single mutants. Such a phenotype would resemble the more severe defect seen in the *hed1Δ dmc1-T159A* double mutant, which showed increased intersister recombination, reduced crossing over, and more severe delays in meiotic progression than the *dmc1-T159A* single mutant (Liu et al. 2014). Liu et al. (2014) suggest that the down-regulation of Rad51 activity in meiosis, which is dependent on *HED1* function, “promotes Dmc1-dependent interhomolog recombination.” Thus, the more severe meiotic defect seen in *hed1Δ dmc1-T159A* could be explained by dmc1-T159A strand exchange functions becoming less effective when Rad51 is up regulated. As shown in Fig. 7A, *dmc1-T159A* modestly elevated crossover rates at *CEN8-THR1* in all backgrounds tested. In *wild-type* and *mlh3*Δ, *dmc1-T159A* increased TTs by 6%; in *mlh3-42* and *mlh3-54*, TTs were *increased* by 9% and 10%, respectively (compare dark grey bars to orange bars). The *dmc1-T159A* meiotic progression delay was not exacerbated by *mlh3-42* or *mlh3-54,* suggesting that dmc1-T159A function was not further compromised by the *mlh3* alleles (Fig. 7B). These observations suggest that the crossover defect seen when *DMC1* was haploinsufficient in *mlh3-42* or *mlh3-54* was unlikely to be due to a defect that impacted Dmc1 strand invasion but could be better explained by a model in which Dmc1 nucleates Mlh1-Mlh3 polymers to act in post strand invasion steps. One possibility is that the meiotic progression delay seen in *dmc1-T159A* might benefit an unstable Mlh1-mlh3 complex by providing a longer time period to form polymers.

**Fig. 7.**
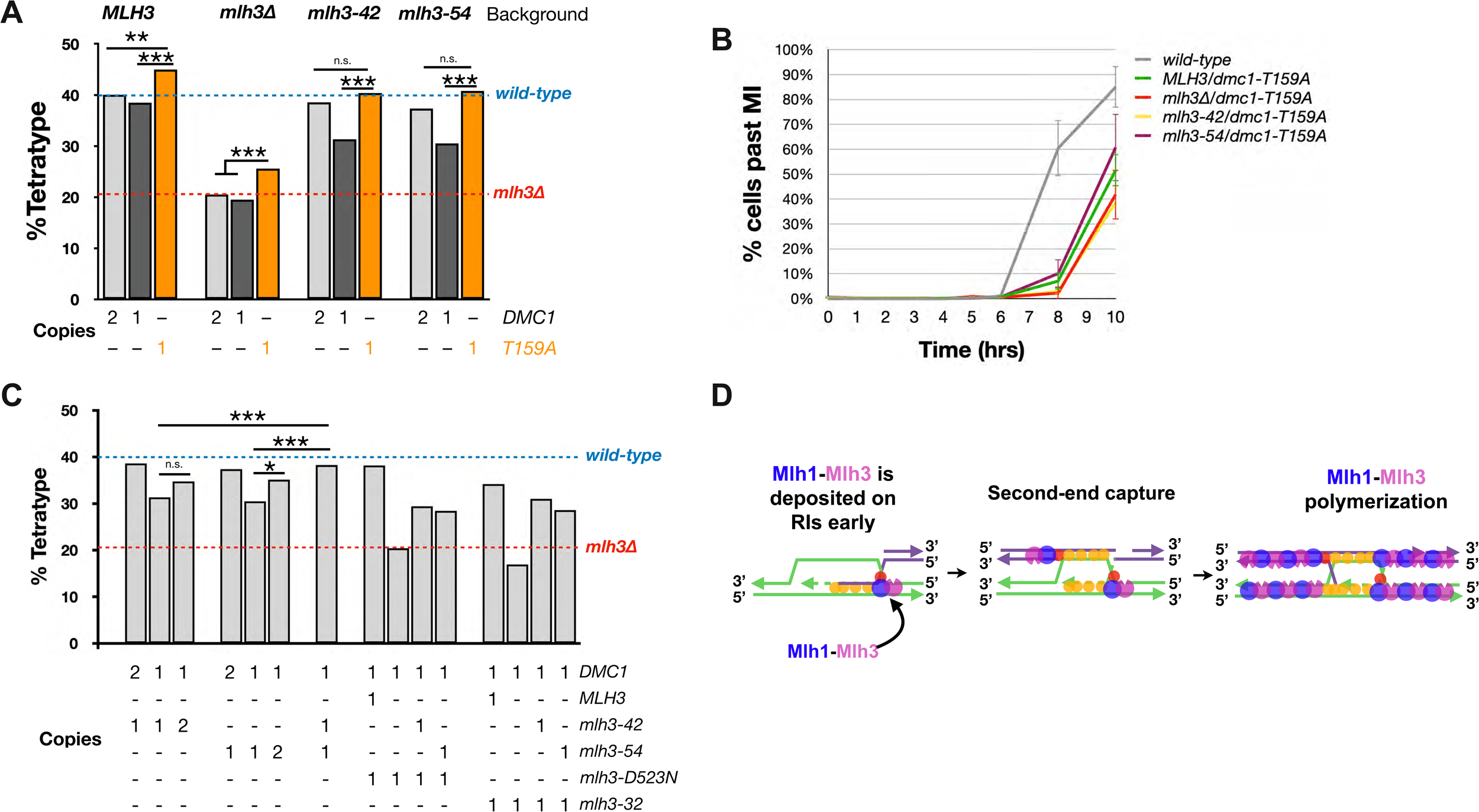
***DMC1* haploinsufficiency phenotype is rescued by *dmc1-T159A* and functional *MLH3* supplementation**. A. Crossover analysis at *CEN8-THR1* for the indicated *wild-type* and *dmc1-T159A* strains. B. Meiotic progression of strains shown in panel A. Error bars represent standard error. C. *CEN8-THR1* interval crossover data for strains containing 1 or 2 copies of *DMC1* in *mlh3-42, −54, -D523N* (endonuclease defective), and -*32* (crossover defective strains. D. A model proposing that retention of Dmc1 in meiotic recombination intermediates nucleates the formation of stable Mlh1-Mlh3 polymers required for dHJ resolution. Asterisks indicate significance (p>0.05 = n.s., p<0.05 = *, p<0.01 = **, and p<0.001 = ***).

If Dmc1 acts to nucleate Mlh1-Mlh3 polymer formation, then supplementation of additional *mlh3-42* and *mlh3-54* copies might rescue *DMC1* haploinsufficiency by stabilizing Mlh1-mlh3-42 and Mlh1-mlh3-54 complexes through mass action. To test this, we maintained *DMC1* in single copy and included additional copies of *mlh3-42* and *mlh3-54*. Two copies of either *mlh3-42* or *mlh3-54* modestly increased crossover rates in *DMC1* single copy backgrounds. When both *mlh3-42* and *mlh3-54* were expressed, crossing over was restored to *wild-type* (38%; Fig. 7C). Because *mlh3-42* and *mlh3-54* may disrupt heterodimerization through distinct mechanisms (Fig. 2), a mix of the two alleles might offset their individual defects.

To test if the *DMC1* haploinsufficiency rescue required increased dosage of meiotically functional Mlh3, we expressed *mlh3-D523N* (endo-dead) or *mlh3-32* (MMR^+^/CO^−^) in *mlh3-42* and *mlh3-54* backgrounds. The *mlh3-32* allele does not disrupt MMR and displays endonuclease activity that is stimulated by the Msh2-Msh3 MMR complex; however, it confers an *mlh3-null* like phenotype for meiotic crossing over (Al-Sweel et al. 2017). We found that neither crossover defective mutation could restore *wild-type* levels of crossing over (Fig. 7C). Given that mlh3-D523N and mlh3-32 are stable interactors of Mlh1 (Al-Sweel et al. 2017), they may outcompete mlh3-42 and mlh3-54 for binding to Mlh1, forming primarily non-functional polymers. We conclude that supplementation of only meiotically functional Mlh3 can restore crossing over when *DMC1* is haploinsufficient.

## Discussion

We were interested in identifying a meiotic network of Mlh3 interactors by performing a screen for genes haploinsufficient in *mlh3* backgrounds in which the stability of Mlh1-Mlh3 was predicted to be impaired (*mlh3-42* and *mlh3-54* strains). We screened 34 meiotic genes and discovered established Mlh1-Mlh3 interactors such as *EXO1, MSH5, PMS1,* and *RTK1,* a putative kinase of unknown function. Unexpectedly, one of the strongest genetic interactions involved the meiotic recombinase gene *DMC1*. We then showed that Mlh1-Mlh3 physically interacts with Dmc1 *in vitro* and *in vivo*. This link between “early” and “late” meiotic recombination factors may help contextualize various *MLH3* phenotypes which have been difficult to reconcile with a purely dHJ resolution role for Mlh1-Mlh3.

When is Mlh3 required for function in meiosis? We cautiously determined this by cross referencing the timing of the MI/MII divisions in our experiments with previous studies. In our time courses, 50% of cells complete the first meiotic division by ∼7 hrs (T=0 indicates transfer to sporulation media; Fig. 6D). This timing is very similar to that seen in several studies which analyzed recombination intermediates at the *HIS4::LEU2* recombination hotspot including meiotically induced DSBs, joint molecules, and crossovers (Cao et al. 1990; Schwacha and Kleckner 1994; Schwacha and Kleckner 1995; Kosaka et al. 2008; Wanat et al. 2008; Zakharyevich et al. 2012). In these studies, maximum DSBs were observed at ∼4-5 hrs, followed by peak SEI/JM/dHJ formation at ∼5 hrs. Crossovers reached 50% of their maximum level at ∼6 hrs. In our *MLH3*-induction time courses we estimated that functional Mlh3 levels occurred 60 to 90 minutes post β-estradiol induction and that *MLH3* induction as early as 3 hrs after transfer to sporulation media resulted in crossover defects. Assuming the appearance of recombination intermediates at *HIS4::LEU2* is similar to that seen at other loci (e.g. *CEN8-THR),* we infer that Mlh3 would be needed at roughly the time of DSB and JM formation in order to act in crossover resolution.

At what stage of meiotic recombination does Mlh3 interact with Dmc1? We found that meiotic progression was delayed by ∼2 hrs in our immunoprecipitation time courses (Fig. 5D) compared to the β-estradiol induction time courses (Fig. 6D). Accounting for this delay, we estimated that DSBs in the immunoprecipitation time courses would likely peak at 6 hrs and crossing over would reach 50% maximum levels by ∼8 hrs. We detected Mlh3-Dmc1 interactions at 6 and 8 hrs in these experiments, consistent with Mlh3 interacting with Dmc1 at stages when its expression is required for its strand exchange functions. The detection of robust interactions at 8 hrs suggests that Dmc1 may be retained at sites of recombination until crossovers form, providing a stable platform to nucleate Mlh1-Mlh3 polymer formation (Fig. 7D). More precise time courses that include the measurement of physical recombination intermediates will be necessary to confirm this idea, and ChIP and immunofluorescence experiments would likely be useful to determine if Mlh3 recruitment to recombination intermediates is dependent on Dmc1. Finally, because we have not obtained a mutation which disrupts the interaction between Mlh3 and Dmc1, our experiments do not rule out the possibility that other factors such as Mlh1 may be responsible for Mlh3 recruitment to Dmc1.

### Potential early functions for Mlh1-Mlh3

Little is known about the mechanism or purpose for an early Mlh1-Mlh3 role. One possibility is that early recruitment of Mlh1-Mlh3 ensures that it acts in the Class I crossover pathway as a functional polymer. However, another MLH complex, Mlh1-Mlh2, has been implicated in regulating both crossover and noncrossover associated gene conversion tracts in conjunction with the Mer3 helicase. Recent studies proposed that Mer3-Mlh1-Mlh2 limits gene conversion through the inhibition of the Pif1, a 5’-3’ DNA helicase known to stimulate Polδ strand displacement through its interaction with the DNA polymerase processivity clamp PCNA (Abdullah et al. 2004; Wilson et al. 2013; Buzovetsky et al. 2017; Duroc et al. 2017; Altmannova et al. 2023). In support of this model, Vernekar et al. (2021) found that Mer3 interacts with both Pif1 and Rfc1 *in vivo* and that the Mer3-Mlh1-Mlh2 complex blocks Polδ-mediated D-loop extension *in vitro*.

The phenotypes associated with *mlh3Δ* mutants in meiosis, including longer gene conversion tracts and effects on branch migration, are reminiscent of the effect of the *mlh2Δ* mutation on meiotic gene conversion tract length (Duroc et al. 2017; Al-Sweel et al. 2017; Marsolier-Kergoat et al. 2018). One possibility consistent with the above phenotypes is that Mlh1-Mlh3 limits gene conversion tracts in yeast by modulating Dmc1 activity which should be concentrated near free 3’ DNA ends (Brown et al. 2015; Hinch et al. 2019; Slotman et al. 2020; Fig. 7D). It is important to note that the *mlh3Δ* phenotypes seen in baker’s yeast contrast with work in mice suggesting an endonuclease-independent role for MLH3 in lengthening gene conversion tracts (Premkumar et al. 2023). Our results also raise the question of whether an early Mlh3 role requires enzymatic activity. In the filamentous fungus *Sordaria*, Storlazzi et al. (2010) uncovered a role for Mlh1 in the resolution of chromosome entanglements during zygotene, prior to crossover formation in pachytene, possibly suggesting an early nucleolytic role for Mlh1 and one of its binding partners (either Pms1 or Mlh3). In addition, recent work in mammalian cells showed that human MLH3 functions in mitotic DSB repair in a manner that is dependent on its nuclease activity but does not require MLH1 (Rahman et al. 2020; reviewed in Pannafino and Alani 2021).

Lastly, we note a correlation between recombinase (Dmc1 vs. Rad51) and resolvase choice for meiotic crossing over (Mlh1-Mlh3 vs. SSN). While budding yeast, plants, and mammals utilize Dmc1 and Mlh1-Mlh3, other organisms such as *S. pombe*, *D. melanogaster, C. elegans,* and *N. crassa* do not fit this paradigm (Hunter 2015). Instead, these organisms use Rad51 homologs to drive a homology search and dHJ resolution is accomplished by SSNs (Hunter 2015). Such a pattern is intriguing because it suggests that crossover resolution mechanisms are linked to the recombinase that is employed.

### Haploinsufficiency as a tool to identify factors that interact in meiotic prophase

Haploinsufficiency screens have been performed to study factors that act in chromosome segregation in mitosis and meiosis (Baetz et al. 2004) and for expression of sporulation specific genes (Bungard et al. 2004). These approaches identified new interactions that were confirmed in biochemical and genetic assays. Our limited haploinsufficiency screen identified both previously known interactors of Mlh3 (Exo1, Pms1, Rtk1, Msh5) and a new factor Dmc1, suggesting that a genome-wide screen, while labor intensive, would likely identify other components that interact with Mlh3 or other factors that act in crossing over.

## Data Availability

Strains and plasmids are available upon request. The authors affirm that all data necessary for confirming the conclusions of the article are present within the article, figures, tables, supplemental figures, and supplemental files. Supplementary Fig. 1 contains examples of haploinsufficiency analysis; Supplementary Fig. 2 displays the *RTK1* analysis; Supplementary Fig. 3 shows co-immunoprecipitation analysis of yMlh1-Mlh3 and yDmc1 in the presence of DNase I; Supplementary Fig. 4 displays the effect of *dmc1* mutations on meiotic crossing over in the *CEN8-THR1* interval; Supplementary Fig. 5 provides a summary of yeast two-hybrid analysis; Supplementary Fig. 6 shows mRNAseq profiling of *CLB1* and *MLH3* in SK1 meiosis and the effect of inducing *GAL1-MLH3* with estradiol. Supplementary Files 1, 2, and 3, present plasmids, oligonucleotides and strains used in this study, respectively; Supplementary File 4 displays the spore autonomous fluorescence assay data; Supplementary File 5 shows the raw data for the time-course phenotypic data; Supplementary File 6 shows the spore viability profiles; Supplementary File 7 displays the raw data for the Chromosome XV genetic map distance analysis of *wild-type* and *rtk1Δ* strains. Supplementary material is available at GENETICS online.

## Acknowledgments

We are grateful to Nancy Hollingsworth, Doug Bishop, Maureen Hanson, and Michael Sehorn for reagents and helpful advice for experiments performed in this study, Marcus Smolka and Doug Bishop for comments on the manuscript, and members of the Alani lab and Michael Lichten for helpful discussions.

## Funding

G.N.P., J.J.C., V.M., M.C.G. and E.A. were supported by the National Institute of General Medical Sciences of the National Institutes of Health (https://www.nih.gov/): R35GM134872. L.P. was funded by a Sloan Fellowship and by National Institutes of Health grant F31GM145163. J.B.C. was supported by National Institute of General Medical Sciences of the National Institutes of Health (https://www.nih.gov/): R35GM142457. The content of this work is solely the responsibility of the authors and does not necessarily represent the official views of the National Institutes of Health. The funders had no role in study design, data collection and analysis, decision to publish, or preparation of the manuscript.

## Conflicts of interest

None declared.

## Legends, Supplementary Figures

**Supplementary Fig. 1.**
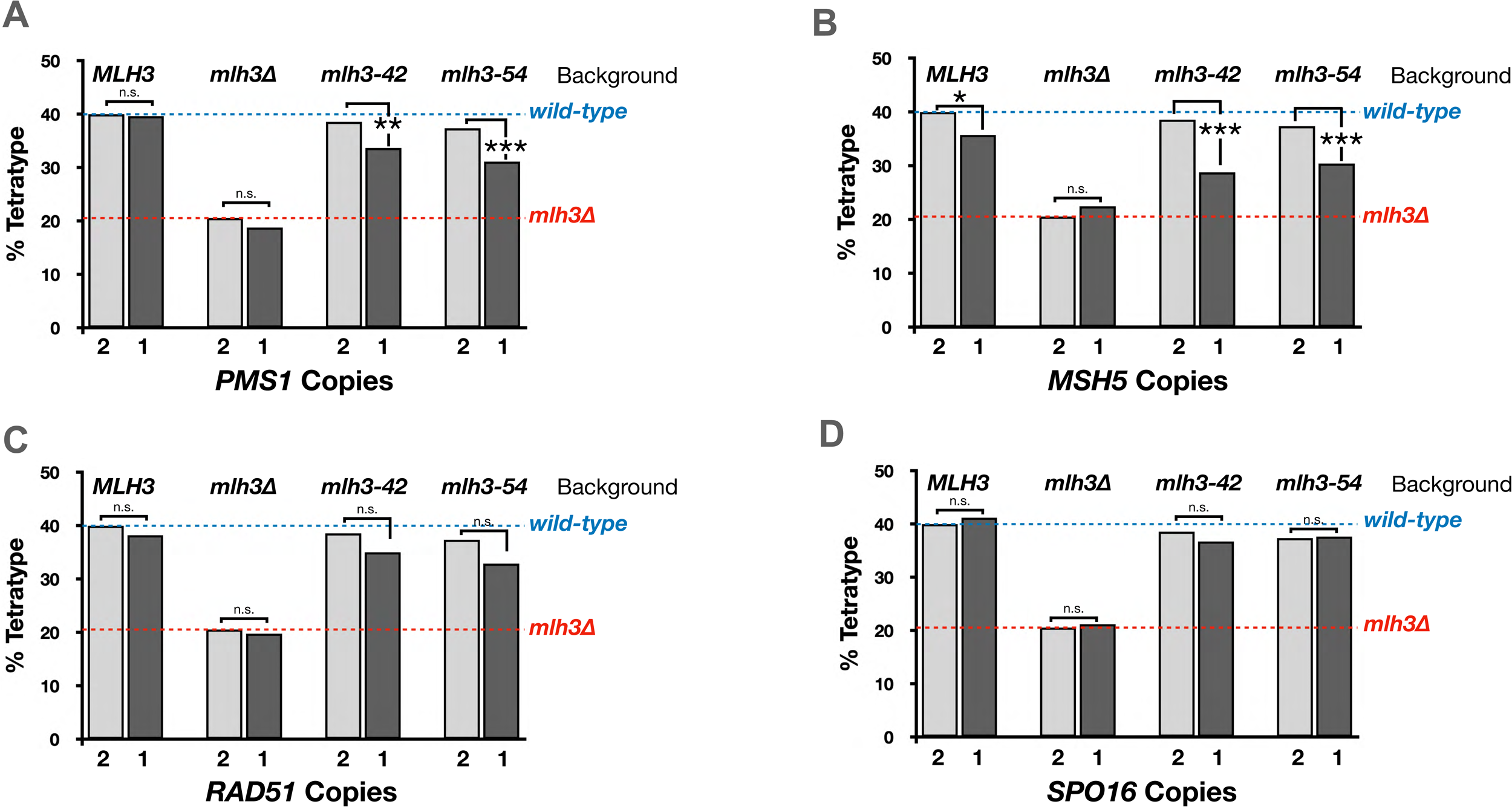
Examples of haploinsufficiency analysis. Strains containing one or two copies of 34 meiotic genes (Fig. 3A) were analyzed for crossing over at the *CEN8-THR1* interval in *wild-type*, *mlh3Δ* and *mlh3-42* and *mlh3-54* strains. Examples of the data shown in Supplementary File 4 are provided in bar graph form for *PMS1* (A), *MSH5* (B), *RAD51* (C) and *SPO16* (D). Significance was assessed by *χ*^2^ test between haplosufficient and haploinsufficient conditions. To minimize ***α*** inflation due to multiple comparisons, we applied a Benjamini-Hochberg correction at a 5% false discovery rate. Asterisks indicate significance (p>0.05 = n.s., p<0.05 = *, p<0.01 = **, and p<0.001 = ***).

**Supplementary Fig. 2.**
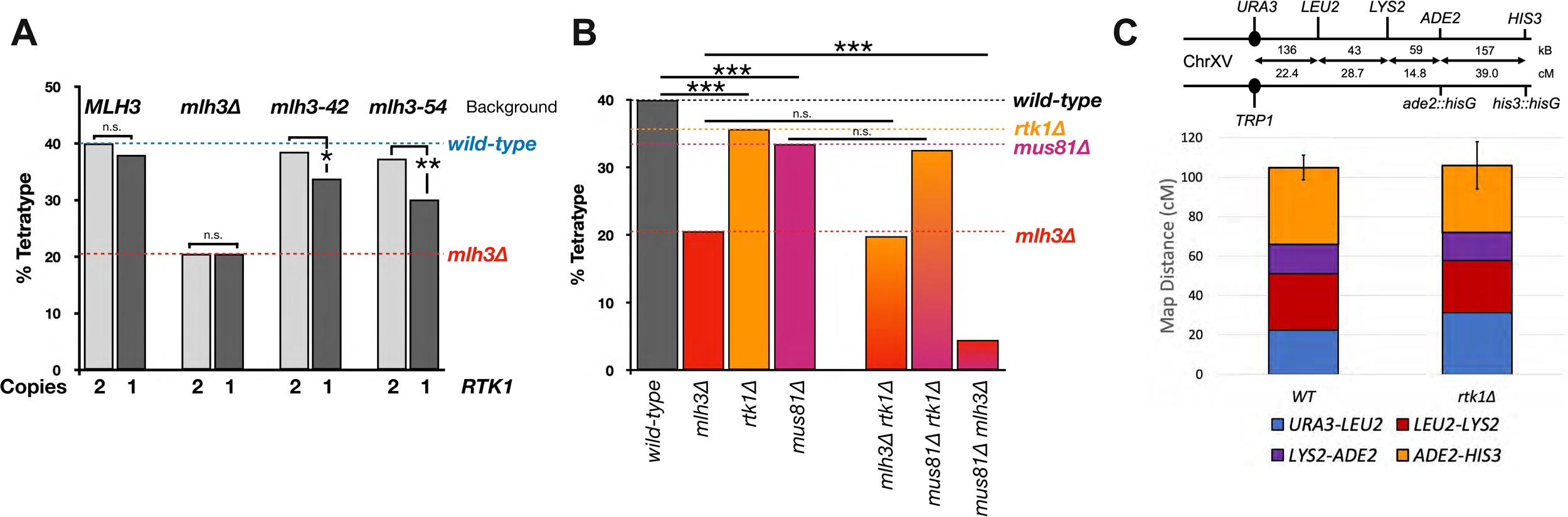
*RTK1* analysis. A. Crossover analysis in the *CEN8-THR1* interval on chromosome VIII expressed as percent tetratype with the indicated *RTK1* gene dosage in *wild-type, mlh3Δ*, *mlh3-42*, and *mlh3-54* strains. B. Crossover analysis in the *CEN8-THR1* interval for strains homozygous for the indicated deletion mutations. C. Analysis of crossing over in the chromosome XV *CENXV-HIS3* interval in *rtk1Δ* strains. Top panel: Cartoon showing the *CENXV-HIS3* interval. The solid circle indicates the centromere and distances between markers in KB and cM are shown. Bottom panel: Cumulative genetic distance (cM) in *wild-type* (WT) and *rtk1Δ* strains. Each bar is divided into sectors corresponding to genetic intervals in the *URA3-HIS3*, as measured from tetrads (T). Standard error was calculated for each interval using Stahl Online Tools (https://elizabethhousworth.com/StahlLabOnlineTools/). Error bars represent the cumulative standard error across all four intervals. Underlying data for panels A and B can be found in Supplementary File 4. Underlying data for panel C can be found in Supplementary File 7. Asterisks indicate significance (p>0.05 = n.s., p<0.05 = *, p<0.01 = **, and p<0.001 = ***). A Benjamini-Hochberg correction was applied to data in panel A to account for multiple comparisons (5% false discovery rate). *Analysis of Supplementary Fig. 2. RTK1* encodes a putative kinase of unknown function and physically interacts Mlh1-Mlh3 in meiotic cells depleted of the anaphase-promoting complex activator Cdc20 (*P_CLB2_-CDC20*) which arrest at Metaphase I (Wild et al. 2019). The haploinsufficiency phenotype displayed by *RTK1* encouraged us to further probe the role of *RTK1* in meiosis (Panel A). First, we generated *rtk1*Δ diploids in the SKY3575/SKY3576 background to examine crossover rates on chromosome VIII. We observed a small reduction in crossing over (36% TT) in the absence of *RTK1*. This reduction, while distinct from *wild-type* (40% TT), is mild when compared to the crossover defect observed in Class I crossover mutants (∼20% TT). We then generated *rtk1Δ mlh3*Δ and *rtk1Δ mus81*Δ double mutants to test if *RTK1* was epistatic to Class I or Class II crossovers. We found that *rtk1Δ mlh3*Δ double mutants displayed a crossover rate of 20% TT, similar to *mlh3*Δ (21% TT). Surprisingly, *rtk1Δ mus81*Δ phenocopied *mus81*Δ, with a 33% TT value displayed by both mutants. This phenotype was in contrast to the dramatic decrease in crossing over seen in the *mus81Δ mlh3*Δ double mutant (4% TT), in agreement with previous studies which examined crossing over on Chromosome XV (Panel B; Argueso et al. 2004; Nishant et al. 2008; Sonntag Brown et al. 2013). To confirm that the lack of a strong crossover defect in *rtk1*Δ was not locus specific, we tested crossing over using tetrad analysis to track the segregation of markers over four consecutive intervals on Chromosome XV (104.9 cM in *wild-type*). As shown in Panel C, *rtk1*Δ resulted in a cumulative genetic distance of 106 cM with a spore viability of 97.5% (Supplementary Files 6, 7). We concluded that Rtk1 is not essential for either Class I or Class II crossovers and but may function to partially stabilize Mlh1-Mlh3. Argueso JL, Wanat J, Gemici Z, Alani E. 2004. Competing crossover pathways act during meiosis in *Saccharomyces cerevisiae*. Genetics. 168:1805–1816. Nishant KT, Plys AJ, Alani E. 2008. A mutation in the putative MLH3 endonuclease domain confers a defect in both mismatch repair and meiosis in *Saccharomyces cerevisiae*. Genetics. 179:747-755. Sonntag Brown M, Lim E, Chen C, Nishant KT, Alani E. 2013. Genetic analysis of *mlh3* mutations reveals interactions between crossover promoting factors during meiosis in baker’s yeast. G3. 3:9-22. Wild P, Susperregui A, Piazza I, Dorig C, Oke A, Arter M, et al. 2019. Network Rewiring of Homologous Recombination enzymes during mitotic proliferation and meiosis. Mol Cell. 22:859-874.e4. doi:10.1016/j.molcel.2019.06.022.

**Supplementary Fig. 3.**
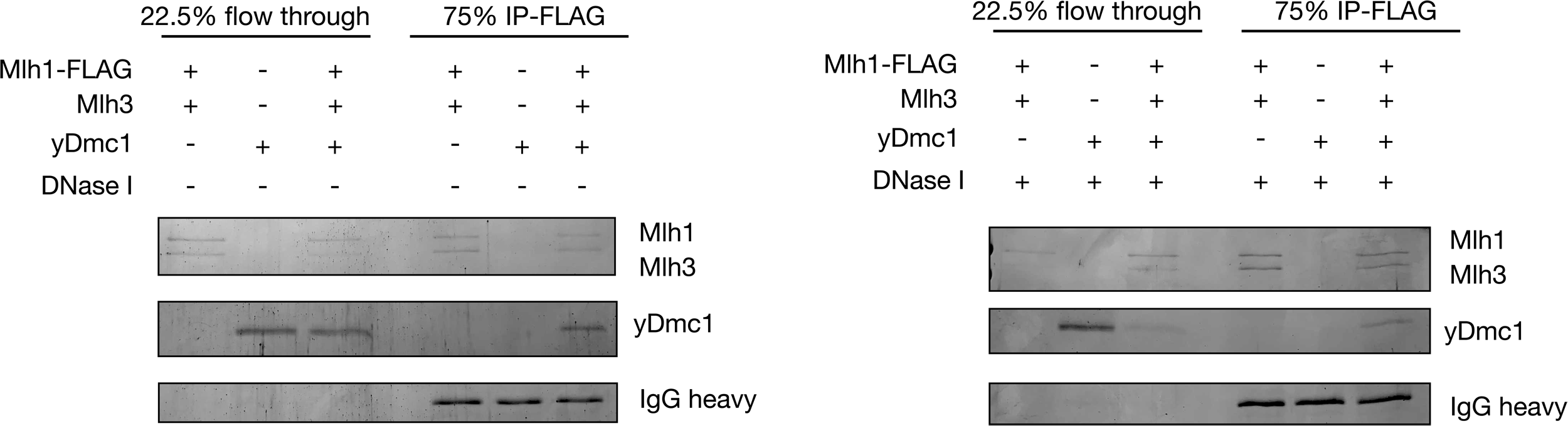
yMlh1-Mlh3 Co-IPs with yDmc1 in the presence and absence of DNase I. *In vitro* co-immunoprecipitation reactions (see Fig. 4 and Materials and methods) performed with Dynabeads Protein G conjugated to anti-FLAG M2 antibody and yMlh1-FLAG-Mlh3 and yDmc1 in the absence and presence of DNase I. Proteins were electrophoresed in 10% SDS-PAGE.

**Supplementary Fig. 4.**
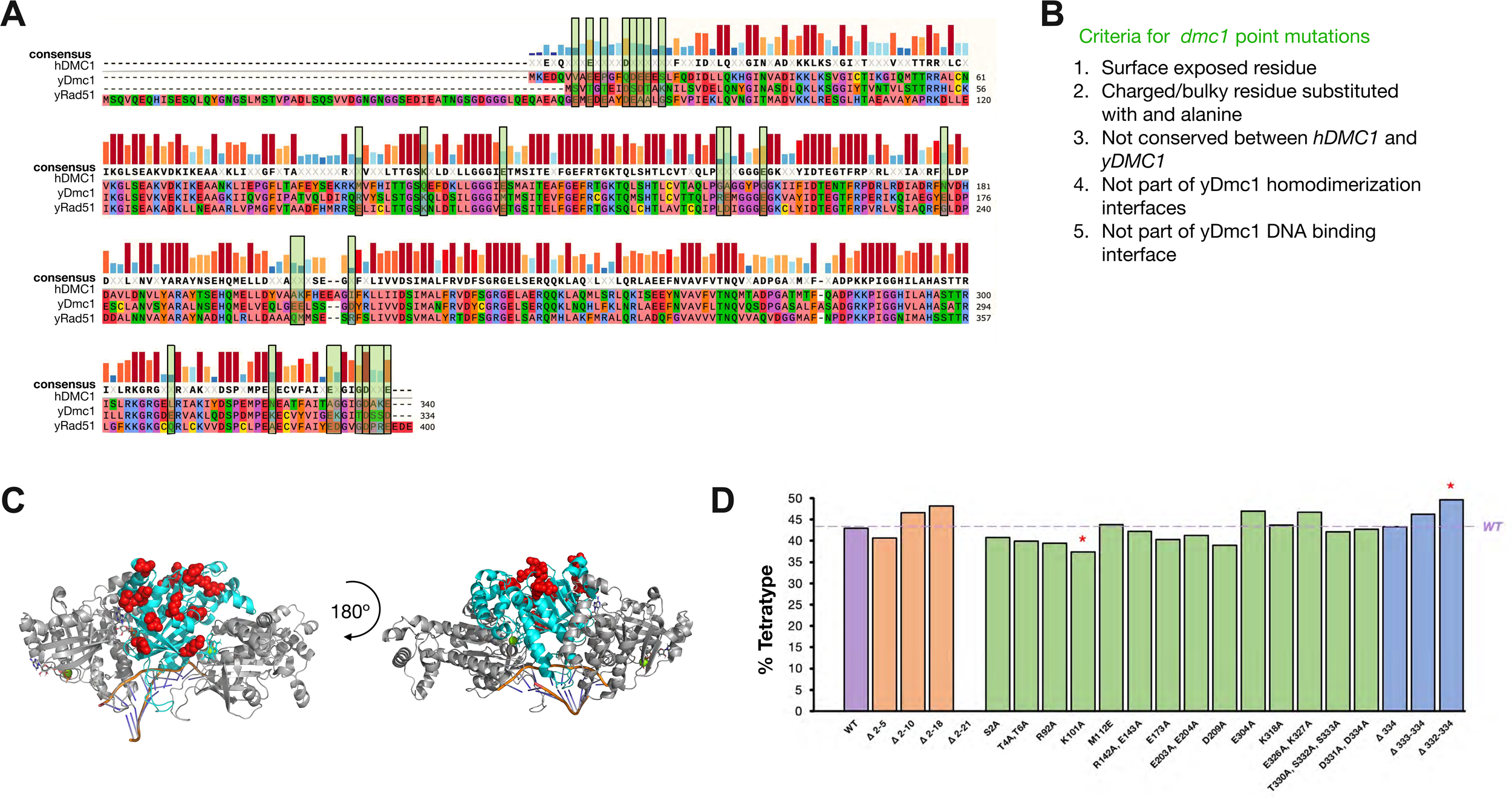
Effect of *dmc1* mutations on meiotic crossing over in the *CEN8-THR1* interval. A. MUSCLE (https://www.ebi.ac.uk/Tools/msa/muscle/) alignments of yRad51, yDmc1 and human DMC1 generated through SnapGene (https://www.snapgene.com/). B. We made mutations in Dmc1 using the following criteria: We first aligned the yDmc1, hDMC1 and yRad51 amino acids using the multiple sequence alignment tool in Snapgene. We then located charged/interesting yDmc1 unique residues and mapped them to the yDmc1 postsynaptic complex (PDB: 7EJ7) to confirm that they are surface exposed, and are not part of yDmc1 polymerization interface, not part of the yDmc1 DNA binding interface, and not a previously characterized mutant (Steinfeld et al. 2019). C. Mutated yDmc1 residues (red spheres) mapped to middle subunit (cyan) of trimeric yDmc1 postsynaptic complex (PDB: 7EJ7). D. Crossover analysis in the *CEN8-THR1* interval for strains bearing the indicated *dmc1* alleles. Steinfeld JB, Beláň O, Kwon Y, Terakawa T, Al-Zain A, Smith MJ, Crickard JB, Qi Z, Zhao W, Rothstein R, Symington LS, Sung P, Boulton SJ, Greene EC. 2019. Defining the influence of Rad51 and Dmc1 lineage-specific amino acids on genetic recombination. Genes Dev. 33:1191-1207.

**Supplementary Fig. 5.**
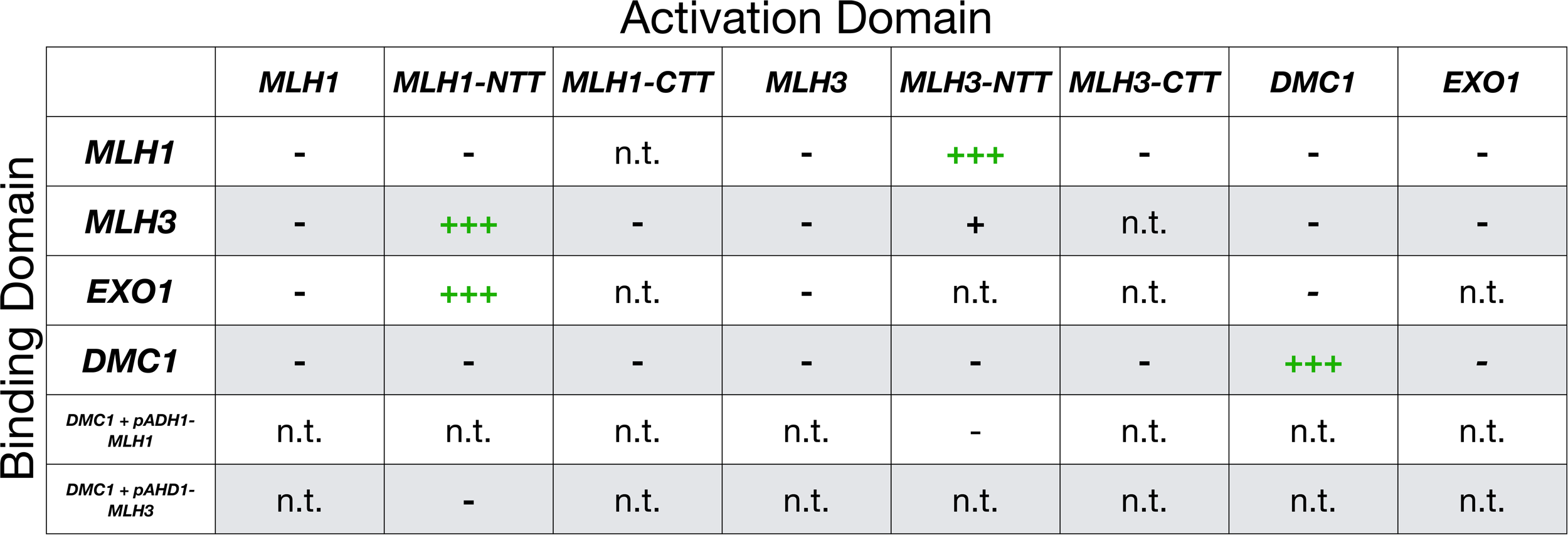
Yeast two-hybrid analysis. Two-hybrid interactions involving the indicated Gal4 DNA binding and Gal4 activation domains fused to Mlh1, Mlh3, Exo1, Dmc1, and N- and C-terminal domains of Mlh1 and Mlh3. See Materials and methods for details. When indicated, a 2μ vector expressing *MLH1* or *MLH3* driven by the *ADH1* promoter was included in the two-hybrid strain. Cells containing the indicated plasmids were spotted on -leucine, -tryptophan plates to assess plating efficiency and -leucine, - tryptophan, -histidine, -adenine plates to assess growth after 3 days of incubation at 30.C. n.t.=not tested, +=weak growth, +++=strong growth on -leucine, -tryptophan, -histidine, -adenine plates. *Analysis of Supplementary Fig. 5*. We confirmed Dmc1’s self-interaction (Dresser et al. 1997), and by constructing deletions of the N-termini (containing the ATP binding domain) of either Mlh1 (*mlh1-*Δ*1-344*) or Mlh3 (*mlh3-*Δ*1-374*) were able to detect an interaction between Mlh1 and Mlh3. This observation is consistent with previous studies which showed that two-hybrid interactions between Mlh1 and Mlh3 required truncated gene products (Wang et al. 1999; Al-Sweel et al. 2017). We note that it was necessary to truncate Mlh1 to detect an Mlh1-Exo1 two-hybrid interaction. We detected a weak interaction between mlh3-Δ1-374 and full length Mlh3. The possibility that Mlh3 may self-interact was not seen through meiotic Co-IP (Sanchez et al. 2020) but has indirect support from other studies (Kolas et al. 2005, Rahman et al. 2020, reviewed in Pannafino and Alani 2021). We did not detect any interactions between Dmc1 and either Mlh1 or Mlh3, even when co-overexpressing the complementary MLH subunit to restore a balanced stoichiometry. Dresser ME, Ewing DJ, Conrad MN, Dominguez AM, Barstead R, Jiang H, Kodadek T. 1997. DMC1 functions in a *Saccharomyces cerevisiae* meiotic pathway that is largely independent of the RAD51 pathway. Genetics. 147:533-44. doi: 10.1093/genetics/147.2.533. Kolas NK, Svetlanov A, Lenzi ML, Macaluso FP, Lipkin SM, Liskay RM, Greally J, Edelmann W, Cohen PE. 2005.Localization of MMR proteins on meiotic chromosomes in mice indicates distinct functions during prophase I. J Cell Biol. 171:447-458. Pannafino G, Alani E. 2021. Coordinated and Independent Roles for MLH Subunits in DNA Repair. Cells. 10:948. doi: 10.3390/cells10040948 Rahman MM, Mohiuddin M, Shamima Keka I, Yamada K, Tsuda M, Sasanuma H, Andreani J, Guerois R, Borde V, Charbonnier JB, Takeda S. 2020. Genetic evidence for the involvement of mismatch repair proteins, PMS2 and MLH3, in a late step of homologous recombination. J Biol Chem. 295:17460-17475. doi: 10.1074/jbc.RA120.013521.

**Supplementary Fig. 6.**
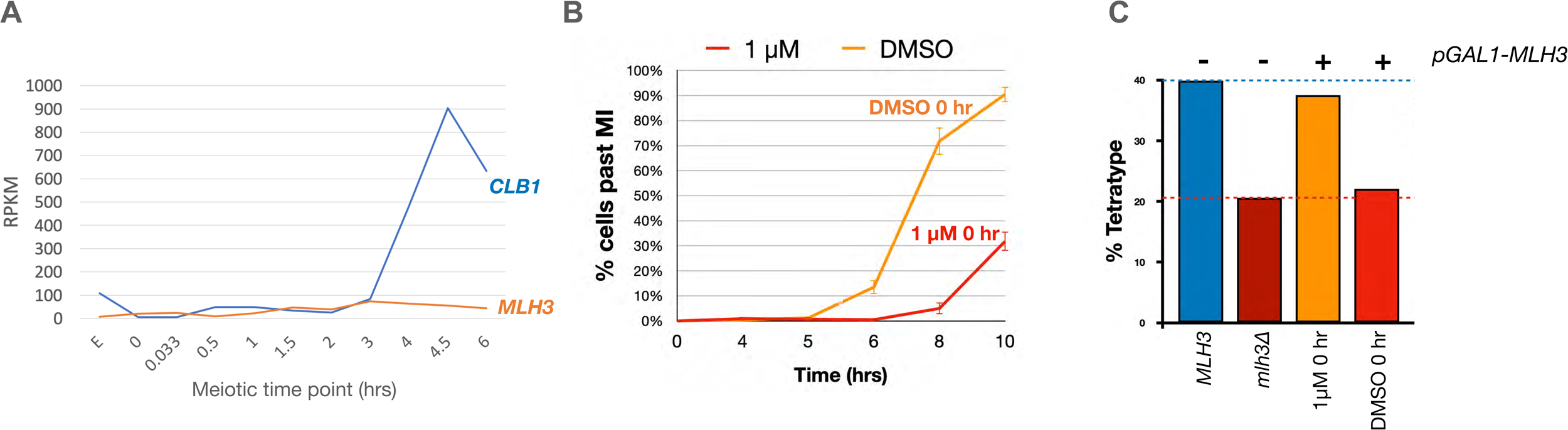
A. mRNAseq profiling of *CLB1* and *MLH3* in SK1 meiosis. Data obtained from Brar et al. (2012). RPKM= Reads per kilobase of coding sequence per million mapped reads. B., C. *GAL1-MLH3* estradiol induced expression (1 μM) at the time of transfer to sporulation media did not impact crossing over in the *CEN8-THR1* interval (B) but resulted in significant delays in completing the Meiosis I division (C). In C, the 0 μM estradiol graph (DMSO control) shows the average of four experiments; the 1 μM estradiol plot shows the average of two experiments. +/−standard error of the mean (SEM). Brar GA, Yassour M, Friedman N, Regev A, Ingolia NT, Weissman JS. 2012. High-resolution view of the yeast meiotic program revealed by ribosome profiling. Science. 335:552-557.

**Supplementary File 1.**
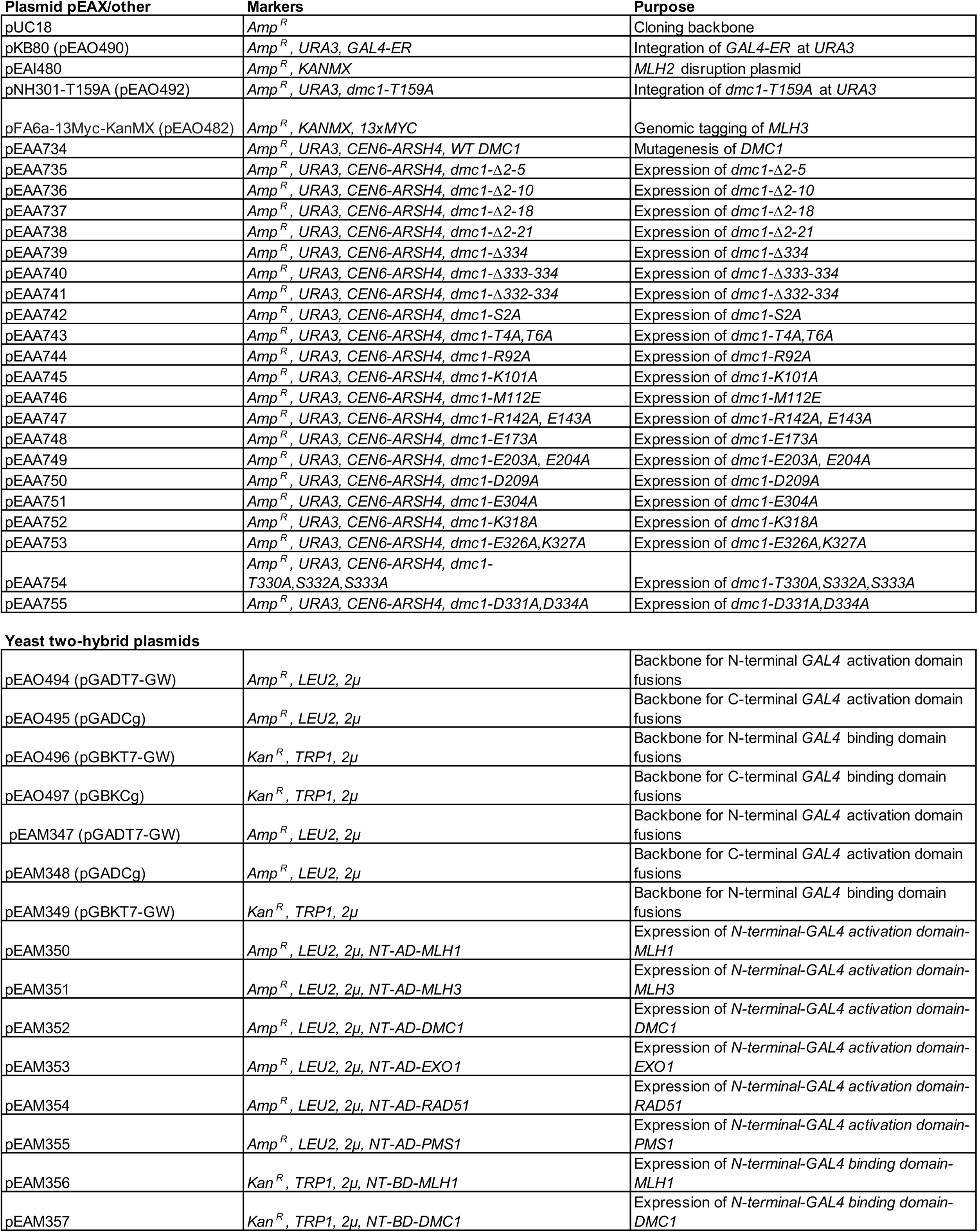
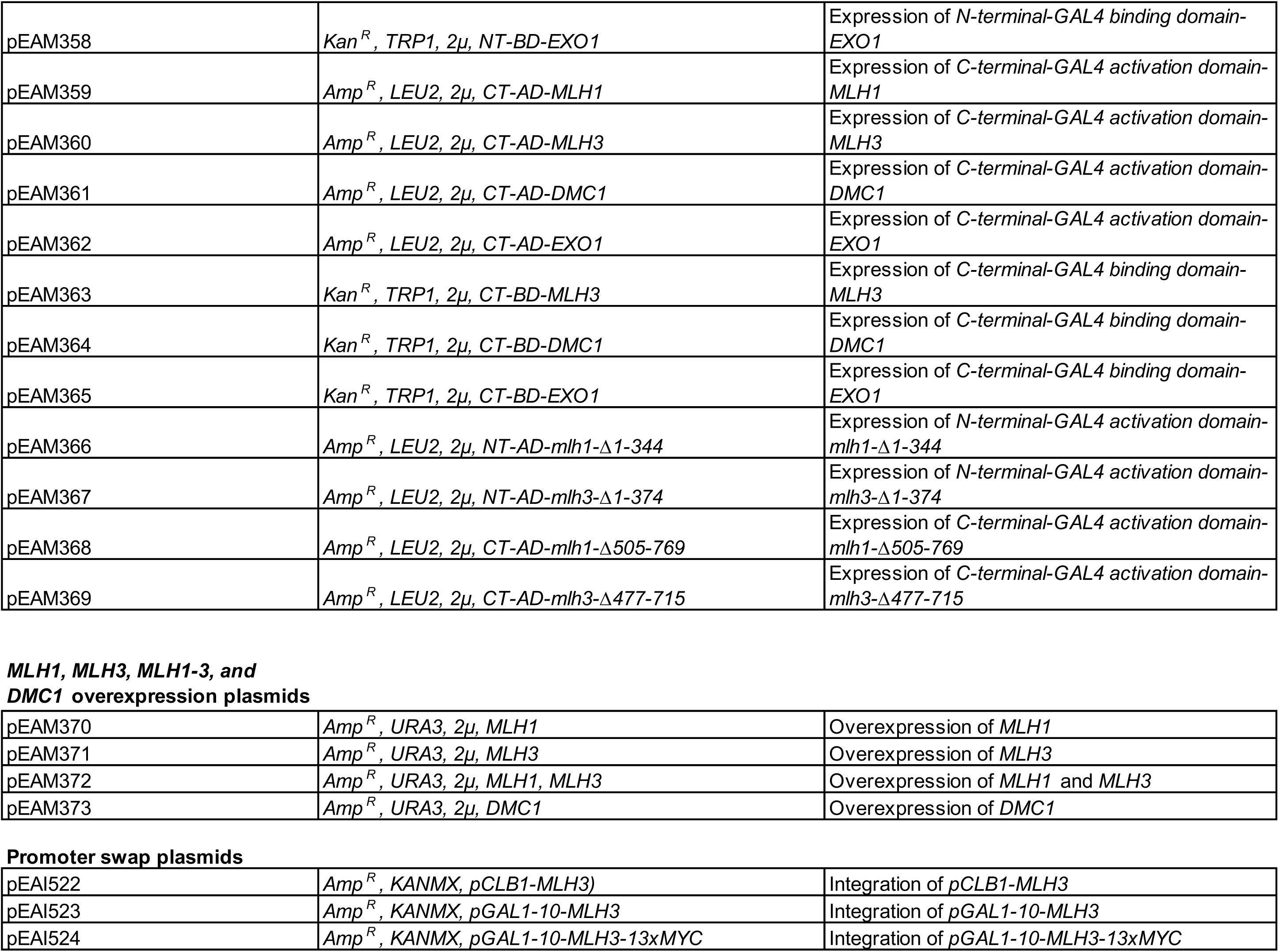
Plasmids used in this study.

| Plasmid pEAX/other | Markers | Purpose |
| --- | --- | --- |
| pUC18 | *Amp<sup>R</sup>* | Cloning backbone |
| pKB80 (pEAO490) | *Amp<sup>R</sup>, URA3, GAL4-ER* | Integration of *GAL4-ER* at *URA3* |
| pEAI480 | *Amp<sup>R</sup>, KANMX* | *MLH2* disruption plasmid |
| pNH301-T159A (pEAO492) | *Amp<sup>R</sup>, URA3, dmc1-T159A* | Integration of *dmc1-T159A* at *URA3* |
| pFA6a-13Myc-KanMX (pEAO482) | *Amp<sup>R</sup>, KANMX, 13xMYC* | Genomic tagging of *MLH3* |
| pEAA734 | *Amp<sup>R</sup>, URA3, CEN6-ARSH4, WT DMC1* | Mutagenesis of *DMC1* |
| pEAA735 | *Amp<sup>R</sup>, URA3, CEN6-ARSH4, dmc1-Δ2-5* | Expression of *dmc1-Δ2-5* |
| pEAA736 | *Amp<sup>R</sup>, URA3, CEN6-ARSH4, dmc1-Δ2-10* | Expression of *dmc1-Δ2-10* |
| pEAA737 | *Amp<sup>R</sup>, URA3, CEN6-ARSH4, dmc1-Δ2-18* | Expression of *dmc1-Δ2-18* |
| pEAA738 | *Amp<sup>R</sup>, URA3, CEN6-ARSH4, dmc1-Δ2-21* | Expression of *dmc1-Δ2-21* |
| pEAA739 | *Amp<sup>R</sup>, URA3, CEN6-ARSH4, dmc1-Δ334* | Expression of *dmc1-Δ334* |
| pEAA740 | *Amp<sup>R</sup>, URA3, CEN6-ARSH4, dmc1-Δ333-334* | Expression of *dmc1-Δ333-334* |
| pEAA741 | *Amp<sup>R</sup>, URA3, CEN6-ARSH4, dmc1-Δ332-334* | Expression of *dmc1-Δ332-334* |
| pEAA742 | *Amp<sup>R</sup>, URA3, CEN6-ARSH4, dmc1-S2A* | Expression of *dmc1-S2A* |
| pEAA743 | *Amp<sup>R</sup>, URA3, CEN6-ARSH4, dmc1-T4A,T6A* | Expression of *dmc1-T4A,T6A* |
| pEAA744 | *Amp<sup>R</sup>, URA3, CEN6-ARSH4, dmc1-R92A* | Expression of *dmc1-R92A* |
| pEAA745 | *Amp<sup>R</sup>, URA3, CEN6-ARSH4, dmc1-K101A* | Expression of *dmc1-K101A* |
| pEAA746 | *Amp<sup>R</sup>, URA3, CEN6-ARSH4, dmc1-M112E* | Expression of *dmc1-M112E* |
| pEAA747 | *Amp<sup>R</sup>, URA3, CEN6-ARSH4, dmc1-R142A, E143A* | Expression of *dmc1-R142A, E143A* |
| pEAA748 | *Amp<sup>R</sup>, URA3, CEN6-ARSH4, dmc1-E173A* | Expression of *dmc1-E173A* |
| pEAA749 | *Amp<sup>R</sup>, URA3, CEN6-ARSH4, dmc1-E203A, E204A* | Expression of *dmc1-E203A, E204A* |
| pEAA750 | *Amp<sup>R</sup>, URA3, CEN6-ARSH4, dmc1-D209A* | Expression of *dmc1-D209A* |
| pEAA751 | *Amp<sup>R</sup>, URA3, CEN6-ARSH4, dmc1-E304A* | Expression of *dmc1-E304A* |
| pEAA752 | *Amp<sup>R</sup>, URA3, CEN6-ARSH4, dmc1-K318A* | Expression of *dmc1-K318A* |
| pEAA753 | *Amp<sup>R</sup>, URA3, CEN6-ARSH4, dmc1-E326A,K327A* | Expression of *dmc1-E326A,K327A* |
| pEAA754 | *Amp<sup>R</sup>, URA3, CEN6-ARSH4, dmc1-T330A,S332A,S333A* | Expression of *dmc1-T330A,S332A,S333A* |
| pEAA755 | *Amp<sup>R</sup>, URA3, CEN6-ARSH4, dmc1-D331A,D334A* | Expression of *dmc1-D331A,D334A* |
| Yeast two-hybrid plasmids |  |  |
| pEAO494 (pGADT7-GW) | *Amp<sup>R</sup>, LEU2, 2μ* | Backbone for N-terminal *GAL4* activation domain fusions |
| pEAO495 (pGADCg) | *Amp<sup>R</sup>, LEU2, 2μ* | Backbone for C-terminal *GAL4* activation domain fusions |
| pEAO496 (pGBKT7-GW) | *Kan<sup>R</sup>, TRP1, 2μ* | Backbone for N-terminal *GAL4* binding domain fusions |
| pEAO497 (pGBKCg) | *Kan<sup>R</sup>, TRP1, 2μ* | Backbone for C-terminal *GAL4* binding domain fusions |
| pEAM347 (pGADT7-GW) | *Amp<sup>R</sup>, LEU2, 2μ* | Backbone for N-terminal *GAL4* activation domain fusions |
| pEAM348 (pGADCg) | *Amp<sup>R</sup>, LEU2, 2μ* | Backbone for C-terminal *GAL4* activation domain fusions |
| pEAM349 (pGBKT7-GW) | *Kan<sup>R</sup>, TRP1, 2μ* | Backbone for N-terminal *GAL4* binding domain fusions |
| pEAM350 | *Amp<sup>R</sup>, LEU2, 2μ, NT-AD-MLH1* | Expression of *N-terminal-GAL4 activation domain-MLH1* |
| pEAM351 | *Amp<sup>R</sup>, LEU2, 2μ, NT-AD-MLH3* | Expression of *N-terminal-GAL4 activation domain-MLH3* |
| pEAM352 | *Amp<sup>R</sup>, LEU2, 2μ, NT-AD-DMC1* | Expression of *N-terminal-GAL4 activation domain-DMC1* |
| pEAM353 | *Amp<sup>R</sup>, LEU2, 2μ, NT-AD-EXO1* | Expression of *N-terminal-GAL4 activation domain-EXO1* |
| pEAM354 | *Amp<sup>R</sup>, LEU2, 2μ, NT-AD-RAD51* | Expression of *N-terminal-GAL4 activation domain-RAD51* |
| pEAM355 | *Amp<sup>R</sup>, LEU2, 2μ, NT-AD-PMS1* | Expression of *N-terminal-GAL4 activation domain-PMS1* |
| pEAM356 | *Kan<sup>R</sup>, TRP1, 2μ, NT-BD-MLH1* | Expression of *N-terminal-GAL4 binding domain-MLH1* |
| pEAM357 | *Kan<sup>R</sup>, TRP1, 2μ, NT-BD-DMC1* | Expression of *N-terminal-GAL4 binding domain-DMC1* |
| pEAM358 | *Kan<sup>R</sup>, TRP1, 2μ, NT-BD-EXO1* | Expression of *N-terminal-GAL4 binding domain-EXO1* |
| pEAM359 | *Amp<sup>R</sup>, LEU2, 2μ, CT-AD-MLH1* | Expression of *C-terminal-GAL4 activation domain-MLH1* |
| pEAM360 | *Amp<sup>R</sup>, LEU2, 2μ, CT-AD-MLH3* | Expression of *C-terminal-GAL4 activation domain-MLH3* |
| pEAM361 | *Amp<sup>R</sup>, LEU2, 2μ, CT-AD-DMC1* | Expression of *C-terminal-GAL4 activation domain-DMC1* |
| pEAM362 | *Amp<sup>R</sup>, LEU2, 2μ, CT-AD-EXO1* | Expression of *C-terminal-GAL4 activation domain-EXO1* |
| pEAM363 | *Kan<sup>R</sup>, TRP1, 2μ, CT-BD-MLH3* | Expression of *C-terminal-GAL4 binding domain-MLH3* |
| pEAM364 | *Kan<sup>R</sup>, TRP1, 2μ, CT-BD-DMC1* | Expression of *C-terminal-GAL4 binding domain-DMC1* |
| pEAM365 | *Kan<sup>R</sup>, TRP1, 2μ, CT-BD-EXO1* | Expression of *C-terminal-GAL4 binding domain-EXO1* |
| pEAM366 | *Amp<sup>R</sup>, LEU2, 2μ, NT-AD-mlh1-Δ1-344* | Expression of *N-terminal-GAL4 activation domain-mlh1-Δ1-344* |
| pEAM367 | *Amp<sup>R</sup>, LEU2, 2μ, NT-AD-mlh3-Δ1-374* | Expression of *N-terminal-GAL4 activation domain-mlh3-Δ1-374* |
| pEAM368 | *Amp<sup>R</sup>, LEU2, 2μ, CT-AD-mlh1-Δ505-769* | Expression of *C-terminal-GAL4 activation domain-mlh1-Δ505-769* |
| pEAM369 | *Amp<sup>R</sup>, LEU2, 2μ, CT-AD-mlh3-Δ477-715* | Expression of *C-terminal-GAL4 activation domain-mlh3-Δ477-715* |
| pEAM370 | *Amp<sup>R</sup>, URA3, 2μ, MLH1* | Overexpression of *MLH1* |
| pEAM371 | *Amp<sup>R</sup>, URA3, 2μ, MLH3* | Overexpression of *MLH3* |
| pEAM372 | *Amp<sup>R</sup>, URA3, 2μ, MLH1, MLH3* | Overexpression of *MLH1* and *MLH3* |
| pEAM373 | *Amp<sup>R</sup>, URA3, 2μ, DMC1* | Overexpression of *DMC1* |
| **Promoter swap plasmids** |  |  |
| pEAI522 | *Amp<sup>R</sup>, KANMX, pCLB1-MLH3)* | Integration of *pCLB1-MLH3* |
| pEAI523 | *Amp<sup>R</sup>, KANMX, pGAL1-10-MLH3* | Integration of *pGAL1-10-MLH3* |
| pEAI524 | *Amp<sup>R</sup>, KANMX, pGAL1-10-MLH3-13xMYC* | Integration of *pGAL1-10-MLH3-13xMYC* |

**Supplementary File 2.**
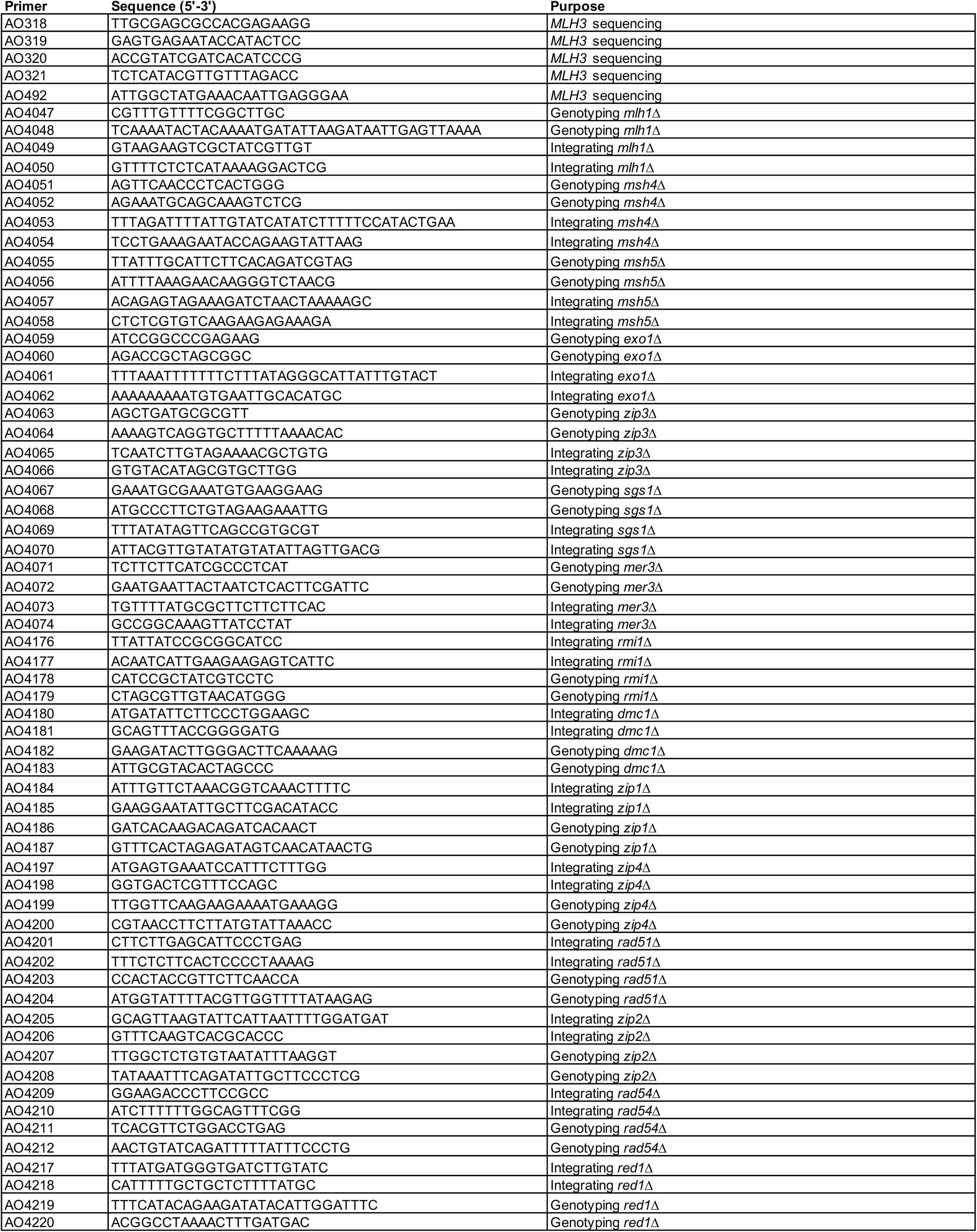
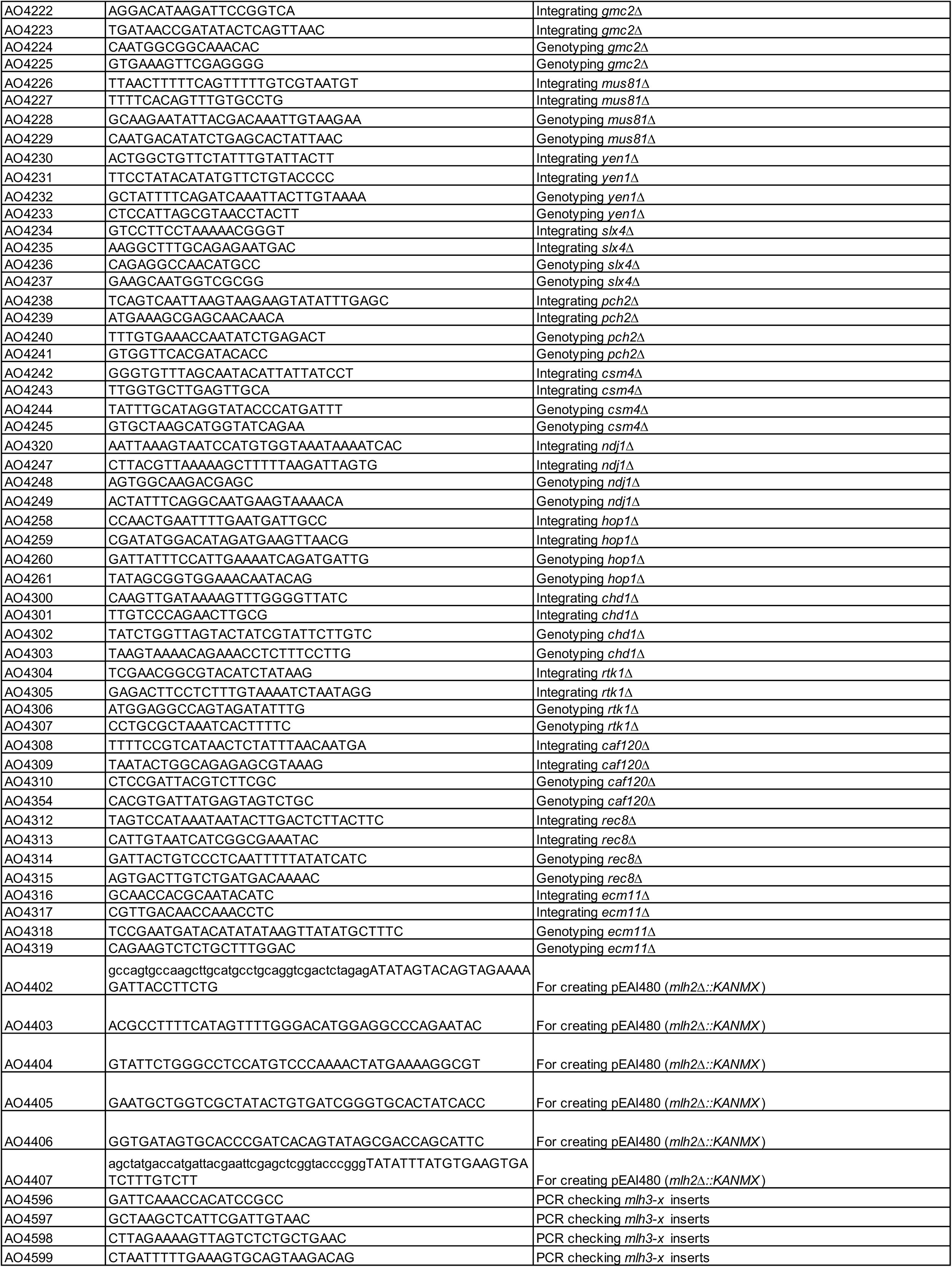
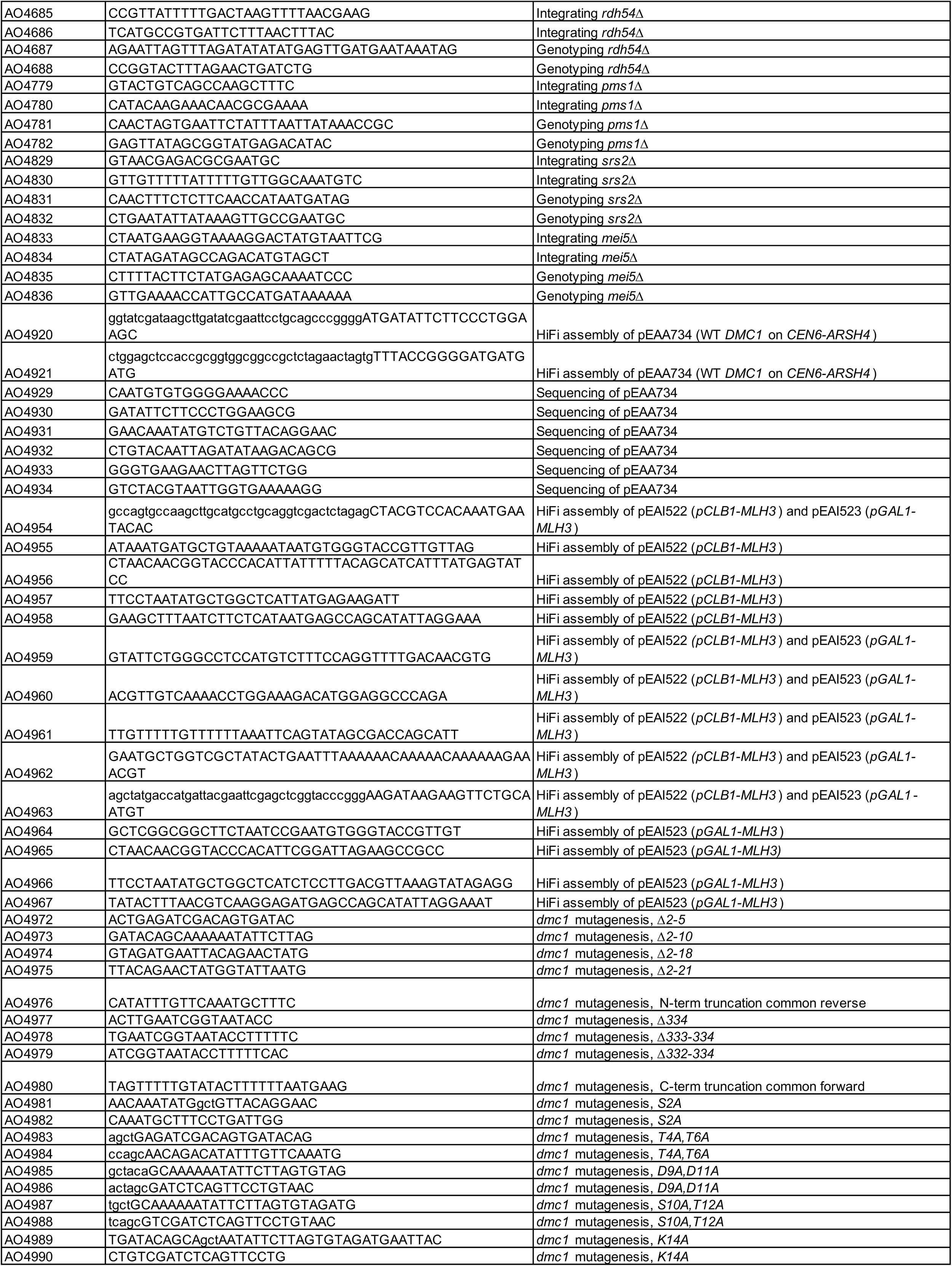
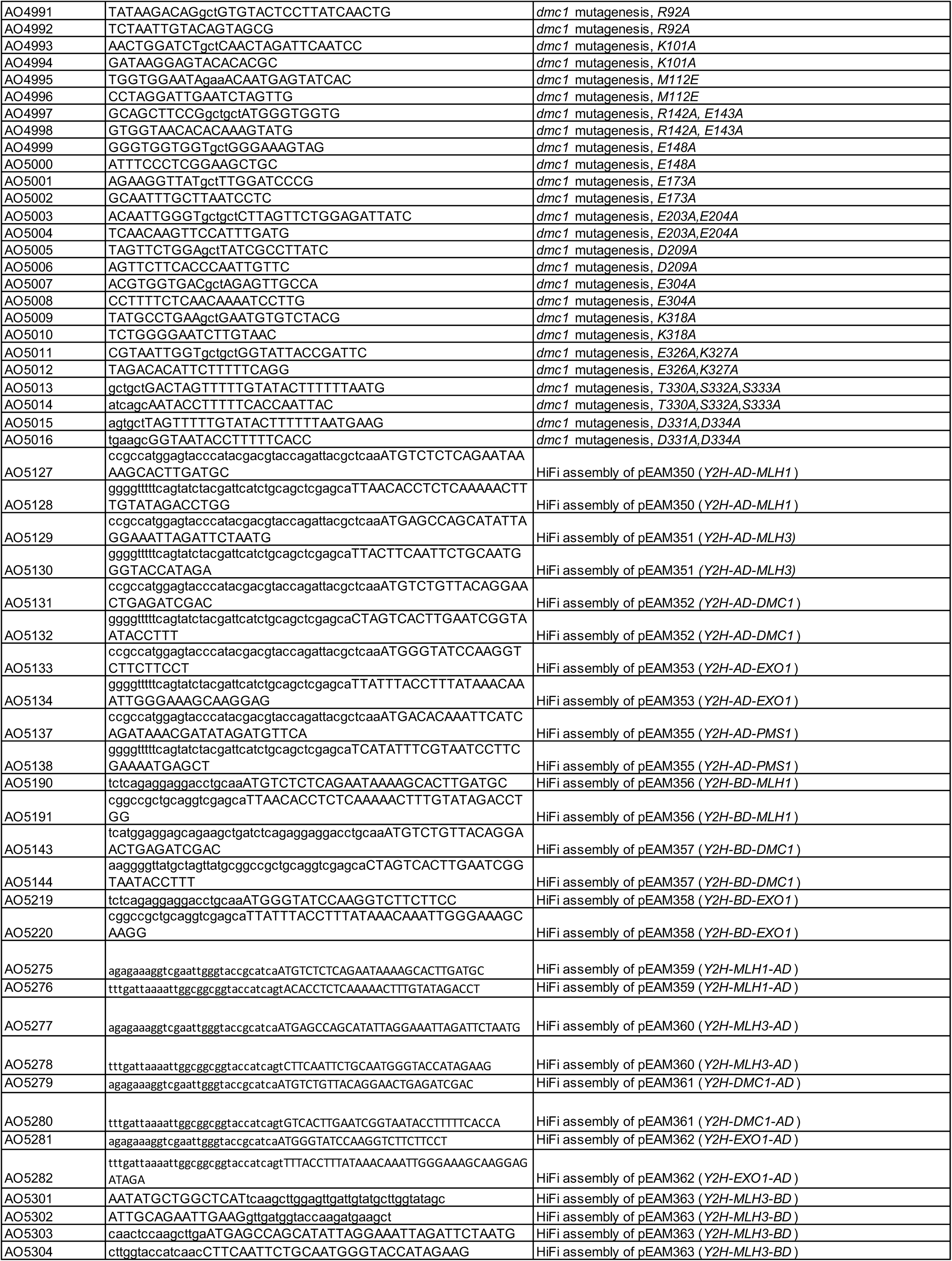
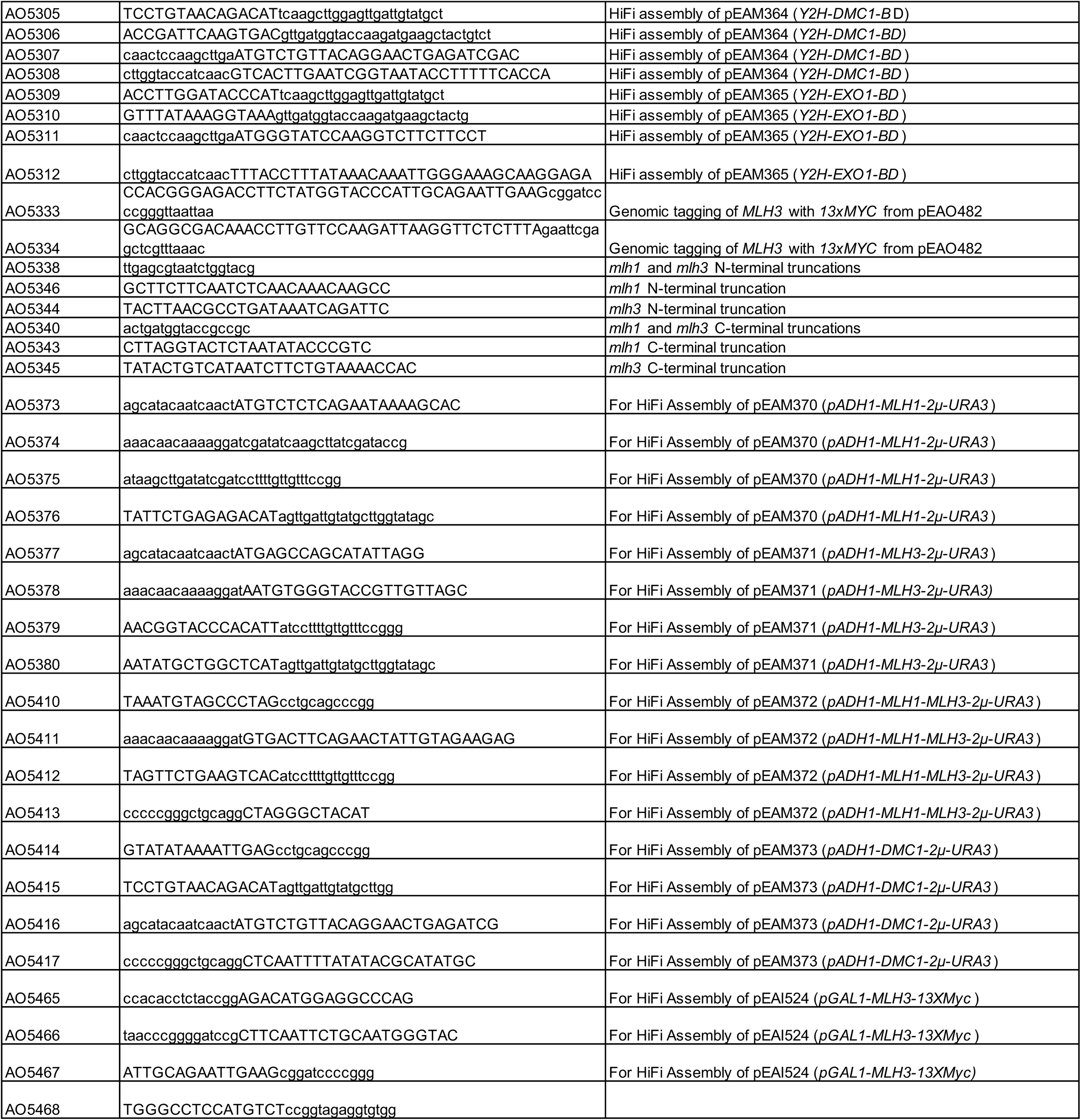
Oligonucleotides used in this study.

**Supplementary File 3.**
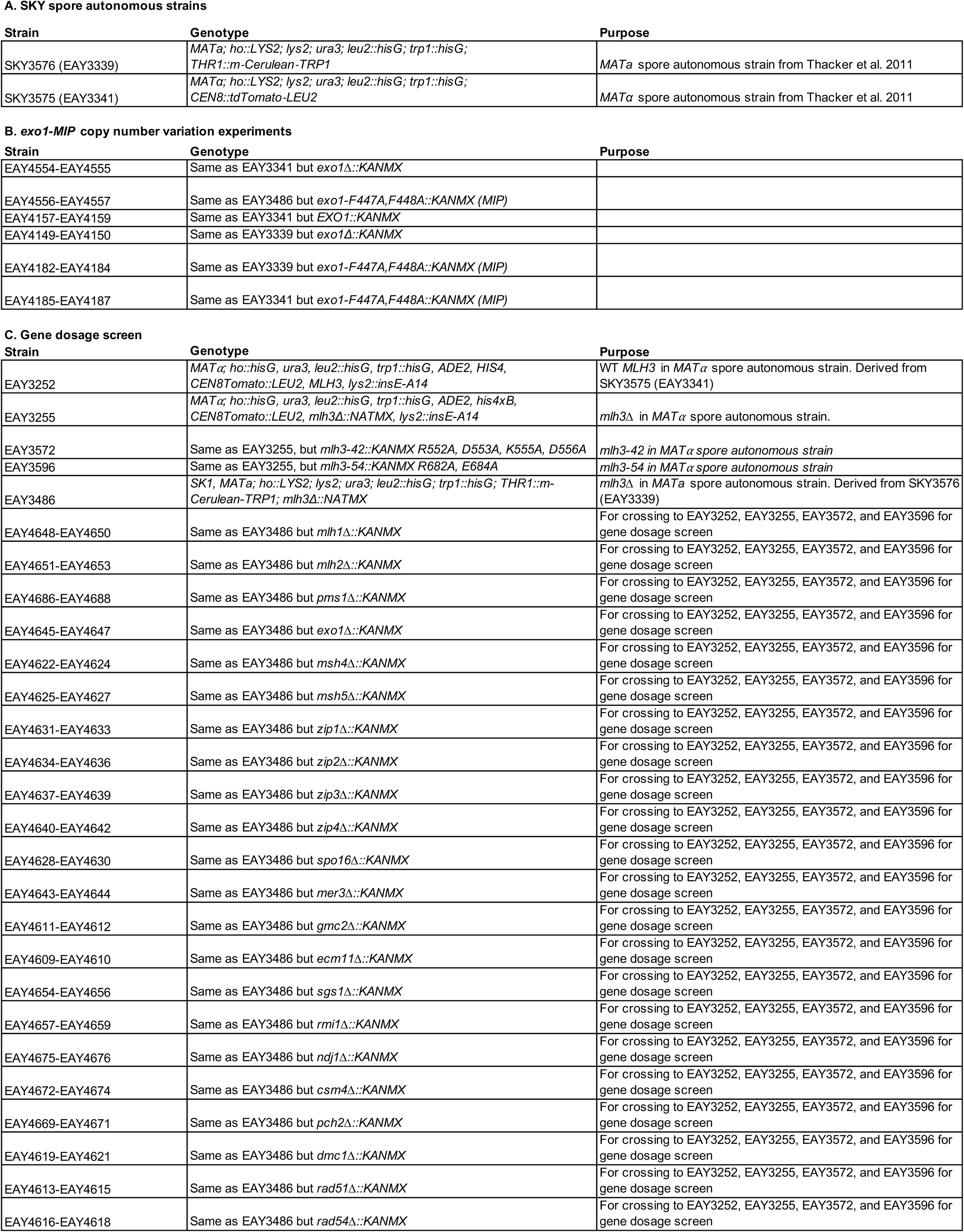
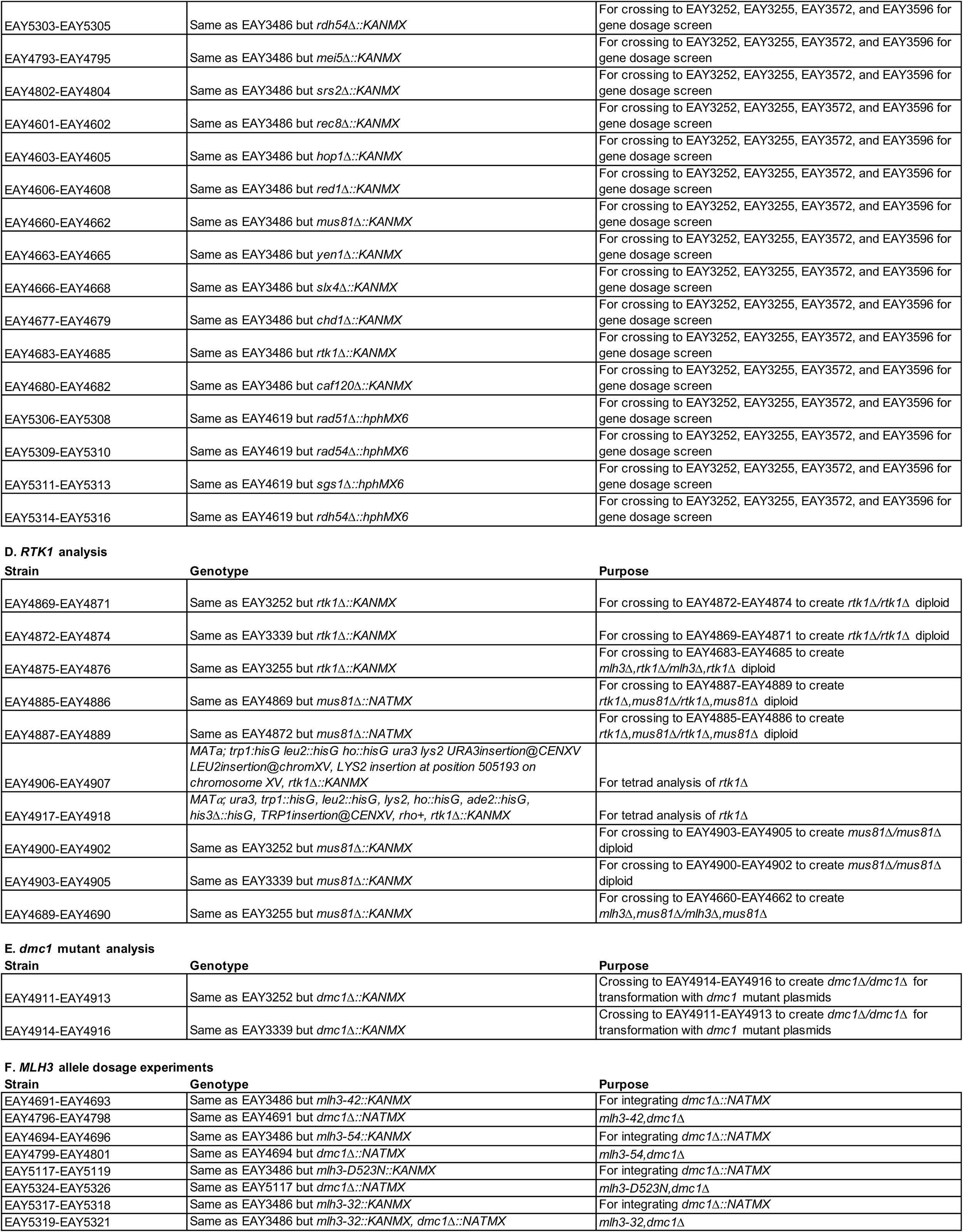
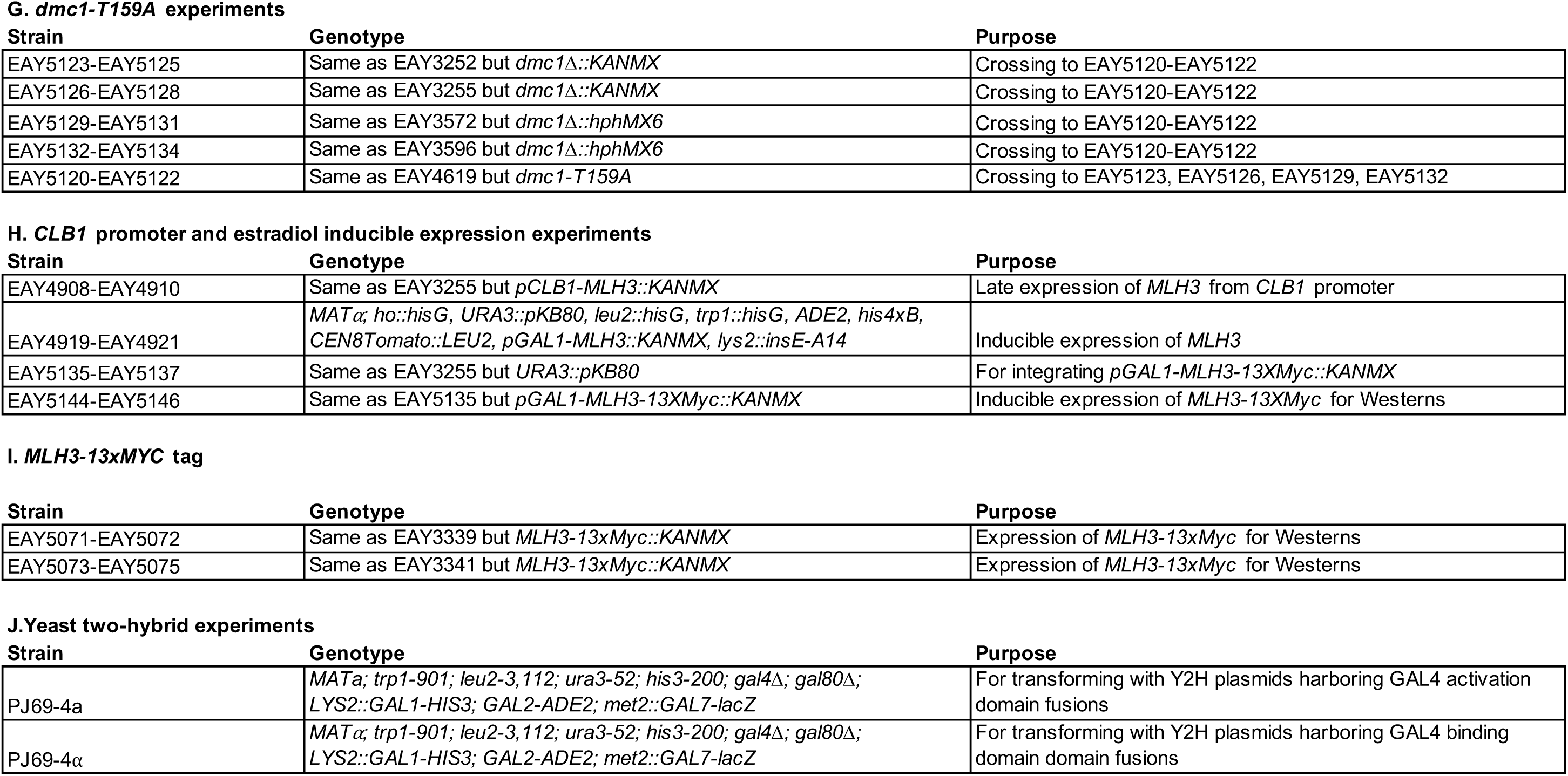
Strains used in this study.

**Supplementary File 4.**
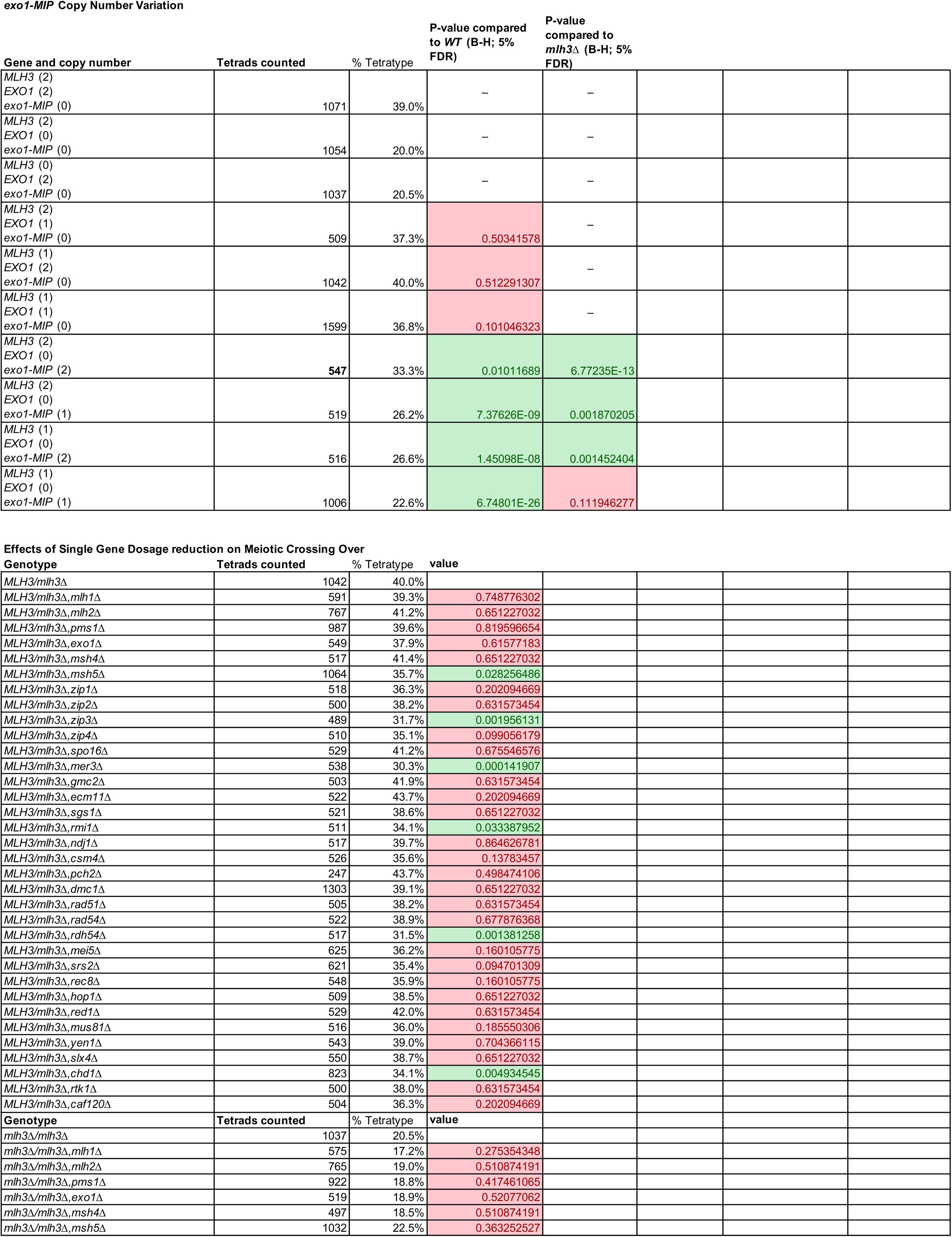
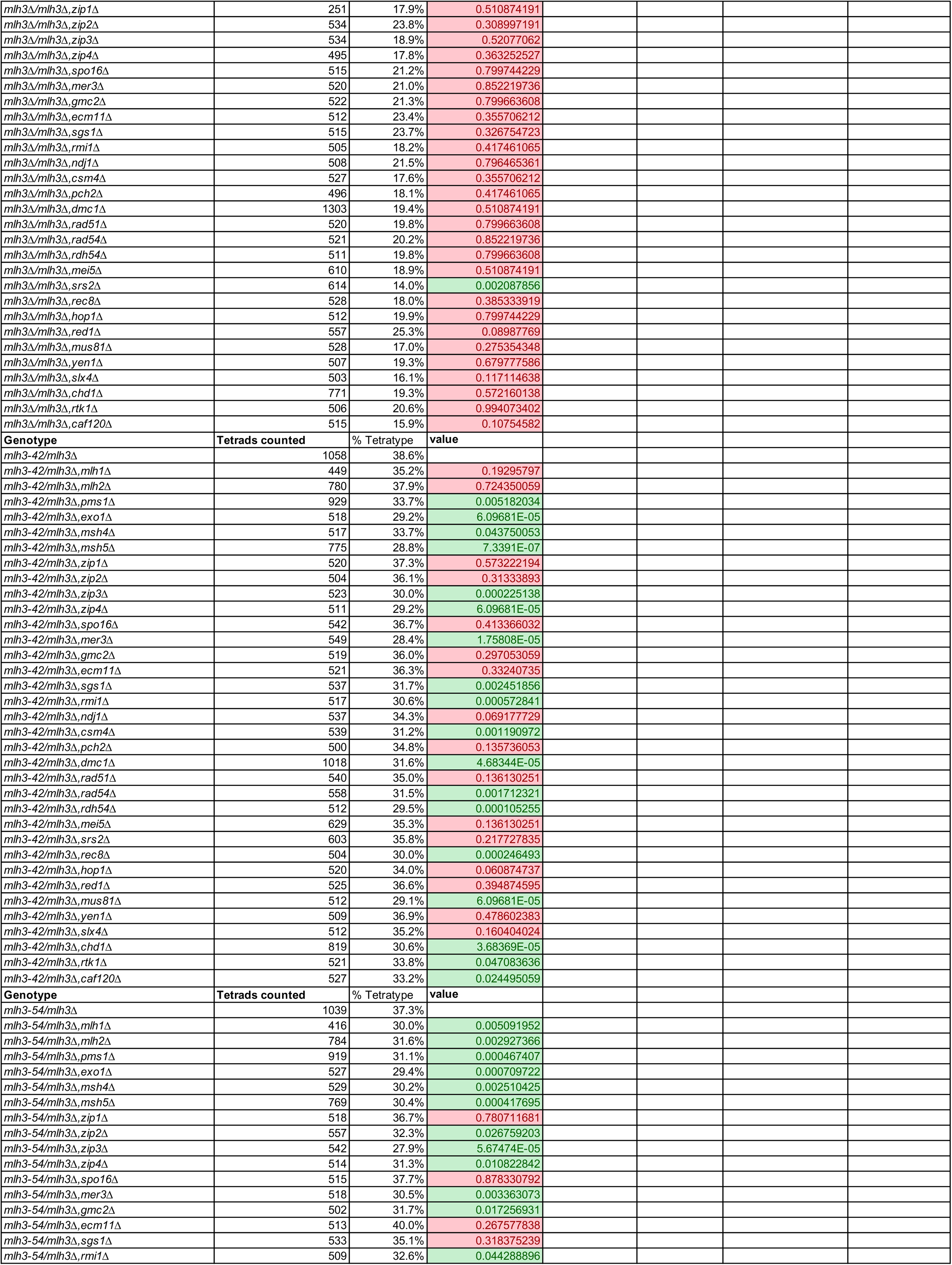
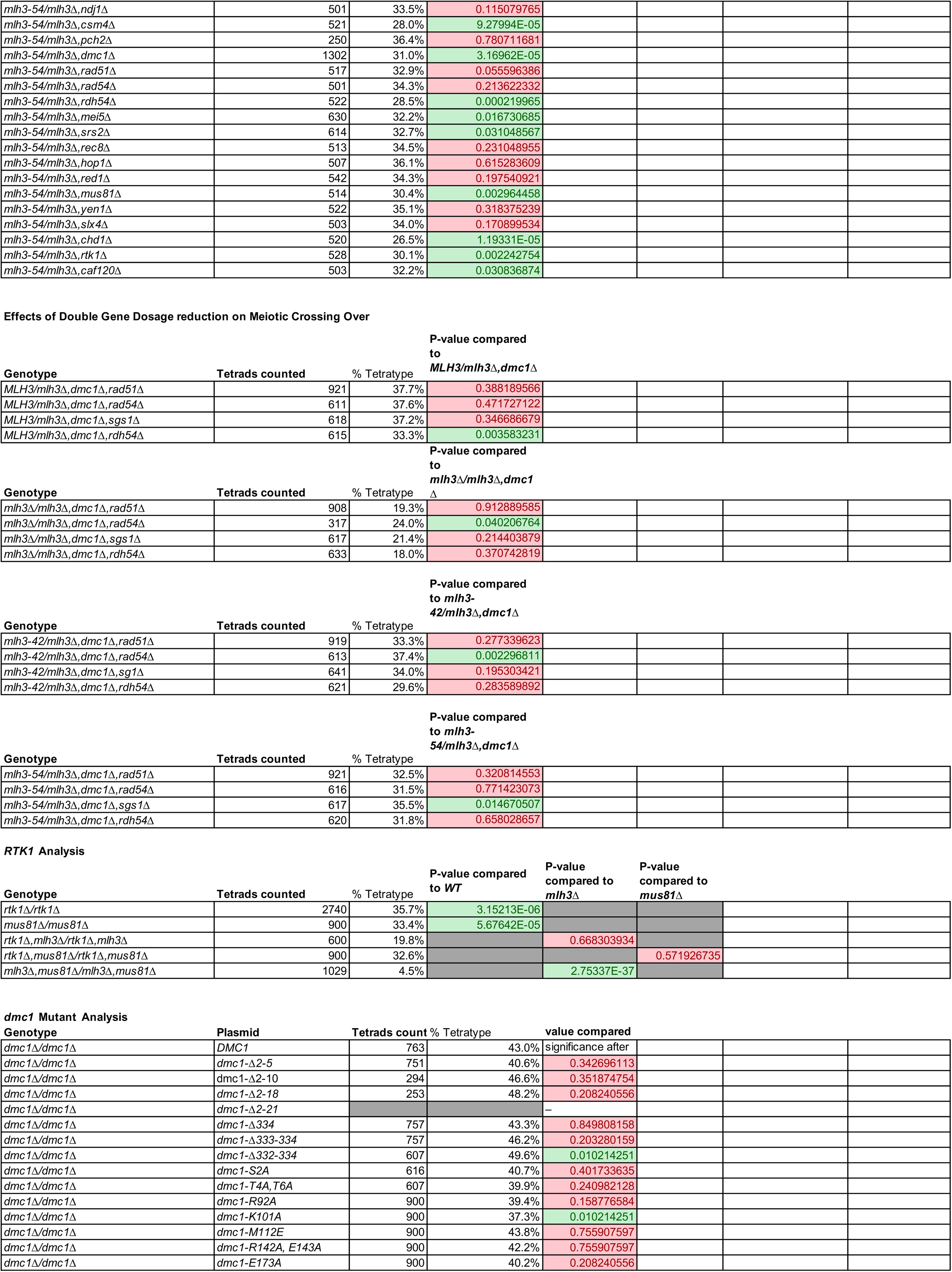
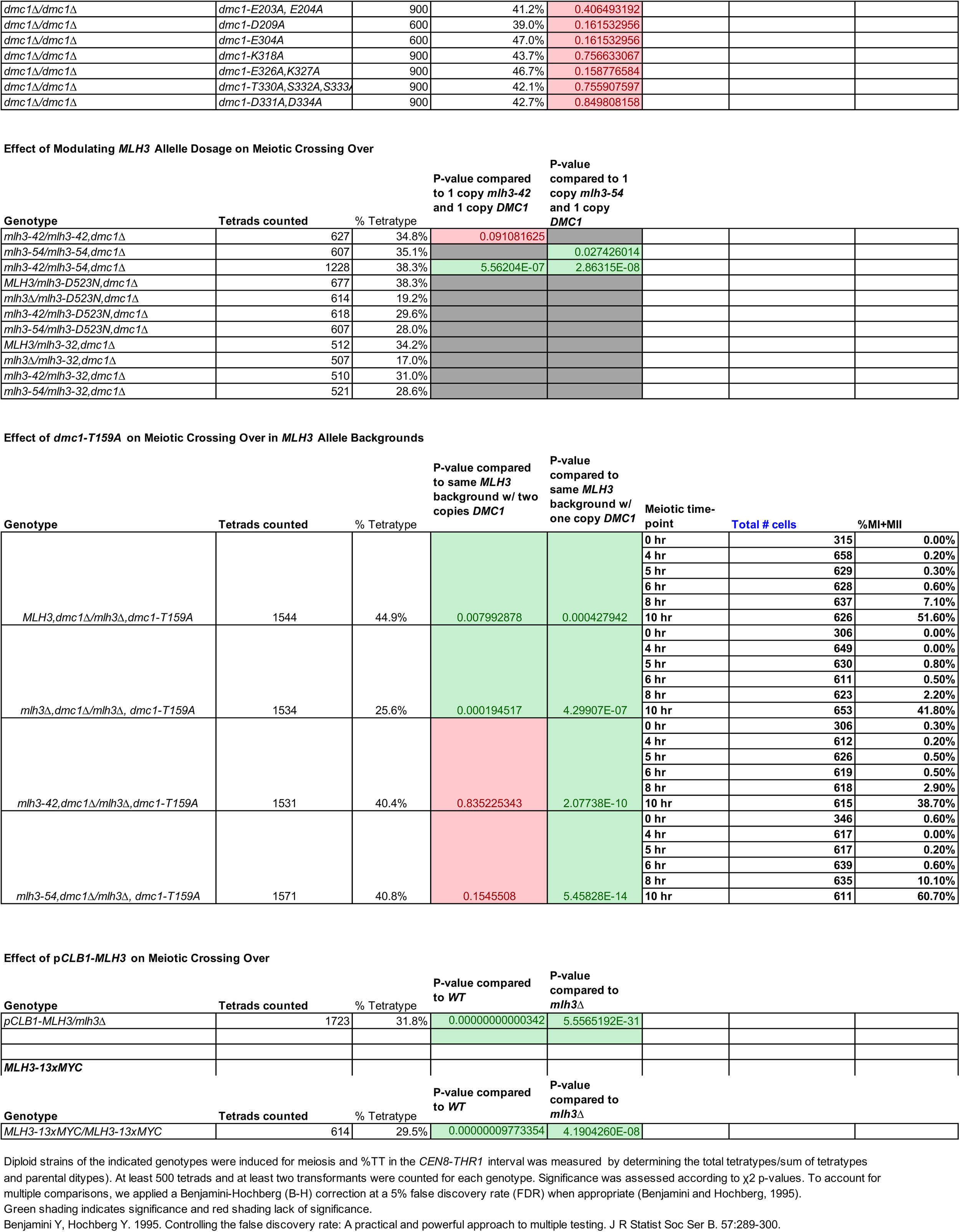
Spore Autonomous Data.

**Supplementary File 5.**
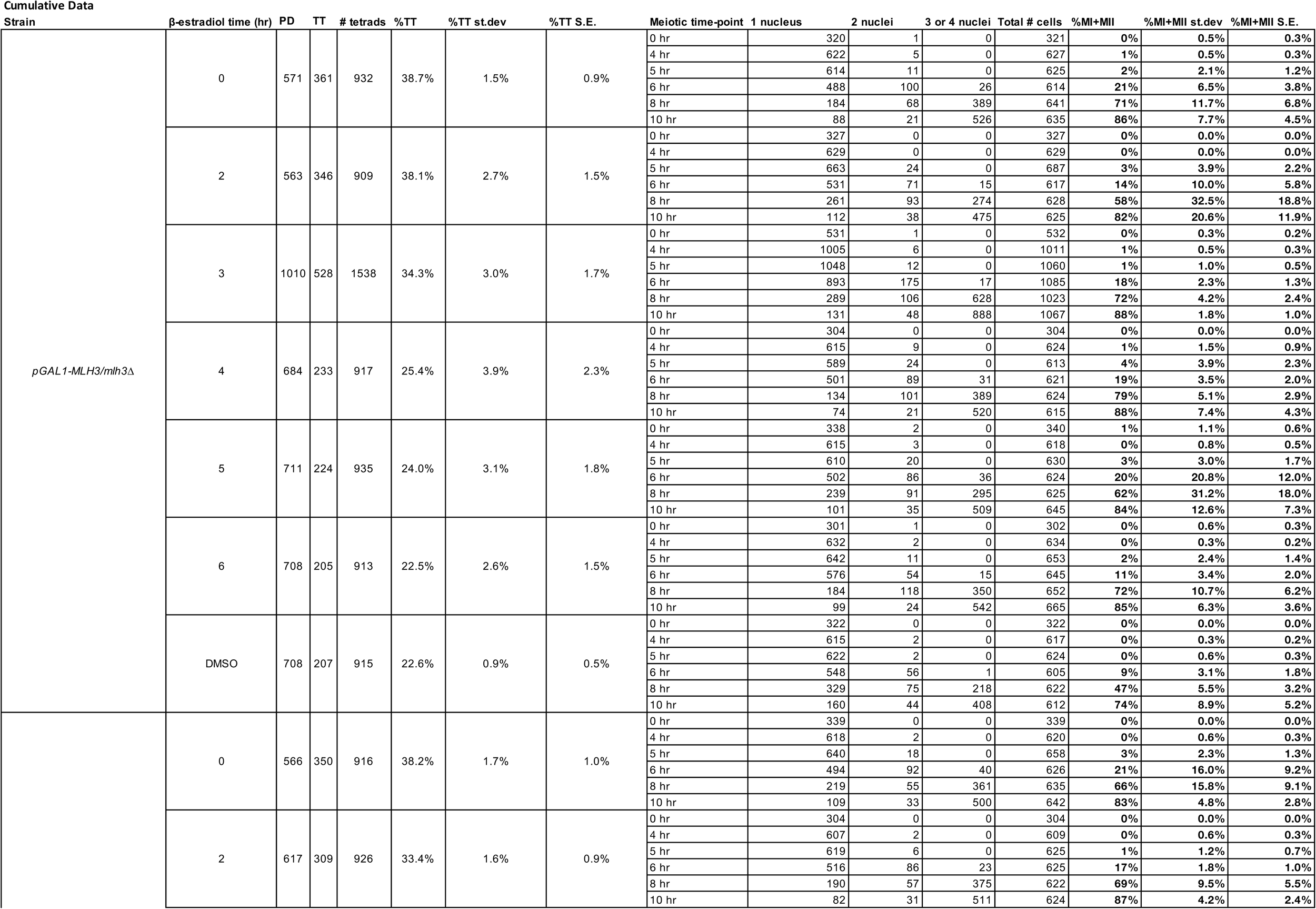
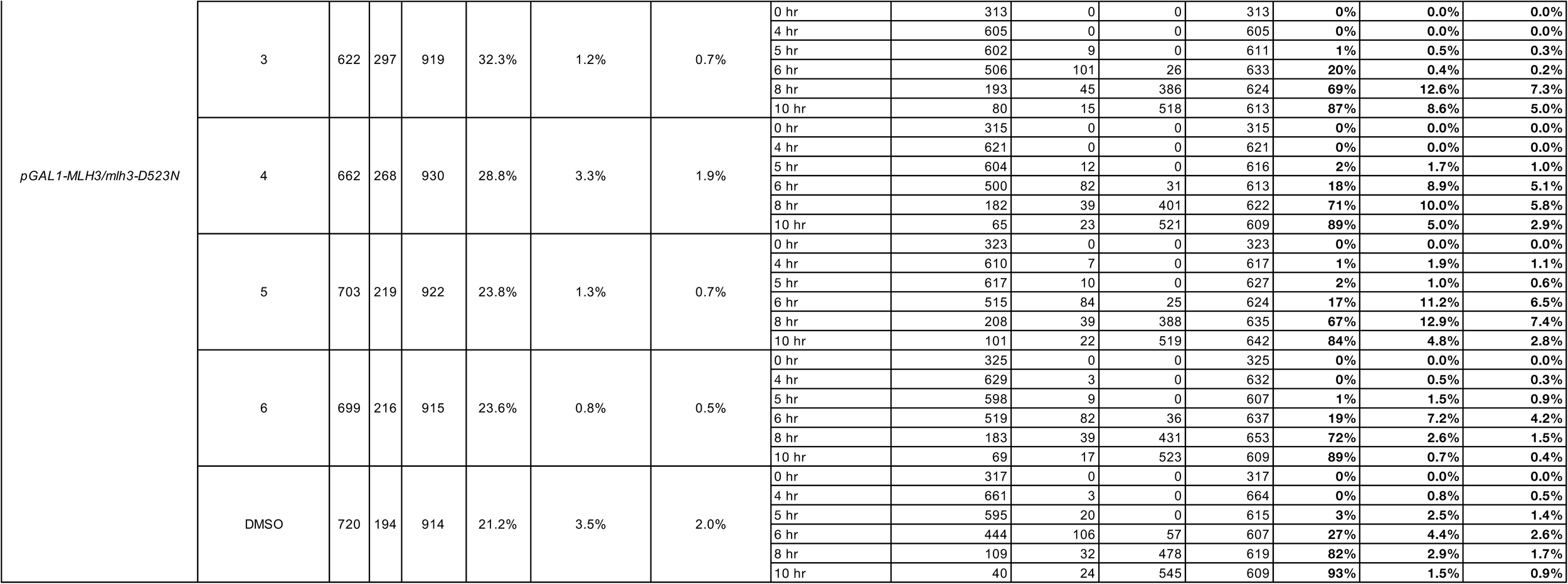
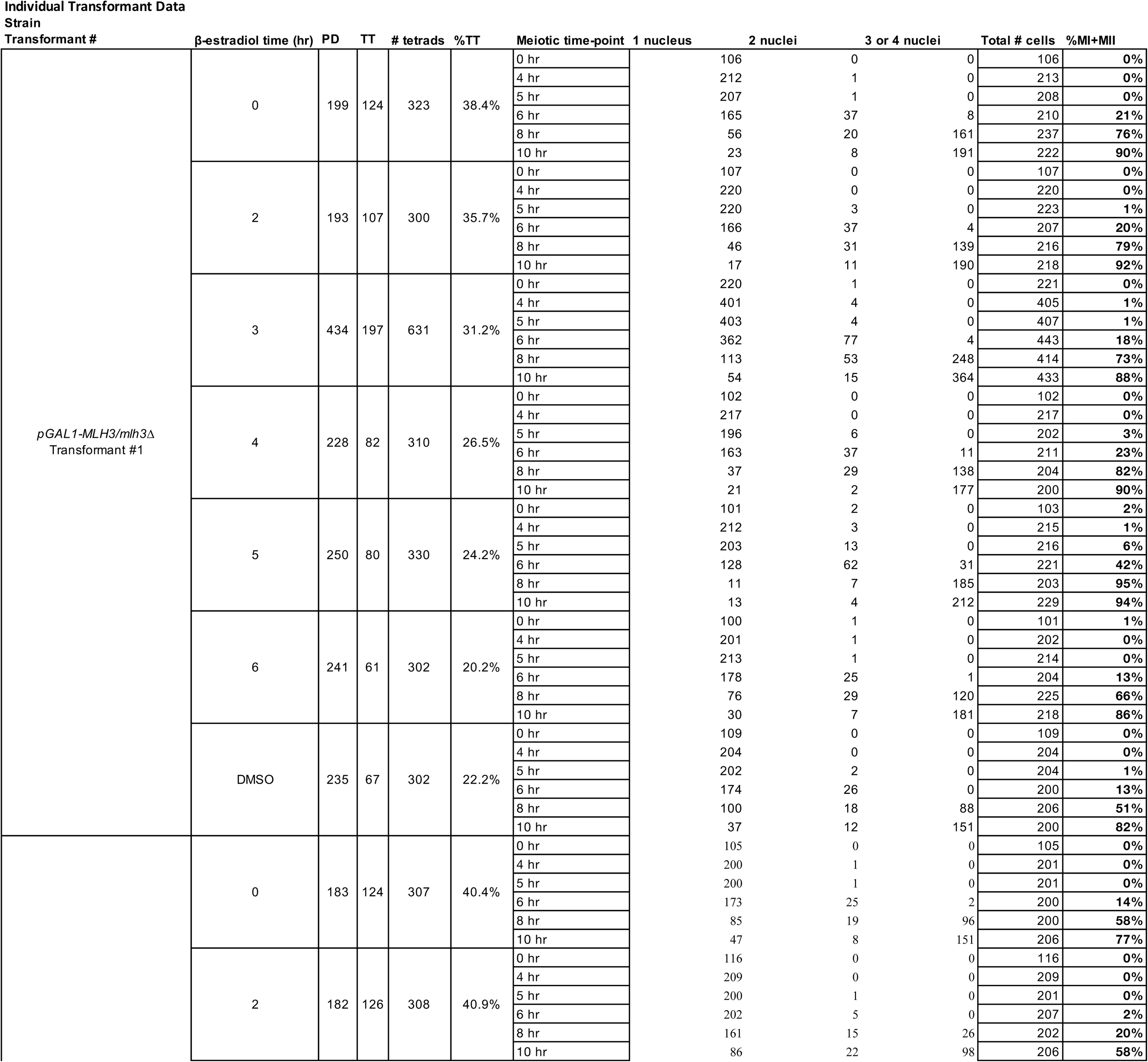
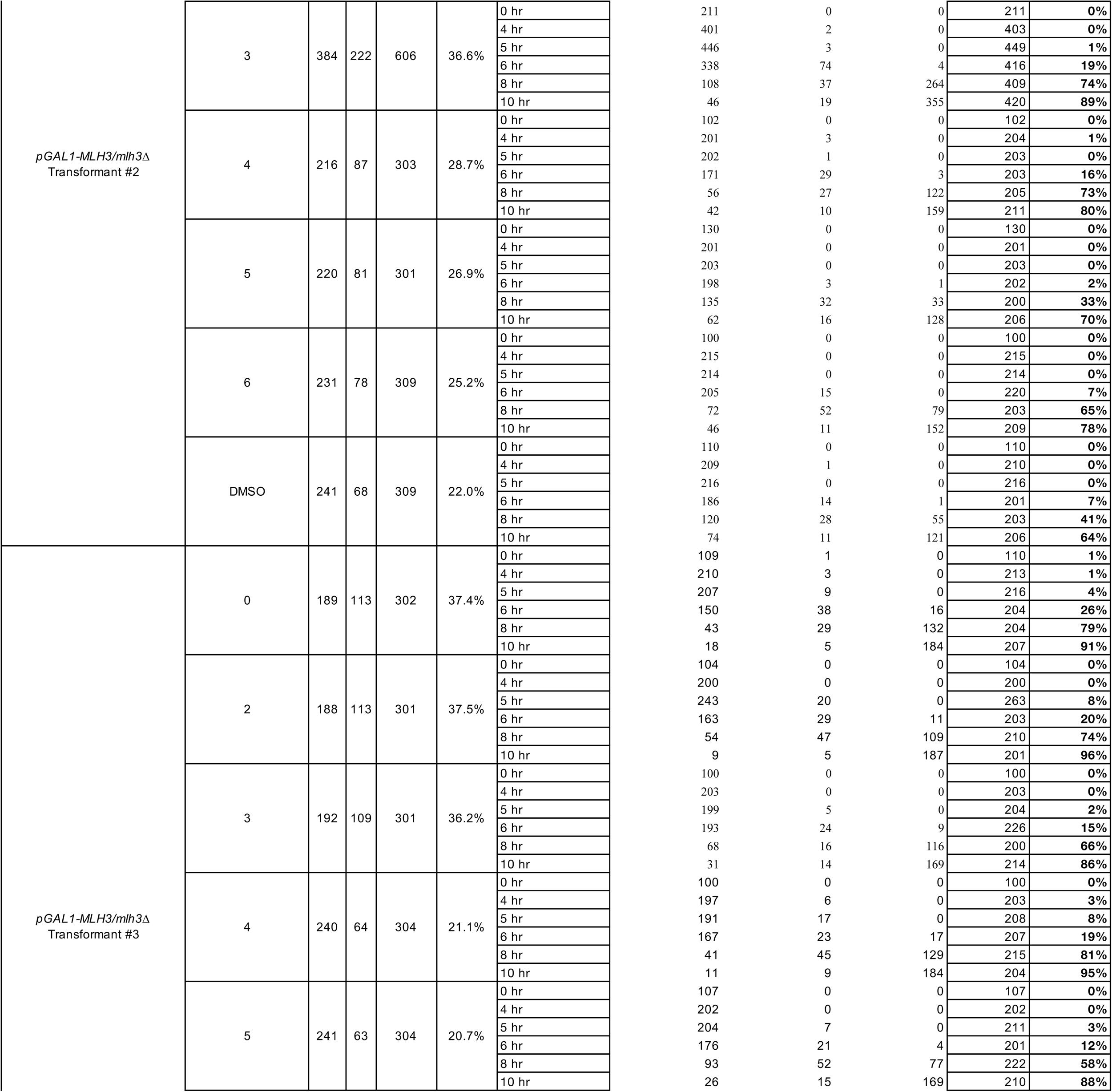
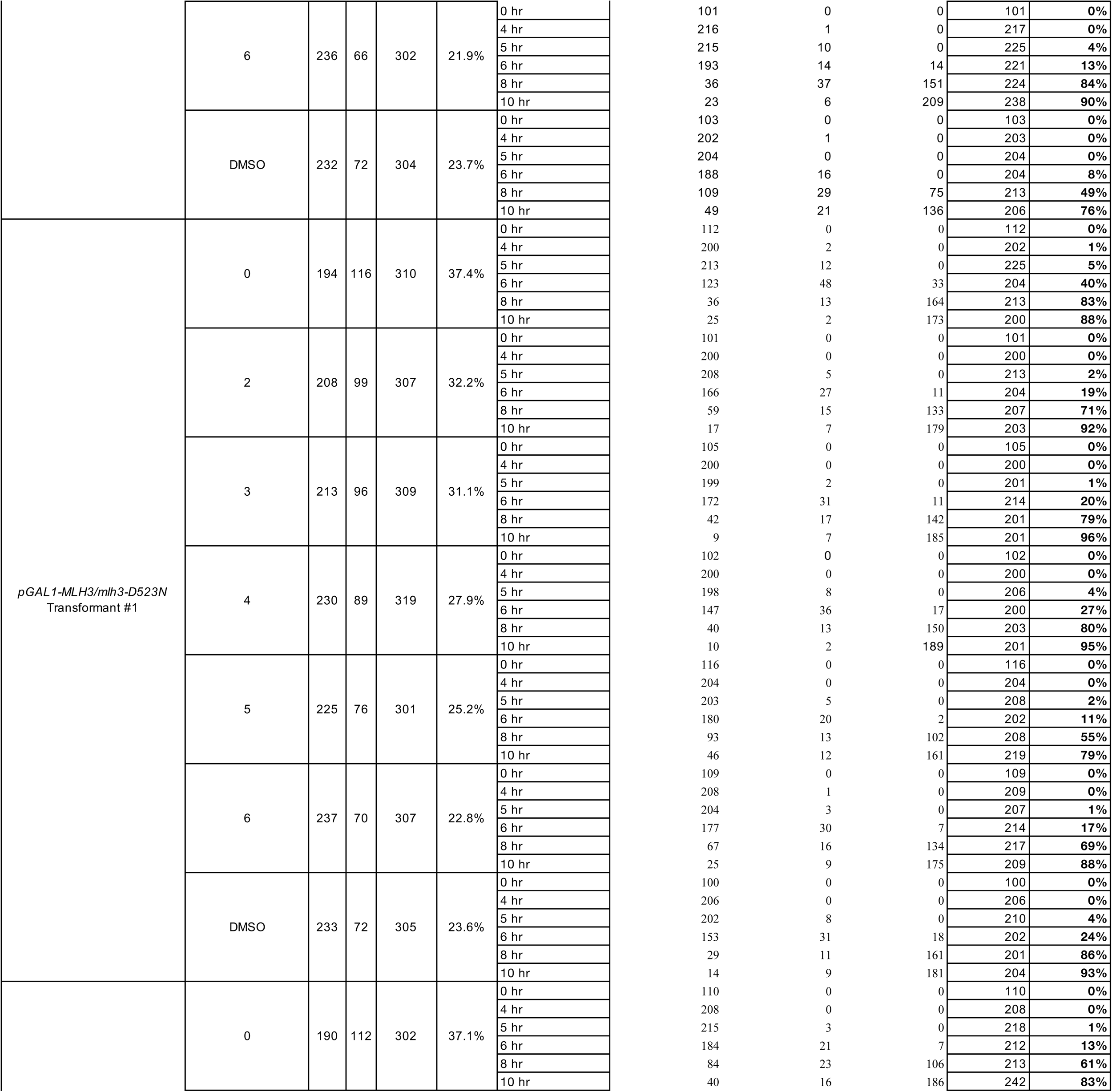
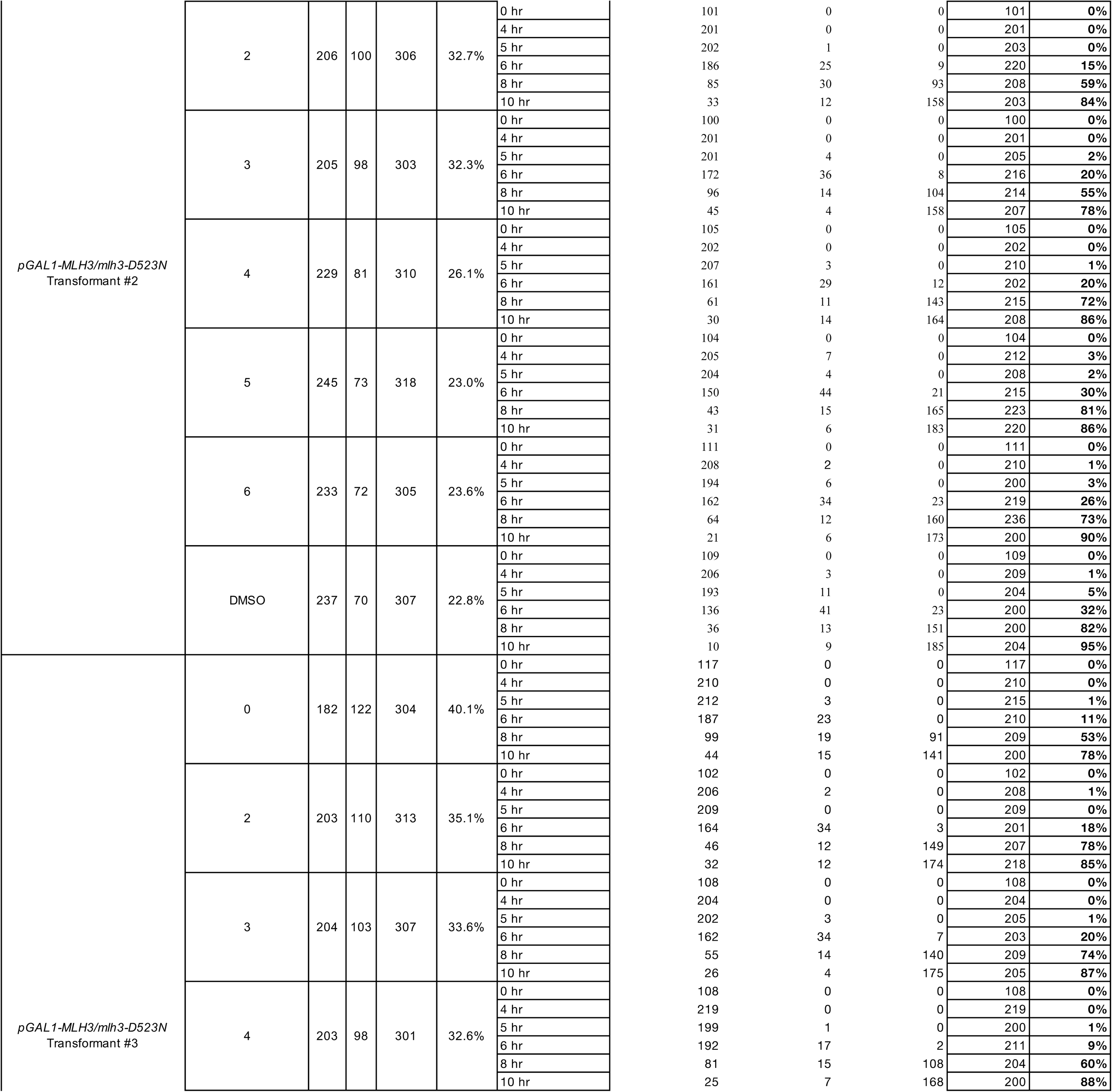
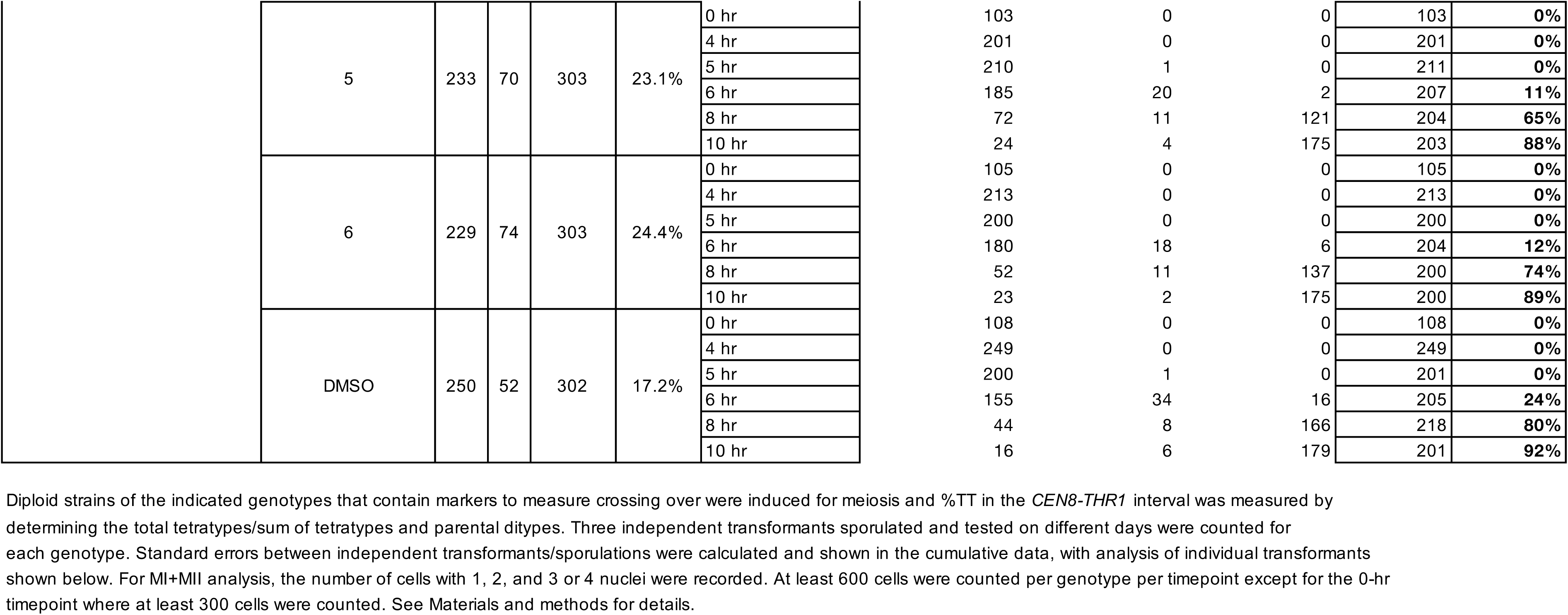
Time-course phenotypic data.

**Supplementary File 6.**
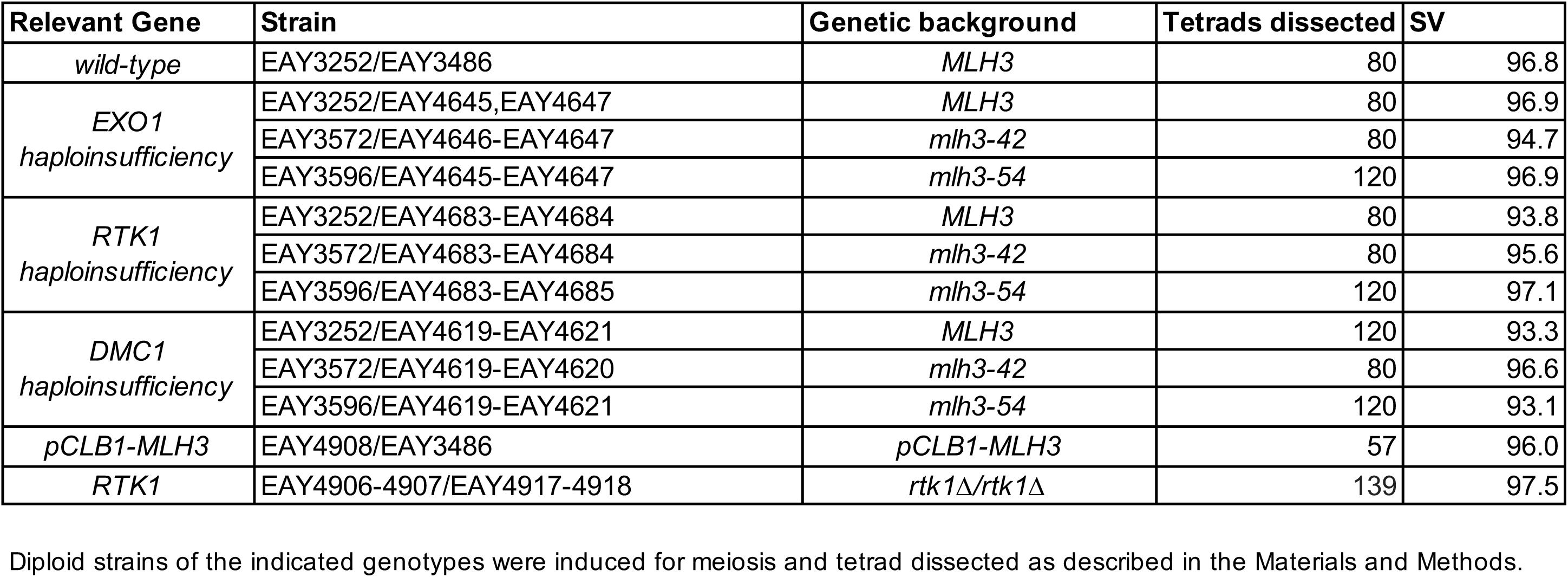
Spore viability data.

**Supplementary File 7.**
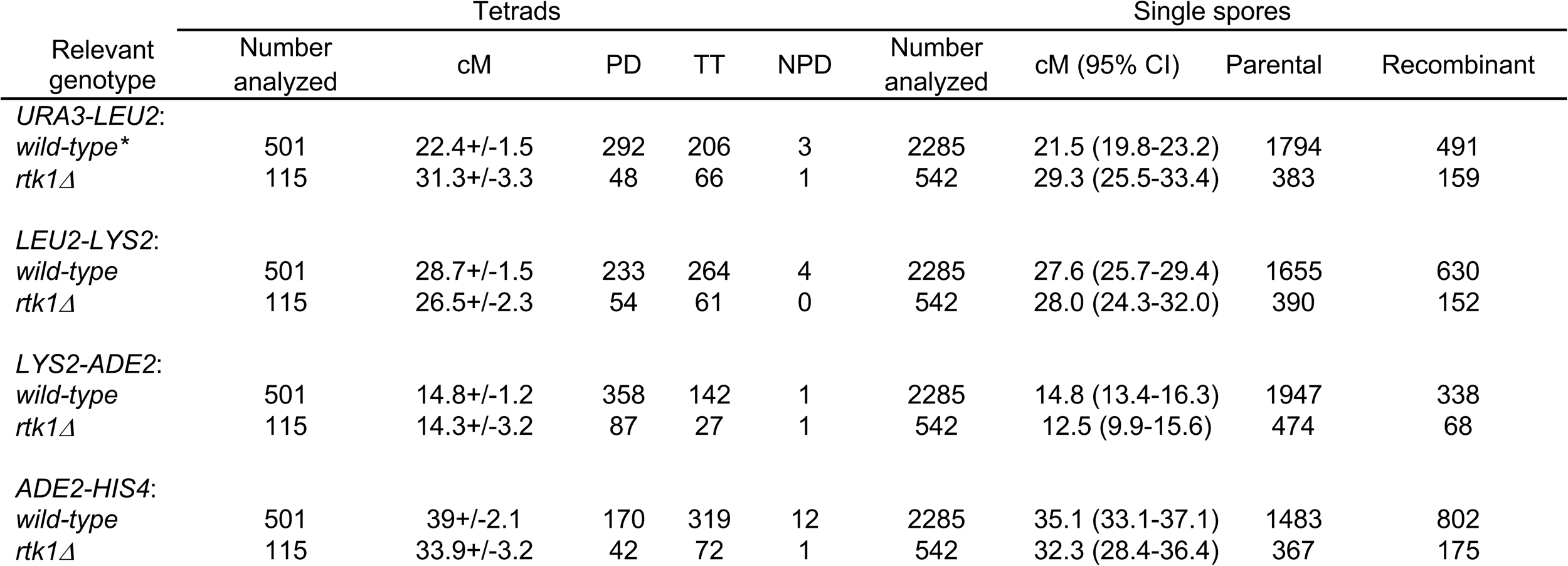
Genetic map distances (cM) and the distribution of parental and recombinant progeny for the EAY1108/EAY1112 strain background in *WT and rtk1Δ* background on Chromosome XV. Mutants are isogenic derivatives of EAY1108/EAY1112 (Supplementary File 3). Genetic intervals correspond to the genetic distance calculated from tetrads +/− one standard error. Standard error was calculated using the Stahl Laboratory Online Tools website (https://elizabethhousworth.com/StahlLabOnlineTools/). For single spore analysis, data are shown as 95% confidence intervals around the recombination frequency. For tetrad analysis the centimorgan (cM) map distance was calculated using the formula of Perkins (1949): 50{TT+(6NPD)}/(PD+TT+NPD). To compare to the tetrad data, recombination frequencies obtained from single spores (Parental/(Parental+Recombinant)) were multiplied by 100 to yield genetic map distances (cM). \**wild-type* was dissected as part of a larger data set (Gioia et al. 2023) in which *rtk11* was also dissected. Perkins DD. 1949. Biochemical mutants in the smut fungus *Ustilago maydis*. Genetics. 34:607-626. Gioia M, Payero L, Salim S, Fajish V G, Farnaz AF, Pannafino G, Chen JJ, Ajith VP, Momoh S, Scotland M, Raghavan V, Manhart CM, Shinohara A, Nishant KT, Alani E. 2023. Exo1 protects DNA nicks from ligation to promote crossover formation during meiosis. PLoS Biol. 21:e3002085.

| Relevant genotype | Tetrads |  |  |  |  | Single spores |  |  |  |
| --- | --- | --- | --- | --- | --- | --- | --- | --- | --- |
|  | Number analyzed | cM | PD | TT | NPD | Number analyzed | cM (95% CI) | Parental | Recombinant |
| *URA3-LEU2:* |  |  |  |  |  |  |  |  |  |
| *wild-type\** | 501 | 22.4+/-1.5 | 292 | 206 | 3 | 2285 | 21.5 (19.8-23.2) | 1794 | 491 |
| *rtk1Δ* | 115 | 31.3+/-3.3 | 48 | 66 | 1 | 542 | 29.3 (25.5-33.4) | 383 | 159 |
| *LEU2-LYS2:* |  |  |  |  |  |  |  |  |  |
| *wild-type* | 501 | 28.7+/-1.5 | 233 | 264 | 4 | 2285 | 27.6 (25.7-29.4) | 1655 | 630 |
| *rtk1Δ* | 115 | 26.5+/-2.3 | 54 | 61 | 0 | 542 | 28.0 (24.3-32.0) | 390 | 152 |
| *LYS2-ADE2:* |  |  |  |  |  |  |  |  |  |
| *wild-type* | 501 | 14.8+/-1.2 | 358 | 142 | 1 | 2285 | 14.8 (13.4-16.3) | 1947 | 338 |
| *rtk1Δ* | 115 | 14.3+/-3.2 | 87 | 27 | 1 | 542 | 12.5 (9.9-15.6) | 474 | 68 |
| *ADE2-HIS4:* |  |  |  |  |  |  |  |  |  |
| *wild-type* | 501 | 39+/-2.1 | 170 | 319 | 12 | 2285 | 35.1 (33.1-37.1) | 1483 | 802 |
| *rtk1Δ* | 115 | 33.9+/-3.2 | 42 | 72 | 1 | 542 | 32.3 (28.4-36.4) | 367 | 175 |

## Literature cited

1. Abdullah MF, Hoffmann ER, Cotton VE, Borts RH. 2004. A role for the MutL homologue MLH2 in controlling heteroduplex formation and in regulating between two different crossover pathways in budding yeast. Cytogenet Genome Res. 107:180–190.

2. Ahuja JS, Harvey CS, Wheeler DL, Lichten M. 2021. Repeated strand invasion and extensive branch migration are hallmarks of meiotic recombination. Mol Cell. 81:4258–4270.

3. Allers T, Lichten M. 2001. Differential timing and control of noncrossover and crossover recombination during meiosis. Cell. 106:47–57.

4. Altmannova V, Firlej M, Müller F, Janning P, Rauleder R, Rousova D, Schäffler A, Bange T, Weir JR. 2023. Biochemical characterisation of Mer3 helicase interactions and the protection of meiotic recombination intermediates. Nucleic Acids Res. 51:4363–4384. doi: 10.1093/nar/gkad175.

5. Al-Sweel N, Raghavan V, Datta A, Ajith VP, Di Vietro L, Khondakar N, et al. 2017. *mlh3* separation of function and endonuclease defective mutants display an unexpected effect on meiotic recombination outcomes. PLoS Genet. 13: e1006974.

6. Amin NS, Nguyen M, Oh S, Kolodner RD. 2001. *exo1*-dependent mutator mutation: Model system for studying functional interactions in mismatch repair. Mol Cell Biol. 21:5142–5155.

7. Argueso JL, Kijas AW, Sarin S, Heck J, Waase M, Alani, E. 2003. Systematic mutagenesis of the *Saccharomyces cerevisiae MLH1* gene reveals distinct roles for Mlh1p in meiotic crossing over and in vegetative and meiotic mismatch repair. Mol Cell Biol. 23:873–886.

8. Argueso JL, Wanat J, Gemici Z, Alani E. 2004. Competing crossover pathways act during meiosis in *Saccharomyces cerevisiae*. Genetics. 168:1805–1816.

9. Arter M, Hurtado-Nieves V, Oke A, Zhuge T, Wettstein R, Fung JC, Blanco MG, Matos J. 2018. Regulated Crossing-Over Requires Inactivation of Yen1/GEN1 Resolvase during Meiotic Prophase I. Dev Cell. 45:785–800.e6. doi: 10.1016/j.devcel.2018.05.020.

10. Baetz, KK, Korgan, NJ, Emili A, Greenblatt J, Hieter P. 2004. The ctf13-30/CTF13 Genomic Haploinsufficiency Modifier Screen Identifies the Yeast Chromatin Remodeling Complex RSC, Which Is Required for the Establishment of Sister Chromatid Cohesion. Mol Cell Biol. 24:1232–1244.

11. Bahler J, Wu JQ, Longtine MS, Shah NG, McKenzie A 3rd, Steever AB, et al. 1998. Heterologous modules for efficient and versatile PCR-based gene targeting in *Schizosaccharomyces pombe*. Yeast. 14:943–951.

12. Benjamin KR, Zhang C, Shokat KM, Herskowitz I. 2003. Control of landmark events in meiosis by the CDK Cdc28 and the meiosis-specific kinase Ime2. Genes Dev. 17:1524–1539. doi: 10.1101/gad.1101503.

13. Benjamini Y, Hochberg Y. 1995. Controlling the false discovery rate: A practical and powerful approach to multiple testing. J R Statist Soc Ser B. 57:289–300.

14. Bentolila S, Gipson AB, Kehl AJ, Hamm LN, Hayes ML, Mulligan RM, Hanson MR. 2021. A RanBP2-type zinc finger protein functions in intron splicing in Arabidopsis mitochondria and is involved in the biogenesis of respiratory complex I. Nucleic Acids Res. 49:3490–3506.

15. Bishop DK. 1994. RecA homologs Dmc1 and Rad51 interact to form multiple nuclear complexes prior to meiotic chromosome synapsis. Cell. 79:1081–1092.

16. Börner GV, Kleckner N, Hunter N. 2004. Crossover/noncrossover differentiation, synaptonemal complex formation, and regulatory surveillance at the leptotene/zygotene transition of meiosis. Cell. 117:29–45.

17. Bradford MM. 1976. A rapid and sensitive method for the quantitation of microgram quantities of protein utilizing the principle of protein-dye binding. Anal Biochem. 72:248–254.

18. Brar GA, Yassour M, Friedman N, Regev A, Ingolia NT, Weissman JS. 2012. High-resolution view of the yeast meiotic program revealed by ribosome profiling. Science. 335:552–557.

19. Brown MS, Bishop DK. 2014. DNA strand exchange and RecA homologs in meiosis. Cold Spring Harb Perspect Biol. 7:a016659. doi: 10.1101/cshperspect.a016659.

20. Brown MS, Grubb J, Zhang A, Rust MJ, Bishop DK. 2015. Small Rad51 and Dmc1 Complexes Often Co-occupy Both Ends of a Meiotic DNA Double Strand Break. PLoS Genet. 11:e1005653. doi: 10.1371/journal.pgen.1005653.

21. Bungard D, Reed M, Winter E. 2004. RSC1 and RSC2 Are Required for Expression of Mid-Late Sporulation-Specific Genes in *Saccharomyces cerevisiae*. Eukaryotic Cell 3:910–918.

22. Busygina V, Gaines WA, Xu Y, Kwon Y, Williams GJ, Lin SW, Chang HY, Chi P, Wang HW, Sung P. 2013. Functional attributes of the *Saccharomyces cerevisiae* meiotic recombinase Dmc1. DNA Repair. 12:707–12. doi: 10.1016/j.dnarep.2013.05.004.

23. Buzovetsky O, Kwon Y, Pham NT, Kim C, Ira G, Sung P, Xiong Y. 2017. Role of the Pif1-PCNA Complex in Pol δ-Dependent Strand Displacement DNA Synthesis and Break-Induced Replication. Cell Rep. 21:1707–1714. doi: 10.1016/j.celrep.2017.10.079.

24. Cannavo E, Sanchez A, Anand R, Ranjha L, Hugener J, Adam C, et al. 2020. Regulation of the MLH1-MLH3 endonuclease in meiosis. Nature 586:618–622.

25. Cao L, Alani E, Kleckner N. 1990. A pathway for generation and processing of double-strand breaks during meiotic recombination in *S. cerevisiae*. Cell. 61:1089–1101.

26. Chakraborty P, Pankajam AV, Lin G, Dutta A, Krishnaprasad GN, Tekkedil MM, et al. 2017. Modulating crossover frequency and interference for obligate crossovers in *Saccharomyces cerevisiae*. G3 7:1511–1524.

27. Chan YL, Zhang A, Weissman BP, Bishop DK. 2019. RPA resolves conflicting activities of accessory proteins during reconstitution of Dmc1-mediated meiotic recombination. Nucleic Acids Res. 47:747–761. doi: 10.1093/nar/gky1160.

28. Christianson TW, Sikorski RS, Dante M, Shero JH, Hieter P. 1992. Multifunctional yeast high-copy-number shuttle vectors. Gene 110:119–122.

29. Chu S, Herskowitz I. 1998. Gametogenesis in yeast is regulated by a transcriptional cascade dependent on Ndt80. Mol Cell. 1:685–96. doi: 10.1016/s1097-2765(00)80068-4.

30. Cloud V, Chan YL, Grubb J, Budke B, Bishop DK. 2012. Rad51 is an accessory factor for Dmc1-mediated joint molecule formation during meiosis. Science. 337:1222–1225. doi: 10.1126/science.1219379.

31. Crickard JB, Kaniecki K, Kwon Y, Sung P, Greene EC. 2018. Meiosis-specific recombinase Dmc1 is a potent inhibitor of the Srs2 antirecombinase. Proc Natl Acad Sci U S A. 115:E10041–E10048. doi: 10.1073/pnas.1810457115.

32. Dai J, Sanchez A, Adam C, Ranjha L, Reginato G, Chervy P, Tellier-Lebegue C, Andreani J, Guérois R, Ropars V, Le Du MH, Maloisel L, Martini E, Legrand P, Thureau A, Cejka P, Borde V, Charbonnier JB. 2021. Molecular basis of the dual role of the Mlh1-Mlh3 endonuclease in MMR and in meiotic crossover formation. Proc Natl Acad Sci U S A. 118: e2022704118.

33. De Muyt A, Pyatnitskaya A, Andréani J, Ranjha L, Ramus C, Laureau R, et al. 2018. A meiotic XPF-ERCC1-like complex recognizes joint molecule recombination intermediates to promote crossover formation. Genes Dev. 32:283–296.

34. Duroc Y, Kumar R, Ranjha L, Adam C, Guérois R, Md Muntaz K, Marsolier-Kergoat MC, Dingli F, Laureau R, Loew D, Llorente B, Charbonnier JB, Cejka P, Borde V. 2017. Concerted action of the MutLβ heterodimer and Mer3 helicase regulates the global extent of meiotic gene conversion. Elife. 6:e21900. doi: 10.7554/eLife.21900.

35. Fiorentini P, Huang KN, Tishkoff DX, Kolodner RD, Symington LS. 1997. Exonuclease I of *Saccharomyces cerevisiae* functions in mitotic recombination *in vivo* and *in vitro*. Mol Cell Biol. 17: 2764–2773.

36. Fung JC, Rockmill B, Odell M, Roeder GS. 2004. Imposition of crossover interference through the nonrandom distribution of synapsis initiation complexes. Cell. 116:795–802.

37. Gellon L, Werner M, Boiteux S. 2002. Ntg2p, a *Saccharomyces cerevisiae* DNA *N*-glycosylase/apurinic or apyrimidinic lyase involved in base excision repair of oxidative DNA damage, interacts with the DNA mismatch repair protein Mlh1p. Identification of a Mlh1p binding motif. J Biol Chem. 277:29963–29972.

38. Giaever G, Nislow C. 2014. The yeast deletion collection: a decade of functional genomics. Genetics. 197:451–465.

39. Gietz RD, Schiestl RH, Willems AR, Woods RA. 1995. Studies on the transformation of intact yeast cells by the LiAc/SS-DNA/PEG procedure. Yeast. 11: 355–360.

40. Gioia M, Payero L, Salim S, Fajish V G, Farnaz AF, Pannafino G, Chen JJ, Ajith VP, Momoh S, Scotland M, Raghavan V, Manhart CM, Shinohara A, Nishant KT, Alani E. 2023. Exo1 protects DNA nicks from ligation to promote crossover formation during meiosis. PLoS Biol. 21:e3002085.

41. Goldstein AL, McCusker JH. 1999. Three new dominant drug resistance cassettes for gene disruption in *Saccharomyces cerevisiae*. Yeast. 15:1541–1553.

42. Hall MC, Wang H, Erie DA, Kunkel TA. 2001. High affinity cooperative DNA binding by the yeast Mlh1-Pms1 heterodimer. J Mol Biol. 312:637–647.

43. Hassold T, Hunt P. 2001. To err (meiotically) is human: the genesis of human aneuploidy. Nat Rev Genet. 2:280–291.

44. Hillers KJ. 2004. Crossover interference. Curr Biol. 14:R1036–R1037.

45. Hinch AG, Becker PW, Li T, Moralli D, Zhang G, Bycroft C, Green C, Keeney S, Shi Q, Davies B, Donnelly P. 2020. The Configuration of RPA, RAD51, and DMC1 Binding in Meiosis Reveals the Nature of Critical Recombination Intermediates. Mol Cell. 79: 689–701.e10. doi: 10.1016/j.molcel.2020.06.015.

46. Hinch AG, Zhang G, Becker PW, Moralli D, Hinch R, Davies B, Bowden R, Donnelly P. 2019. Factors influencing meiotic recombination revealed by whole-genome sequencing of single sperm. Science. 363:eaau8861. doi: 10.1126/science.aau8861.

47. Ho B, Baryshnikova A, Brown GW. 2018. Unification of Protein Abundance Datasets Yields a Quantitative *Saccharomyces cerevisiae* Proteome. Cell Syst. 6:192–205.e3. doi: 10.1016/j.cels.2017.12.004.

48. Hoffman CS, Winston F. 1987. A ten-minute DNA preparation from yeast efficiently releases autonomous plasmids for transformation of *Escherichia coli*. Gene. 57:267–272.

49. Hunt LJ, Ahmed EA, Kaur H, Ahuja JS, Hulme L, Chou TC, Lichten M, Goldman ASH. 2019. *S. cerevisiae* Srs2 helicase ensures normal recombination intermediate metabolism during meiosis and prevents accumulation of Rad51 aggregates. Chromosoma. 128:249–265. doi: 10.1007/s00412-019-00705-9.

50. Hunter N, Kleckner, N. 2001. The single-end invasion: An asymmetric intermediate at the double-strand break to double-Holliday Junction transition of meiotic recombination. Cell. 106:59–70.

51. Hunter N. 2015. Meiotic recombination: The essence of heredity. Cold Spring Harb Perspect Biol. 7:a016618.

52. Jones GH, Franklin FCH. 2006. Meiotic crossing-over: Obligation and interference. Cell. 126:246–248.

53. Kadyrov FA, Dzantiev L, Constantin N, Modrich P. 2006. Endonucleolytic function of MutLalpha in human mismatch repair. Cell. 126:297–308. doi: 10.1016/j.cell.2006.05.039.

54. Kane SM, Roth R. 1974. Carbohydrate metabolism during ascospore development in yeast. J Bacteriol. 118:8–14.

55. Keeney S, Giroux CN, Kleckner N. 1997. Meiosis-specific DNA double-strand breaks are catalyzed by Spo11, a member of a widely conserved protein family. Cell. 88:375–384.

56. Kolas NK, Svetlanov A, Lenzi ML, Macaluso FP, Lipkin SM, Liskay RM, Greally J, Edelmann W, Cohen PE. 2005.Localization of MMR proteins on meiotic chromosomes in mice indicates distinct functions during prophase I. J Cell Biol. 171:447–458.

57. Kosaka H, Shinohara M, Shinohara A. 2008. Csm4-dependent telomere movement on nuclear envelope promotes meiotic recombination. PLoS Genet. 4:e1000196. doi: 10.1371/journal.pgen.1000196.

58. Krejci L, Van Komen S, Li Y, Villemain J, Reddy MS, Klein H, Ellenberger T, Sung P. 2003. DNA helicase Srs2 disrupts the Rad51 presynaptic filament. Nature. 423:305–309. doi: 10.1038/nature01577. PMID: 12748644.

59. Kulkarni DS, Owens SN, Honda M, Ito M, Yang Y, Corrigan MW, et al. 2020. PCNA activates the MutL endonuclease to promote meiotic crossing over. Nature. 586:623–627.

60. Liu Y, Gaines WA, Callender T, Busygina V, Oke A, Sung P, Fung JC, Hollingsworth NM. 2014. Down-regulation of Rad51 activity during meiosis in yeast prevents competition with Dmc1 for repair of double-strand breaks. PLoS Genet. 10:e1004005. doi: 10.1371/journal.pgen.1004005.

61. Longtine MS, McKenzie A 3rd, Demarini DJ, Shah NG, Wach A, Brachat A, et al. 1998. Additional modules for versatile and economical PCR-based gene deletion and modification in *Saccharomyces cerevisiae*. Yeast. 14:953–961.

62. Lynn A, Soucek R, Börner GV. 2007. ZMM proteins during meiosis: Crossover artists at work. Chromosome Res. 15:591–605.

63. Maguire MP. 1974. The need for a chiasma binder. J Theor Biol. 48:485–487.

64. Mancera E, Bourgon R, Brozzi A, Huber W, Steinmetz LM. 2008. High-resolution mapping of meiotic crossovers and non-crossovers in yeast. Nature 454:479–485.

65. Manhart CM, Ni X, White MA, Ortega J, Surtees JA, Alani E. 2017. The mismatch repair and meiotic recombination endonuclease Mlh1-Mlh3 is activated by polymer formation and can cleave DNA substrates in trans. PLoS Biol. 15: e2001164.

66. Marsolier-Kergoat, MC, Khan MM, Schott J, Zhu X, Llorente B. 2018. Mechanistic view and genetic control of DNA recombination during meiosis. Mol Cell. 70:9–20.

67. Martini E, Borde V, Legendre M, Audic S, Regnault B, Soubigou G, et al. 2011. Genome-wide analysis of heteroduplex DNA in mismatch repair–deficient yeast cells reveals novel properties of meiotic recombination pathways. PLoS Genet. 7: e1002305.

68. Martini E, Diaz RL, Hunter N, Keeney S. 2006. Crossover homeostasis in yeast meiosis. Cell. 126:285–295.

69. Nagaoka SI, Hassold TJ, Hunt PA. 2010. Human aneuploidy: mechanisms and new insights into an age-old problem. Nat Rev Genet. 13:493–504.

70. Nicolette ML, Lee K, Guo Z, Rani M, Chow JM, Lee SE, Paull TT. 2010. Mre11-Rad50-Xrs2 and Sae2 promote 5’ strand resection of DNA double-strand breaks. Nat Struct Mol Biol. 17:1478–1485.

71. Nishant KT, Plys AJ, Alani E. 2008. A mutation in the putative MLH3 endonuclease domain confers a defect in both mismatch repair and meiosis in *Saccharomyces cerevisiae*. Genetics. 179:747–755.

72. Novak JE, Ross-Macdonald, PB, Roeder GS. 2001. The budding yeast Msh4 protein functions in chromosome synapsis and the regulation of crossover distribution. Genetics. 158:1013–1025.

73. Pan J, Sasaki M, Kniewel R, Murakami H, Blitzbau HG, Tischfield SE, et al. 2011. Hierarchical Combination of Factors Shapes the Genome-Wide Topography of Yeast Meiotic Recombination Initiation. Cell. 144:719–731.

74. Pannafino G, Alani E. 2021. Coordinated and Independent Roles for MLH Subunits in DNA Repair. Cells. 10:948. doi: 10.3390/cells10040948

75. Perkins DD. 1949. Biochemical mutants in the smut fungus *Ustilago maydis*. Genetics. 34:607–626.

76. Peterson SE, Keeney S, Jasin M. 2020. Mechanistic insight into crossing over during mouse meiosis. Mol Cell. 78:1252–1263.e3. doi: 10.1016/j.molcel.2020.04.009.

77. Plys AJ, Rogacheva MV, Greene EC, Alani E. 2012. The unstructured linker arms of Mlh1-Pms1 are important for interactions with DNA during mismatch repair. J Mol Biol. 422:192–203.

78. Premkumar T, Paniker L, Kang R, Biot M, Humphrey E, Destain H, Ferranti I, Okulate I, Nguyen H, Kilaru V, Frasca M, Chakraborty P, Cole F. 2023. Genetic dissection of crossover mutants defines discrete intermediates in mouse meiosis. Mol Cell. 83:2941–2958.e7. doi: 10.1016/j.molcel.2023.07.022.

79. Pyatnitskaya A, Borde V, De Muyt A. 2019. Crossing and zipping: Molecular duties of the ZMM proteins in meiosis. Chromosoma. 128:181–198.

80. Rahman MM, Mohiuddin M, Shamima Keka I, Yamada K, Tsuda M, Sasanuma H, Andreani J, Guerois R, Borde V, Charbonnier JB, Takeda S. 2020. Genetic evidence for the involvement of mismatch repair proteins, PMS2 and MLH3, in a late step of homologous recombination. J Biol Chem. 295:17460–17475. doi: 10.1074/jbc.RA120.013521.

81. Ranjha L, Anand R, Cejka P. 2014. The *Saccharomyces cerevisiae* Mlh1-Mlh3 heterodimer Is an endonuclease that preferentially binds to Holliday Junctions. J Biol Chem. 289:5674–5686.

82. Rogacheva MV, Manhart CM, Chen C, Guarne A, Surtees J, Alani E. 2014. Mlh1-Mlh3, A meiotic crossover and DNA mismatch repair factor, is a Msh2-Msh3-stimulated endonuclease. J Biol Chem. 289:5664–5673.

83. Rose MD, Winston F, Hieter P. 1990. Methods in yeast genetics: A laboratory course manual. Cold Spring Harbor Laboratory Press. Cold Spring Harbor, NY.

84. Sanchez A, Adam C, Rauh F, Duroc Y, Ranjha L, Lombard B, et al. 2020. Exo1 recruits Cdc5 polo kinase to MutL to ensure efficient meiotic crossover formation. Proc Natl Acad Sci USA. 117:30577–30588.

85. Schwacha A, Kleckner N. 1997. Interhomolog bias during meiotic recombination: meiotic functions promote a highly differentiated interhomolog-only pathway. Cell. 90:1123–1135.

86. Schwacha A, Kleckner N. 1995. Identification of double Holliday Junctions as intermediates in meiotic recombination. Cell. 83:783–791.

87. Schwacha A, Kleckner N. 1994. Identification of joint molecules that form frequently between homologs but rarely between sister chromatids during yeast meiosis. Cell. 76:51–63.

88. Serrentino ME, Chaplais E, Sommermeyer V, Borde V. 2013. Differential association of the conserved SUMO ligase Zip3 with meiotic double-strand break sites reveals regional variations in the outcome of meiotic recombination. PLoS Genet. 9:e1003416.

89. Shinohara M, Oh SD, Hunter N, Shinohara A. 2008. Crossover assurance and crossover interference are distinctly regulated by the ZMM proteins during yeast meiosis. Nat Genet. 40:299–309.

90. Shinohara M, Sakai K, Shinohara A, Bishop DK. 2003. Crossover interference in *Saccharomyces cerevisiae* requires a TID1/RDH54- and DMC1-dependent pathway. Genetics. 163:1273–1286.

91. Slotman JA, Paul MW, Carofiglio F, de Gruiter HM, Vergroesen T, Koornneef L, van Cappellen WA, Houtsmuller AB, Baarends WM. 2020. Super-resolution imaging of RAD51 and DMC1 in DNA repair foci reveals dynamic distribution patterns in meiotic prophase. PLoS Genet. 16:e1008595. doi: 10.1371/journal.pgen.1008595.

92. Snowden T, Acharya S, Butz C, Berardini M, Fishel R. 2004. hMSH4-hMSH5 recognizes holliday junctions and forms a meiosis-specific sliding clamp that embraces homologous chromosomes. Mol Cell. 15:437–451.

93. Steinfeld JB, Beláň O, Kwon Y, Terakawa T, Al-Zain A, Smith MJ, Crickard JB, Qi Z, Zhao W, Rothstein R, Symington LS, Sung P, Boulton SJ, Greene EC. 2019. Defining the influence of Rad51 and Dmc1 lineage-specific amino acids on genetic recombination. Genes Dev. 33:1191–1207.

94. Storlazzi A, Gargano S, Ruprich-Robert G, Falque M, David M, Kleckner N, Zickler D. 2010. Recombination proteins mediate meiotic spatial chromosome organization and pairing. Cell. 141:94–106.

95. Sym M, Engebrecht J, Roeder GS. 1993. ZIP1 is a synaptonemal complex protein required for meiotic chromosome synapsis. Cell. 72:365–378.

96. Sym M, Roeder GS. 1994. Crossover interference is abolished in the absence of a synaptonemal complex protein. Cell. 79:283–292.

97. Szostak JW, Orr-Weaver TL, Rothstein RJ, Stahl FW. 1983. The double-strand-break repair model for recombination. Cell. 33:25–35.

98. Thacker D, Lam I, Knop M, Keeney S. 2011. Exploiting spore-autonomous fluorescent protein expression to quantify meiotic chromosome behaviors in *Saccharomyces cerevisiae*. Genetics. 189:423–439.

99. Tran PT, Fey JP, Erdeniz N, Gellon L, Boiteux S, Liskay RM. 2007. A mutation in EXO1 defines separable roles in DNA mismatch repair and post-replication repair. DNA Repair. 6:1572–1583.

100. Trelles-Sticken E, Dresser ME, Scherthan H. 2000. Meiotic telomere protein Ndj1p is required for meiosis-specific telomere distribution, bouquet formation and efficient homologue pairing. J Cell Biol. 151:95–106. doi: 10.1083/jcb.151.1.95.

101. Vernekar DV, Reginato G, Adam C, Ranjha L, Dingli F, Marsolier MC, Loew D, Guérois R, Llorente B, Cejka P, Borde V. 2021. The Pif1 helicase is actively inhibited during meiotic recombination which restrains gene conversion tract length. Nucleic Acids Res. 49:4522–4533. doi: 10.1093/nar/gkab232.

102. Wang TF, Kleckner N, Hunter N. 1999. Functional specificity of MutL homologs in yeast: evidence for three Mlh1-based heterocomplexes with distinct roles during meiosis in recombination and mismatch correction. Proc Natl Acad Sci U S A. 96:13914–13919. doi: 10.1073/pnas.96.24.13914.

103. Wanat JJ, Kim KP, Koszul R, Zanders S, Weiner B, Kleckner N, Alani E. 2008. Csm4, in collaboration with Ndj1, mediates telomere-led chromosome dynamics and recombination during yeast meiosis. PLoS Genet. 4:e1000188. doi: 10.1371/journal.pgen.1000188.

104. Wild P, Susperregui A, Piazza I, Dorig C, Oke A, Arter M, et al. 2019. Network Rewiring of Homologous Recombination enzymes during mitotic proliferation and meiosis. Mol Cell. 22:859–874.e4. doi:10.1016/j.molcel.2019.06.022.

105. Wilson MA, Kwon Y, Xu Y, Chung WH, Chi P, Niu H, Mayle R, Chen X, Malkova A, Sung P, Ira G. 2013. Pif1 helicase and Polδ promote recombination-coupled DNA synthesis via bubble migration. Nature. 502:393–396. doi: 10.1038/nature12585.

106. Xu J, Zhao L, Peng S, Chu H, Liang R, Tian M, Connell PP, Li G, Chen C, Wang HW. 2021. Mechanisms of distinctive mismatch tolerance between Rad51 and Dmc1 in homologous recombination. Nucleic Acids Res. 49:13135–13149. doi: 10.1093/nar/gkab1141.

107. Zakharyevich K, Ma Y, Tang S, Hwang PY-H, Boiteux S, Hunter N. 2010. Temporally and biochemically distinct activities of Exo1 during meiosis: double-strand break resection and resolution of double Holliday junctions. Mol Cell. 40:1001–1015.

108. Zakharyevich K, Tang S, Ma Y, Hunter N. 2012. Delineation of joint molecule resolution pathways in meiosis identifies a crossover-specific resolvase. Cell. 149:334–347.

109. Zickler D, Kleckner, N. 2015. Recombination, pairing, and synapsis of homologs during meiosis. Cold Spring Harb Perspect Biol. 7:a016626.

